# Guardians of the canopy: two new turtle ant species (Hymenoptera: Formicidae: *Cephalotes*), from Ecuador’s Chocó and Amazonia, revealed by morphology and DNA barcoding

**DOI:** 10.64898/2025.12.02.691873

**Authors:** Adrian Troya, Akayky Yumbay, Elian Ilguan, Alex Pazmiño-Palomino, Fernanda Salazar, David A. Donoso

## Abstract

We describe two new species of arboreal ants of the genus *Cephalotes* Latreille, 1802, from Ecuador: *Cephalotes esthelae* Ilguan & Troya, **sp. nov.** and *Cephalotes sacha* Yumbay & Troya, **sp. nov.** The former, a member of the *grandinosus*-group, was collected from lowland Chocó and Amazonian forests, while the latter, belonging to the *atratus*-group, is an inhabitant of the Amazonian region, including Yasuní National Park. Diagnoses and detailed morphological descriptions of workers and soldiers are provided, illustrated by high-resolution images. Species identification was further corroborated with mitochondrial CO1 sequences, which confirmed their distinctiveness from morphologically similar congeners.

Both species exhibit unique combinations of mesosomal and gastric characters. In addition, we provide the first description of the soldier of the rarely collected *C. dentidorsum* De Andrade, 1999. These discoveries expand the known diversity of the genus in Ecuador to 33 species, highlighting this region’s role as a hotspot for Neotropical myrmecofauna. A fully illustrated key and checklist of all *Cephalotes* species known from Ecuador is also presented. The new taxa occur in well-preserved but increasingly threatened forests, ecosystems subject to high rates of deforestation and mining activity, potentially endangering their populations. The integration of molecular and morphological evidence provides a robust framework for delimiting species such as in this morphologically complex and species-diverse genus. We highlight the need of further exploration of under-collected regions such as the Chocó and Andean–Amazonian foothills. Continued taxonomic and molecular investigations will be crucial to elucidate phylogenetic relationships, assess biogeographic barriers such as the Andes, and inform conservation strategies for Ecuador’s arboreal ant fauna.

## INTRODUCTION

*Cephalotes* Latreille, 1802, usually referred to as turtle ants due to their armor resembling a shell (Price et al., 2014), is currently comprised of 118 extant and 18 fossil, valid species distributed throughout the Neotropical realm, but reaching also the southern Nearctic: Arizona, southern Texas, Kentucky, and Florida (De Andrade & Baroni Urbani, 1999; Antweb, 2025). The countries with most species records of this genus are Brazil (64) and Colombia (45) (Antweb, 2025), and the biomes with the highest species richness are Amazonia and the Atlantic Forest (Antweb, 2025).

*Cephalotes* is polymorphic, both morphologically and behaviorally (Corn, 1980), and while species distinction based on workers is relatively straightforward, this is not necessarily the case for the other castes, perhaps because these are not as abundant in collections as are the workers. The reader may refer to recent revisions of the regional *Cephalotes* fauna of Colombia (Sandoval-Gómez & Sánchez-Restrepo, 2019), and Brazil (Oliveira et al., 2021) which provide excellent tools for the identification of these ants. In general, the workers are readily distinguished from other similar genera, for example *Procryptocerus*, its sister genus (see below), by the presence of strongly expanded frontal lobes, hiding the lateral portions of the clypeus, dorsoventrally slightly flattened bodies, and most species bearing spines or projections on the mesosoma, petiole and postpetiole (De Andrade and Baroni Urbani, 1999,Fernandez, 2003).

In regard to their ecology and behavior, all *Cephalotes* species are arboreal, and in general, are shy, relying mainly on pollen and compensating for their limited feeding area by licking wind-borne pollen from leaves (Baroni Urbani & De Andrade, 1997). The ecological role of many species of this genus is significant, particularly in seed dispersal and protection of certain host plants from herbivory (Powell, 2008). The adaptation to arboreal life possibly influenced their diversification and evolutionary success, enabling them to occupy a wide range of microhabitats across the Neotropics (Price et al., 2022).

*Cephalotes* was first proposed by Latreille (1802) based on Linnaeus (1758) *Formica atrata* (currently *Cephalotes atratus*). The genus has a long taxonomic history with numerous names being synonymized under it (see Oliveira et al., 2021 and De Andrade & Baroni Urbani, 1999, for a review). Although the monophyly of *Cephalotes* as sister to *Procryptocerus* has been demonstrated before (Moreau et al., 2006; Moreau & Bell, 2013; Ward et al., 2015), its internal classification in 24 species-groups, proposed by De Andrade & Baroni Urbani (1999), were tested by Price et al. (2014) who showed that some of these groups are not monophyletic. More recently, Price et al. (2022), based on the analysis of ultraconserved elements, proposed an internal conformation of 15 species-groups which need further taxonomic work to stabilize the internal nomenclature of the genus.

The current knowledge about the taxonomy and species distribution of *Cephalotes* in northwestern South America is mainly covered by the works of Kempf (1951), De Andrade & Baroni Urbani (1999), and Sandoval-Gómez & Sánchez-Restrepo (2019). For Ecuador, Salazar et al. (2015) provided the first list with distribution records for 20 species.

Subsequently, Pazmiño-Palomino & Troya (2022) recorded six additional species in the country. Since then, some potential new taxa have been identified from several collections in this country.

In the present contribution, based on morphology and DNA evidence, we describe two new species from this novel material. The new species belong to the *pinelii* + *grandinosus* and *atratus* species-groups, sensu Price et al. (2022), and were collected from the Ecuadorian coastal and Amazonian regions. In addition, we provide the first description of the soldier of the rarely collected *C. dentidorsum* De Andrade, 1999. We also provide an illustrated identification key for all 33 species known to occur in Ecuador, together with a list of all the Ecuadorian species-records we have identified in the genus and recognize that have been collected and/or observed in this territory.

## MATERIALS AND METHODS

### Descriptions and material examined

We employed DELTA (Dallwitz 1980), 1.02 version computer package. A system that allows the translation of taxonomic descriptions, from coding to natural language (Rich Text Format – RTF), which requires subsequent editing by the user. We used Microsoft Excel for converting morphometric data into RTF. The morphometric measurements were organized in a spreadsheet, which also included the collection information, as presented in the “Material examined” section.

The format of the type material follows: # specimens examined and caste, country, first administrative region, second administrative region or, if known, specific locality, coordinates in decimal degrees, elevation in meters, method of collection (if known), habitat (if known), collection date in format dd-mm-yyyy, collector followed by the word “leg.”, and institution where the material is located, usually represented by a unique specimen identifier composed of capitalized letters and numbers.

We used De Andrade & Baroni (1999), Sandoval-Gómez & Sánchez-Restrepo (2019), Oliveira et al. (2021) as guides for species-level determination. In addition, A. Troya revised previously identified material of the genus in the below collections. A. Yumbay and E. Ilguan revised previously identified material of the genus in MEPNINV. Most museum abbreviations are based on Arnett et al. (2019).

IAVH Instituto de Investigación de Recursos Biológicos Alexander von Humboldt, Villa de Leyva, Colombia.

ICN: Instituto de Ciencias Naturales, Universidad Nacional de Colombia, Bogotá, Colombia.
DZUP: Coleção Entomológica, Padre Jesus Santiago Moure, Universidade Federal do Paraná, Curitiba, Brazil.
CISEC: Colección de Invertebrados del Sur del Ecuador, Universidad Técnica Particular de Loja, Loja, Ecuador
CPDC: Coleção de Formicidae do Centro de Pesquisas do Cacau, Ilhéus, Bahia, Brazil.
MECN: Colección entomológica, Instituto Nacional de Biodiversidad, Quito, Ecuador
MEPN: Colección de insectos del Laboratorio de Invertebrados, Departamento de Biología, Escuela Politécnica Nacional, Quito, Ecuador
MUSENUV: Colección del Museo de Entomología, Universidad del Valle, Cali, Colombia.
QCAZ: Museo de Invertebrados, Pontificia Universidad Católica del Ecuador, Quito, Ecuador.
UNMSM: Colección de Insectos, Universidad Nacional Mayor de San Marcos, Lima, Peru.

In addition, we used images of type material available on Antweb ((Antweb.org). This material is deposited in the following museums.

ALWC: Alex L. Wild Collection, TX, USA
CASC: California Academy of Sciences, San Francisco, CA, USA.
JTLC: Jhon T. Longino Collection, Salt Lake City, UT, USA.
PSWC: Philip S. Ward Collection, University of California, Davis, CA, USA
MIZA: Museo del Instituto de Zoología Agrícola, Universidad Central de Venezuela, Maracay, Venezuela
MHNG: Museum of Natural History, Geneva, Switzerland.
MSNG: Museo Civico di Storia Naturale di Genova "Giacomo Doria", Genova, Italy
NHMB: Natural History Museum, Basel, Switzerland.
NHMUK: Natural History Museum, South Kensington, London, UK.
UCDC: University of California, Davis, USA.
USNM: Smithsonian National Museum of Natural History, Washington D.C, USA.

The holotypes and a part of the paratypes are in MEPN. The other paratypes, whose information is shown in the corresponding type section of each species, will be sent to the following institutions: Museo de Invertebrados, Pontificia Universidad Católica del Ecuador (QCAZ), Quito, Ecuador; Colección de Invertebrados del Sur del Ecuador, Universidad Técnica Particular de Loja (CISEC), Loja, Ecuador; Coleção Entomológica Padre Jesus Santiago Moure, Universidade Federal do Paraná (DZUP), Paraná, Brazil; Instituto Alexander von Humboldt (IAVH), Villa de Leyva, Colombia; Instituto de Ciencias Naturales, Universidad Nacional de Colombia (ICN), Bogotá, Colombia; Museum of Comparative Zoology, Harvard University (MCZC), Massachusetts, USA; United States National Museum of Natural History (USNM), Washington D.C., USA.

The material of the other examined *Cephalotes* species, i.e., those we previously identified as valid species-records for Ecuador (Supplementary file 1), are in the following collections (in order of number of specimen-records); Museo de Invertebrados, Pontificia Universidad Católica del Ecuador (QCAZ); Laboratorio de Invertebrados, Escuela Politécnica Nacional; Instituto Nacional de Biodiversidad (MECN); Colección de Invertebrados del Sur del Ecuador, Universidad Técnica Particular de Loja (CISEC); Museo de la Universidad de Guayaquil (MUGT).

### Imaging

We used a Canon® EOS DSLR 70D camera equipped with a Laowa® 25mm F2.8, 2.5X - 5.0X macro lens, coupled to a Cognisys StackShot® macro rail system with a stepper motor controlled by Helicon Remote software v. 3.9.9. The resulting images were stacked with Helicon Focus v. 7.5.6. We used a dome equipped with LED cold lights for illumination. The images of the holotypes and selected paratypes, for example, soldiers, and their respective labels, are available on AntWeb.org.

### Measurements and indices

The measurements of the below morphological variables were taken using an ocular micrometer integrated into a KRÜSS MSZ5000-T-IL-TL stereo microscope with magnifications ranging from 0.7X to 4.5X. To facilitate visual examination, we used a pin–holding stage that enables rotations along the X, Y, and Z axes. Measurements are given in mm, with an accuracy of two decimals. Minimum and maximum values are provided for each morphometric variable. All measurements are in line with those of Oliveira et al. (2021). A schematic illustration of such variables in represented in Fig. 1.

**Figure 1.**
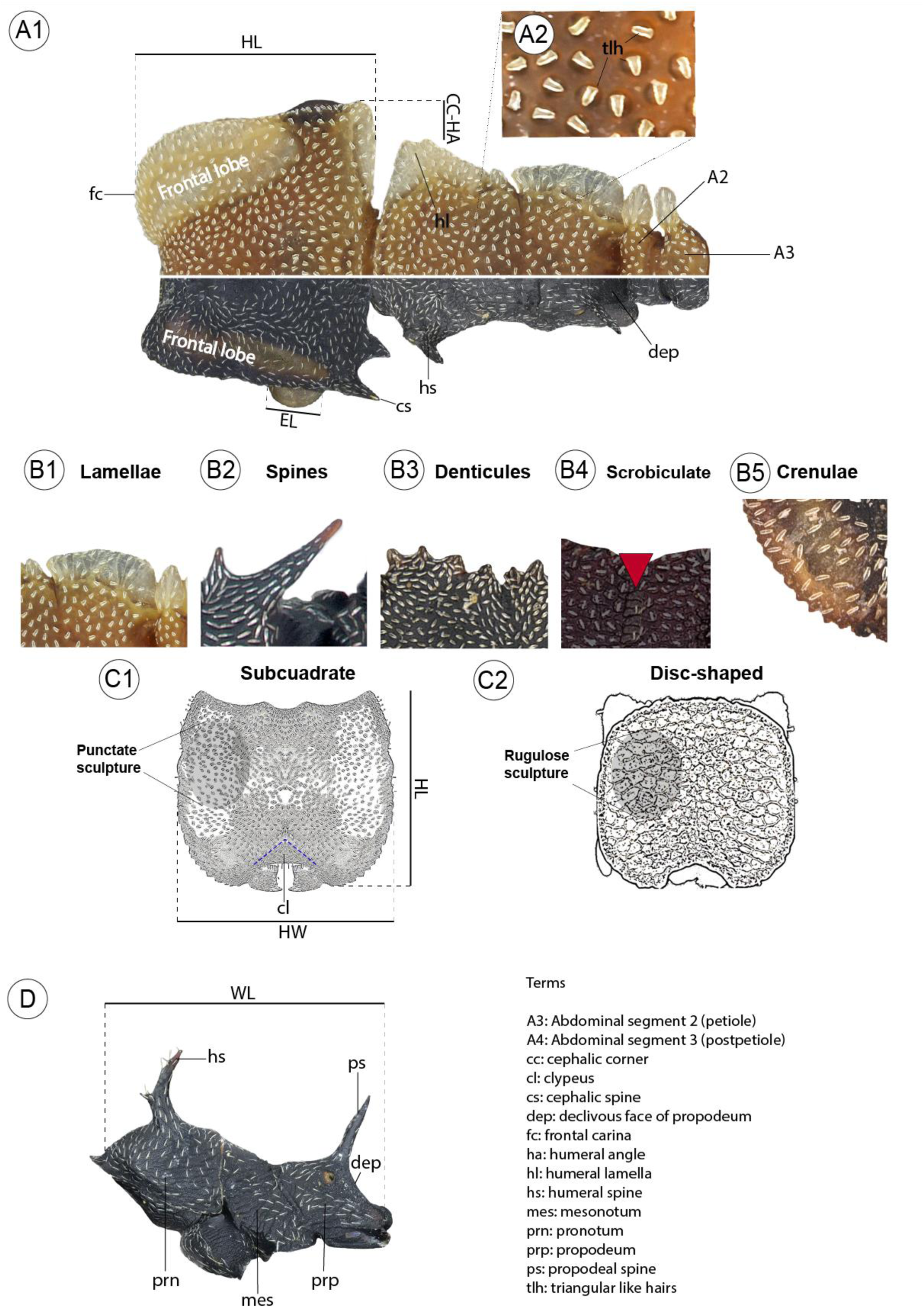
Common morphology structures and measurements cited in text. A1. Body, except gaster, dorsal view. A2. Close-up of body hairs. B1 – B5. Sculpture. C1 & C2. Head shapes. D. Mesosoma, lateral view. Design by Elian Ilguan.

HW: Head width. In full-face view, maximum width of cranium. In the workers, this is measured posterior to the eyes, including spines and/or lamellae.
HL: Head length. In full-face view, maximum distance from the posterior margin of head, including spines and/or lamellae, to the anterior margin of frontal lobe.
EL: Eye length. In lateral view, the maximum measurable length of eye.
PW: Pronotum width. In dorsal view, maximum width of pronotum, including pronotal projections and/or lamellae.
WL: Weber’s length. In lateral view, the diagonal length of mesosoma, from the anterior pronotal margin to the posterior margin of metapleura.
PTW: Petiolar width. In dorsal view, maximum width of petiolar node, including petiolar projections and/or lamellae.
PTL: Petiolar length. In dorsal view, the distance between the anterior most margin of petiole to its posterior margin, excluding the posteroventral ridge.
PPW: Postpetiolar width. In dorsal view, maximum width of postpetiole, including the postpetiolar projections and/or lamellae.
PPL: Postpetiolar length. In dorsal view, distance between the anterior most margin of postpetiole to its posterior margin, excluding the posteroventral ridge.
GW: Gaster width. In dorsal view, maximum width of the first gastral tergite
GL: Gaster length. In dorsal view, maximum length of the first gastral tergite, including the anterior lamellar margins.
HBW: Hind basitarsus width. Maximum width of the hind basitarsus.
HBL: Hind basitarsus length. Maximum length of the hind basitarsus.
CCHA: Cephalic corner to humeral angle. In dorsal view, horizontal distance between the posterolateral cephalic margin to the lateral humeral margin.
ID: Interspinal distance. In frontal view, distance between the apices of pronotal spines.
TL: Total length. The summed length of HL + WL + PTL + PPL + GL.

The following ratios were multiplied by 100:

CI: Cephalic index: HW/HL
OI: Ocular index: EL/HW
HBI: Hind basitarsal index. HBW/HBL.
PI: Petiolar index. PTL/PTW

### DNA sequencing and phylogenetic reconstruction

We used the subunit one (CO1) of the cytochrome c oxidase mitochondrial gene complex to infer the possible phylogenetic placement of our new species in the *Cephalotes* CO1 tree, but also their placement within their respective species-groups sensu Price et al. (2022): *atratus*-group for *C. sacha* sp. nov., and *pinelii* + *grandinosus*-group for *C. esthelae* sp. nov.

Except for *C. esthelae*, the sequences of all species, including the outgroups, were downloaded from Bold Systems, version 4 (boldsystems.org). Of the 2300 available sequences we selected only 295 (Supplementary file 2) based on the following criteria: sequences without contamination and stop codons; sequences with more than 300 base pairs; in case of species or morphospecies with numerous sequenced specimens from a single site, we selected only two representatives per site, per country; sequences lacking voucher images were not selected unless these were linked to previously published studies related to the systematics of *Cephalotes*, e.g., Price et al. (2014), Oliveira et al. (2021), Price et al. (2022); and finally, we did not select sequences of species clearly misplaced in the topology when performing prior phylogenetic tests. Of the 295 sequences, 87 were generated for this study.

Using two legs per specimen, genomic DNA of *C. esthelae* sp. nov. was extracted by the lead author at the Zoologische Staatssammlung München, Germany. Based on the Puregene Core Kit A (Qiagen Sciences, USA), the gene was amplified following a two-step PCR: first step - 1 min/94 °C; 5 cycles (30 sec/94 °C; 40 sec/47 °C; 10 min/72 °C); second step - 30 sec /94 °C; 30 cycles (40 sec/52°C; 1 min /72 °C; 10 min/72 °C). Forward primer LCO1490: 5’-GGT CAACAA ATC ATA AAG ATA TTGG-3’ and reverse HC02198: 5’-TAA ACT TCA GGG TGACCA AAA AAT CA-3’ from Folmer et al. (1994). The amplicons were sent to the University of Munich’s Biozentrum facility for sequencing.

The sequences were aligned using MAFFT version 7 (Katoh and Standley, 2013) and the resulting block was visually inspected for unaligned sites which were removed. The length of the final block consisted of 654 base pairs.

We built three trees based on the following sets of sequences: 1. ‘*Cephalotes*_total’ represented by the whole set of 295 sequences (Fig. 42); 2. ‘*Cephalotes*_*atratus*-group’ represented by the available sequenced species in the *atratus* species-group (Fig. 2); and 3. ‘*Cephalotes*_*pinelii*-*grandinosus*-group’ represented by the available sequenced species in the *pinelii* + *grandinosus* species-group (Fig. 3). The aligned sequences were submitted to the Cyberinfrastructure for Phylogenetic Research - CIPRES (Miller et al., 2010), and the dataset was analyzed with IQ-TREE (Nguyen et al., 2015, Minh et al., 2020), a maximum likelihood-based tree inference package, using the following parameters: model of molecular evolution TIM2+F+I+G4, based in the Bayesian information criterium – BIC, as implemented in ModelFinder (Kalyaanamoorthy et al., 2017); the node supports were calculated using 1000 rounds of ultrafast bootstrap (Hoang et al., 2018) replicates (as recommended in Nguyen et al., 2015), together with the Shimodaira–Hasegawa approximate likelihood ratio test- SH-aLRT, and the Bayesian-like transformation of the aLRT – aBayes (Anisimova et al., 2011) branch support methods. The trees were rooted with species of *Pheidole* (*P. zeteki* Smith, *P. biconstricta* Mayr) and *Procryptocerus* (*P. adlerzi* Mayr*, P. regularis* Emery), which according to Ward et al. (2015) are the closest lineages to *Cephalotes*. The topologies were visualized and edited in FigTree (Rambaut, 2018) and Adobe Illustrator **®**.

**Figure 2.**
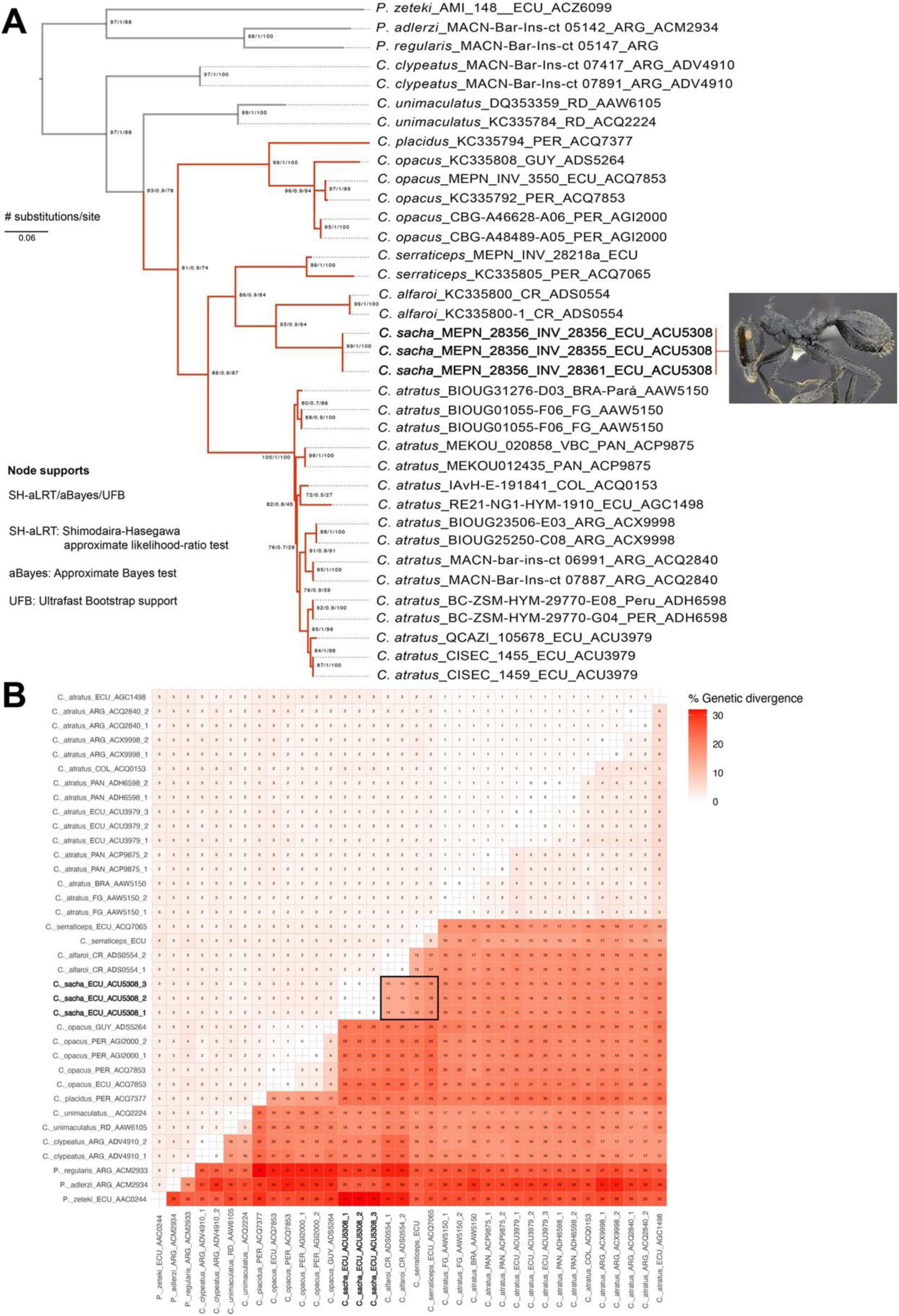
A. Maximum likelihood CO1 phylogram of the *Cephalotes atratus* species-group (red topology). Gray branches are outgroups. The tree tips contain: species name, collection code, country, BIN (Barcode Index Number). Newly described species are bolded. **B.** Genetic divergence. Below the diagonal: estimates (in %) of evolutionary divergence between sequences; above the diagonal: standard error estimates. The genetic divergence between *C. sacha* and its closest species (*C. alfaroi* and *C. serraticeps*) is denoted inside a black square.

**Figure 3.**
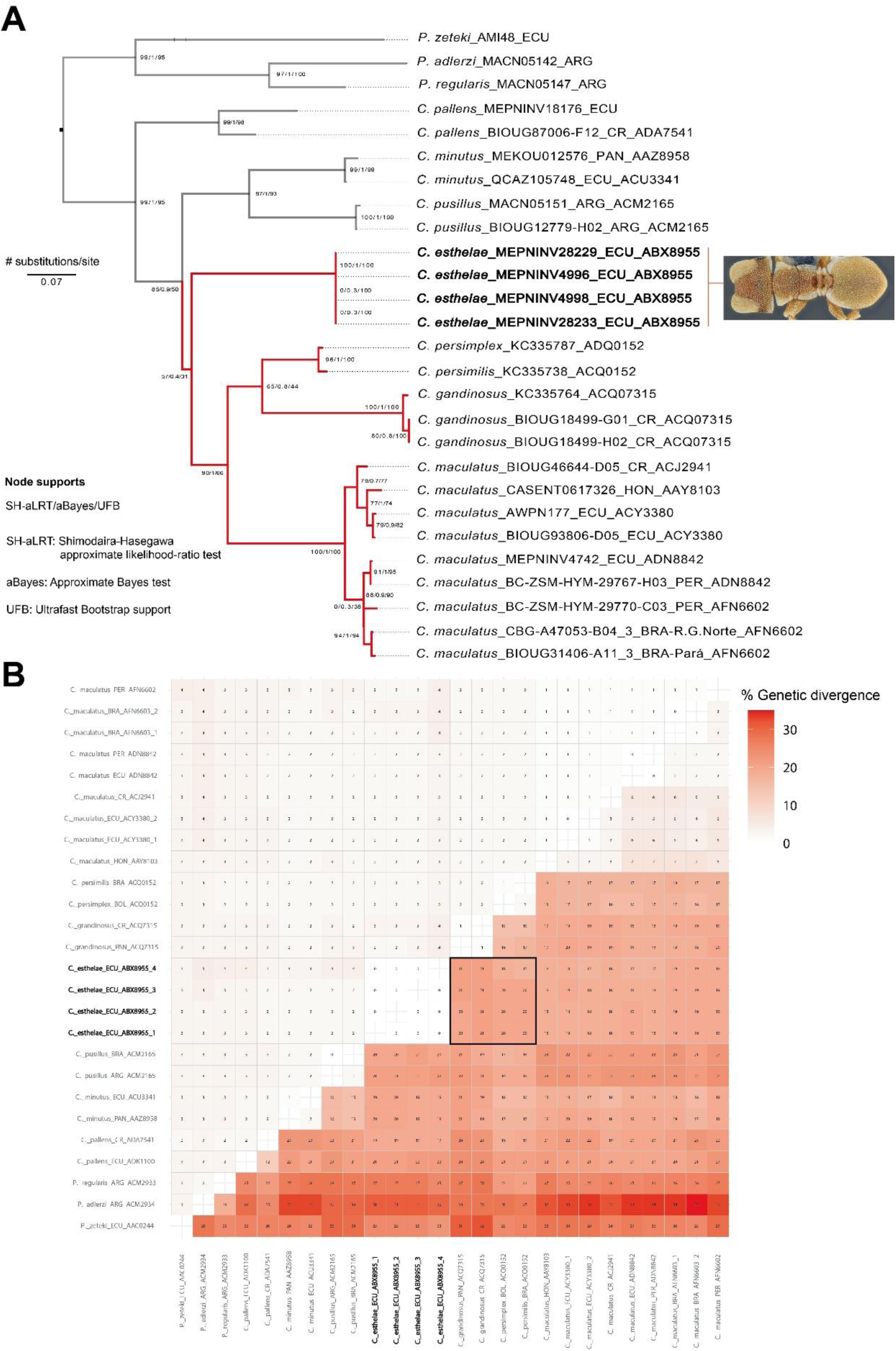
A. Maximum likelihood CO1 phylogram of the *Cephalotes grandinosus+pinelii* species-group (red topology). Gray branches are outgroups. The tree tips contain: species name, collection code, country, BIN (Barcode Index Number). Newly described species are bolded. **B.** Genetic divergence. Below the diagonal: estimates (in %) of evolutionary divergence between sequences; above the diagonal: standard error estimates. The genetic divergence between *C. grandinosus* and its closest species (*C. grandinosus*, *C. persimilis*, *C. persimplex*) is denoted inside a black square.

### Distribution of examined records

We used ArcGIS online (esri.com) to generate maps of distribution of the examined records based on coordinates written on specimen labels as well as on our own personal field notes. When coordinates were not available, we used Google Earth (Google 2025) to obtain them based on specific localities below the circumscription of province. The coordinates and their associated geographic collection data are shown in Supplementary file 1.

### Morphology terms

General morphological terminology follows mainly De Andrade & Baroni (1999), Oliveira et al. (2021) and Fernandez et al. (2019). Surface sculpture is based on Harris (1979). Pilosity follows Serna and MacKay (2010) for appressed, erect and suberect hairs. Some common morphology terms cited in the text are depicted in Fig. 1.

### Species concept

We follow Mayr’s (1942) concept, which defines a species as a group of interbreeding natural populations which are reproductively isolated from other such groups. Since we cannot demonstrate this process by direct evidence, we interpret our examined material as being members of new species due to their high intraspecific morphological similarity, expressed in unique sets of external traits differentiating them from all other known closely related species, thus suggesting reproductive isolation.

### Nomenclatural acts

The electronic version of this article in Portable Document Format (PDF) will represent a published work according to the International Commission on Zoological Nomenclature (ICZN), and hence the new names contained in the electronic version are effectively published under that Code from the electronic edition alone. This published work and the nomenclatural acts it contains have been registered in ZooBank, the online registration system for the ICZN. The ZooBank LSIDs (Life Science Identifiers) can be resolved and the associated information viewed through any standard web browser by appending the LSID to the prefix http://zoobank.org/. The LSID for this publication is: urn:XXX [insert # upon publishing]. The online version of this work is archived and available from the following digital repositories: PeerJ, PubMed Central and CLOCKSS.

## RESULTS

### CO1 Cephalotes tree and molecular discrimination of the new species

Our CO1 *Cephalotes* tree contains 58 valid species and ten morphospecies (Fig. 42). All species-groups proposed in Price et al. (2022) were represented in the topology, but as expected, the arrangement of species-groups with respect to Price et al. (2022), differed, for example, in our phylogeny the *atratus* group is not at the base of the tree and is sister to the *basalis* group, while in Price et al. (2022) the *atratus* group is sister to all the other species-groups (Fig. 1 of their phylogeny). As to the placement of species within each species-group we noted minor variations with respect to Price et al. (2022). In general, within-group species relationships are relatively similar in both our analysis and theirs. Price et al. (2022) however, include in their work 24 species and various other morphospecies which we do not include here because there are no available CO1 sequences for them yet. Possibly, the most notorious difference in within-group species relationships between our study and that of Price et al. (2022), is in the *pallens* group, where the included species are split in two separate groups: the first consisting of *C. pallens*, *C. porrasi*, *C. patellaris*, and two unidentified species from Belize and Costa Rica, and a second group consisting of *C. pallidus* and an unidentified, possibly new species, *C.* YASsp1, from French Guiana (Fig. 42). The first *pallens* group comes out as sister to the new species *C. esthelae*, while the second *pallens* group comes out as sister to *C. umbraculatus*, a monotypic species-group.

The Amazonian new species *C. sacha*, which belongs to the *atratus* species-group, comes out as sister to the Central American *C. alfaroi* in our phylogeny, both are genetically closer to *C. serraticeps*, the node supporting this relationship is relatively high (Fig. 2A). *Cephalotes alfaroi* shows about 14% of genetic divergence from *C. sacha*, as measured by the number of substitutions per site (Fig. 2B). *Cephalotes serraticeps*, on the other hand, which is also morphologically similar to *C. sacha*, has about 16% genetic divergence with it. These three species are closer to *C. atratus* than they are to *C. opacus* and *C. placidus*. *Cephalotes oculatus* is the only member of the atratus group not represented in the atratus species-group.

*Cephalotes esthelae*, a member of the *grandinosus* species-group, is morphologically most similar to *C. persimilis* and *C. persimplex* (see further details under its respective treatment), but as said before, it does not come out as sister, not even closer to them in the *Cephalotes* CO1 tree (Fig. 42). *Cephalotes esthelae* shows about 17 – 20 % of genetic divergence from *C. persimilis* and *C. persimplex*, and about 20% from *C. grandinosus* (Fig. 3B), which is also morphologically similar to the latter. Although these four species are overall similar there is a clear genetic divergence of *C. esthelae* as compared to them, the node supporting this relationship is weak (Fig. 3A).

### Taxonomic account

*Cephalotes species known to occur in Ecuador. New country records with \**

Cephalotes atratus (Linnaeus, 1758)
Cephalotes basalis (Smith, 1876)
Cephalotes cordatus (Smith, 1853)
Cephalotes dentidorsum De Andrade, 1999
Cephalotes depressus (Klug, 1824)
Cephalotes ecuadorialis De Andrade, 1999
Cephalotes esthelae **sp. nov.**
Cephalotes goeldii (Forel, 1912)
Cephalotes grandinosus (Smith, 1860)
Cephalotes inca (Santschi, 1911)
Cephalotes laminatus (Smith, 1860)
Cephalotes maculatus (Smith, 1876)
Cephalotes manni (Kempf, 1951)
Cephalotes minutus (Fabricius, 1804)
Cephalotes opacus Santschi, 1920
Cephalotes pallens (Klug, 1824)
Cephalotes pallidus De Andrade, 1999
Cephalotes patellaris (Mayr, 1866)*
Cephalotes pavonii (Latreille, 1809)
Cephalotes peruviensis De Andrade, 1999
Cephalotes porrasi (Wheeler, 1942)
Cephalotes pusillus (Klug, 1824)
Cephalotes ramiphilus (Forel, 1904)
Cephalotes sacha **sp. nov.**
Cephalotes scutulatus (Smith, 1867)
Cephalotes serraticeps (Smith, 1858)
Cephalotes setulifer (Emery, 1894)
Cephalotes simillimus (Kempf, 1951)
Cephalotes solidus (Kempf, 1974)
Cephalotes spinosus (Mayr, 1862)
Cephalotes targionii (Emery, 1894)*
Cephalotes trichophorus De Andrade, 1999
Cephalotes umbraculatus (Fabricius, 1804)

*Worker-based key to Cephalotes species known to occur in Ecuador*.

Note. Next to each species name the species-group (sensu Price et al., 2022) is shown in parentheses.

1. In full-face view, frontal carina with ocular incision (Fig. 4A) **2**

- In full-face view, frontal carina without ocular incision (Fig. 4B) **5**
2. **2(1).** Head surface rugulose ventrally (Fig. 5A) ***C. patellaris* (Mayr) (*pallens* group)**

- Head surface punctate ventrally (Fig. 5B) **3**
3. **3(2).** In full-face view, frontal lobe with longitudinal striae clearly visible (Fig. 6A); lateral propodeal margin continuous (without teeth), crenulate and membranaceous (Fig. 6C) . ***C. pallidus* De Andrade (*pallens* group)**

- In full-face view, frontal lobe with longitudinal striae barely visible (Fig. 6B); lateral propodeal margin with two or three distinctive membranaceous denticles (Fig. 6D) . **4**
4. **4(3).** In dorsal view, scrobiculate propodeal groove well-marked laterally, absent medially (Fig. 7A) ***C. porrasi* (Wheeler) (*pallens* group)**

- In dorsal view, scrobiculate propodeal groove well-marked along the entire surface (Fig. 7B) ***C. pallens* (Klug) (*pallens* group)**
5. **5(1).** In full-face view, posterolateral margin of head with squared (Fig. 8A) or sharply bicrested (Fig. 8B-C) lamella, either opaque or subhyaline, never with spines … **16**

- In full-face view, posterolateral margin of head either: continuous or feebly crested (Fig. 8D), sharply crested (Fig. 8E), or with spines (Fig. 8F), never with opaque or subhyaline lamella **6**
6. **6(5).** In full-face view, posterolateral margin of head with two spines (Fig. 9A) . **7**

- In full-face view, posterolateral margin of head without spines, although it can be angulate or carinate (Fig. 9B) **10**
7. **7(6).** In lateral view, eye placed behind scrobe (Fig. 10A) ………………………………… ***C. opacus* Santschi (*atratus* group)**

- In lateral view, eye placed under scrobe (Fig. 10B) **8**
8. **8(7).** In dorsal view, pronotum with inverted Y-shaped carina placed anterior to promesonotal groove (Fig. 11A) ***C. serraticeps* (Smith) (*atratus* group)**

- In dorsal view, pronotum without inverted Y-shaped carina (Fig. 11B) **9**
9. **9(8).** Integument opaque; in lateral view, subpostpetiolar process shorter than postpetiolar dorsal spines (Fig. 12A) ***C. sacha* sp. nov. (*atratus* group)**

- Integument shiny; in lateral view, subpostpetiolar process longer than postpetiolar dorsal denticles (Fig. 12B) ***C. atratus* (*atratus* group)**
10. **10(6).** In full-face view, frontal carina crenulate (Fig. 13A) **11**

In full-face view, frontal carina without crenulae, smooth (Fig. 13B) **12**
11. **11(10).** In dorsal view, humerus with narrow, acute lamella (Fig. 14A); gaster relatively shiny (Fig. 14C); in lateral view, subpostpetiolar process elongated (Fig. 14E) … ***C. trichophorus* De Andrade (*coffeae* group)**

- In dorsal view, humerus denticulate (Fig. 14B); gaster opaque (Fig. 14D); in lateral view, subpostpetiolar process short (Fig. 14F).. ***C. setulifer* (Emery) (*coffeae* group)**
12. **12(10).** In dorsal view, mesonotum and propodeum with denticles (Fig. 15A) or spines (Fig. 15B) **13**

- In dorsal view, mesonotum and propodeum without projections (Fig. 15C) . ***C. solidus* (Kempf) (*solidus* group)**
13. **13(12).** In dorsal view, pronotum and propodeum tridenticulate (Fig. 16A) … ***C. manni (Kempf)* (*manni* group)**

- In dorsal view, pronotum with lamella, either angulate or rounded anteriorly and posteriorly; propodeum with two spines (Fig. 16B) **14**
14. **14(13).** In dorsal view, humerus subacute to rounded anteriorly, and rounded posteriorly (Fig. 17A) ***C. ramiphilus* (Forel) (*basalis* group)**

- In dorsal view, humerus with spine anteriorly, and acute posteriorly (Fig. 17B) …**15**
15. **15(14).** In dorsal view, petiolar spine much longer than postpetiolar spine: distance from petiolar spinal tip to postpetiolar spinal tip about as long as mid postpetiolar length (Fig. 18A) ***C. inca* (Santschi) (*basalis* group)**

- In dorsal view, petiolar spine subequal to- or at most slightly longer than postpetiolar spine: distance from petiolar spinal tip to postpetiolar spinal tip clearly shorter than mid postpetiolar length (Fig. 18B) ***C. basalis* (Smith) (*basalis* group)**
16. **16(5).** In dorsal view, propodeum with single, broad lamella (Fig. 19A) **17**

In dorsal view, propodeum with one or more denticles (Fig. 19B) or spines (Fig. 19C) . **20**
17. **17(16).** In dorsal view, pronotal lamella truncate or with incision anterolaterally (Fig. 20A) . **18**

- In dorsal view, pronotal lamella continuous, without incision anterolaterally (Fig. 20B) . **19**
18. **18(17).** In dorsal view, mesonotum with triangular, subhyaline lamella; petiole and postpetiole with subhyaline, rhomboid-shaped lamella (Fig. 21A) … ***C. esthelae* sp. nov. (*pinelii* + *grandinosus* group)**

- In dorsal view, mesonotum with denticle; petiole and postpetiole with spines (Fig. 21B) ***C. scutulatus* (Smith) (*texanus* + *bimaculatus* + *pinelii* group)**
19. **19(17).** In dorsal view, mesonotum with two lamellae, the anterior is bigger than the posterior (Fig. 22A); hind femur with crenulate lamella at anterior and posterior margins (Fig. 22C) … ***C. grandinosus* (Smith) (*pinelii* + *grandinosus* group)**

- In dorsal view, mesonotum with single denticle or lamella (Fig 22B); hind femur without crenulate lamellae (Fig. 22D) ………………………………………….. … ***C. maculatus* (Smith) (*pinelii* + *grandinosus* group)**
20. **20(16).** In dorsal view, gaster with two dark linear spots, connected medially by dark spot (Fig. 23A) ***C. umbraculatus* (Fabricius) (*umbraculatus* group)**

- In dorsal view, gaster concolored (Fig 23B) or heterocolored (Fig. 23C), never with two dark linear spots **21**
21. **21(20).** In dorsal view, propodeum with single denticle (Fig. 24A) ……………………………… ***C. ecuadorialis* De Andrade (*crenaticeps* group)**

- In dorsal view, propodeum with two or more denticles (Fig. 24B) or spines (Fig. 24C) . **22**
22. **22(21).** In dorsal view, propodeum with two spines (Fig. 25A) **23**

- In dorsal view, propodeum with denticles: the median denticle usually longer than the anterior and posterior denticles (Fig. 25B) **30**
23. **23(22).** In dorsal view, posterior propodeal spine longer than anterior spine (Fig. 26A) .. …………………………………………………………………………………………..**24**

- In dorsal view, posterior propodeal spine shorter than anterior spine (Fig. 26B) **28**
24. **24(23).** In dorsal view, mesonotum without denticle (Fig. 27A) …………………..*………………* ***C. spinosus* (Mayr) (*laminatus* + *pusillus* group)**

- In dorsal view, mesonotum with denticle or spine (Fig. 27B) **25**
25. **25(24).** In dorsal view, declivous face of propodeum with marked longitudinal striae (Fig. 28A) ***C. minutus* (Fabricius)** (***laminatus* + *pusillus* group)**

- In dorsal view, declivous face of propodeum without striae, but it can have minute puncta (Fig. 28B) **26**
26. **26(25).** Petiole with tiny denticle laterally (Fig. 29A) ……………………………………..… ***C. pusillus*** (**Klug)** (***laminatus* + *pusillus* group)**

- Petiole with distinctive spine laterally (Fig. 29B) **27**
27. **27(26).** In dorsal view, posterior margin of head and first tergite of gaster with subhyaline lamellae (Fig. 30A) ***C. laminatus* (Smith) (*laminatus* + *pusillus* group)**

- In dorsal view, posterior margin of head and first tergite of gaster without subhyaline lamellae, this is, lamellae are concolorous with rest of body or at most, with light brown shades on lamellar margins (Fig. 30B) …………………………………………………………………………………………….… ***C. simillimus* (Kempf) (*laminatus* + *pusillus* group)**
28. **28(23).** In posterior view, vertex with two denticles (Fig. 31A); pronotal lamellar margin mostly plain (Fig. 31C) ***C. cordatus* (Smith) (*depressus* group)**

- In posterior view, vertex without denticles (Fig. 31B); pronotal lamellar margin bidentate (Fig. 31D) or crenulate (Fig. 31E) **29**
29. **29(28).** Pronotum with bidenticulate lamella; a deep notch is present between lamella and posterior pronotal teeth (Fig. 32A) ***C. depressus* (Klug) (*depressus* group)**

- Pronotum with broad, crenulate lamella; a shallow notch is present between lamella and posterior pronotal teeth (Fig. 32B). ***C. pavonii* (Latreille) (*depressus* group)**
30. **30(22).** In dorsal view, propodeum with four to five well-developed denticles, with the second and fourth longer than the rest (Fig. 33A) **31**

- In dorsal view, propodeum with three or less well-developed denticles, the median is always longer than the rest (Fig. 33B) Note: Denticles not always well-visible … ***C. peruviensis* De Andrade (*coffeae* group)**
31. **31 (30).** In dorsal view, pronotum with spine anteriorly, followed by broad, crenulate lobe upturned (Fig 34A), more evident in lateral view (Fig. 34C) ……………………………………… ***C. dentidorsum* De Andrade (*angustus* + *fiebrigi* + *bruchi* group)**

- In dorsal view, pronotum with three membranaceous denticles (Fig. 34B), without upturned lobe (Fig. 34D) **32**
32. **32(31).** In dorsal view, postpetiole with wing-shaped, lateral lamella, recurved and broad distally (Fig. 35A) ….. ***C. goeldii* (Forel) (*angustus* + *fiebrigi* + *bruchi* group)**

- In dorsal view, postpetiole with thinner, lateral spine, never wing-shaped (Fig. 35B) ***C. targionii* (Emery) (*angustus* + *fiebrigi* + *bruchi* group)**

**Figure 4.**
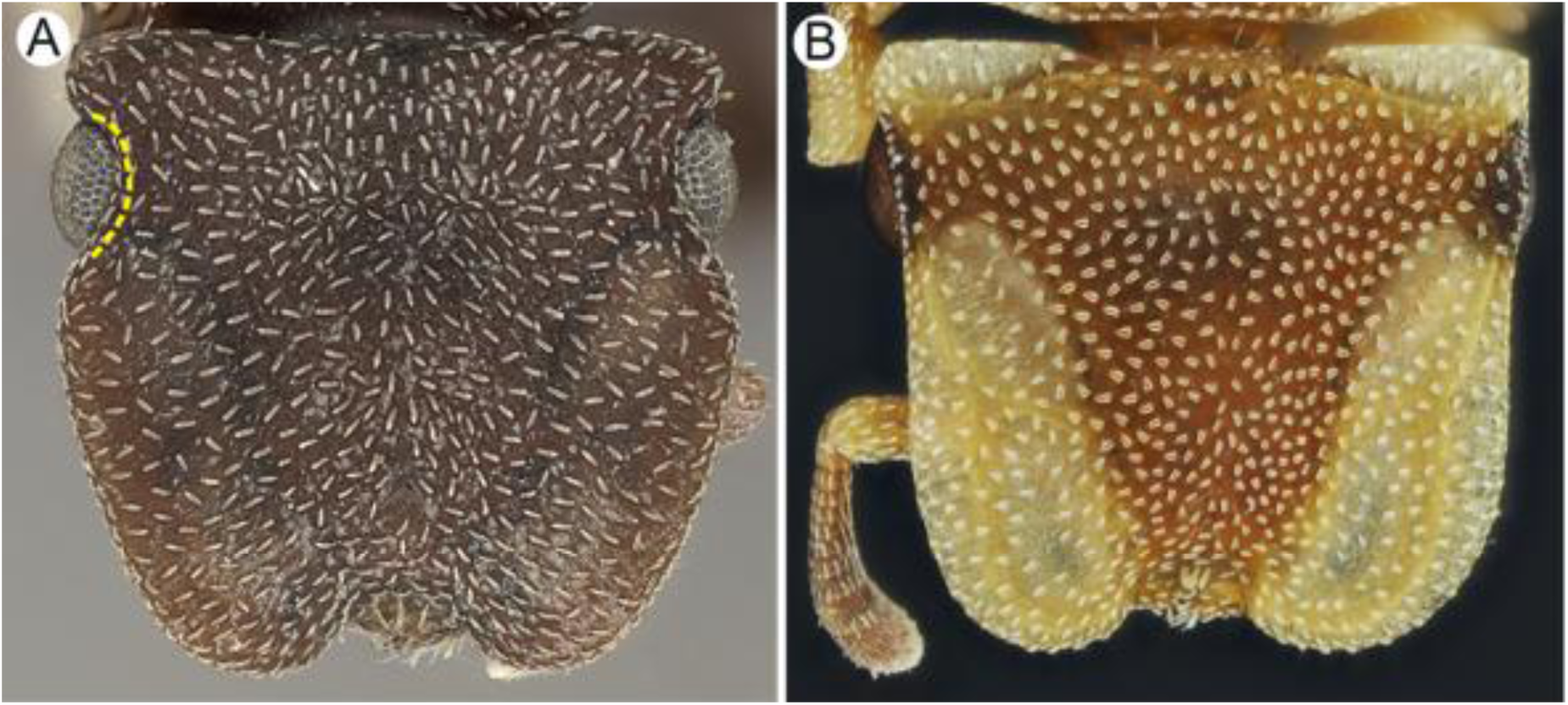
Frontal carina (full-face view): **A.** Notched (yellow dotted line). Image by Shannon Hartman. **B**. Plain. Image by Elian Ilguan.

**Figure 5.**
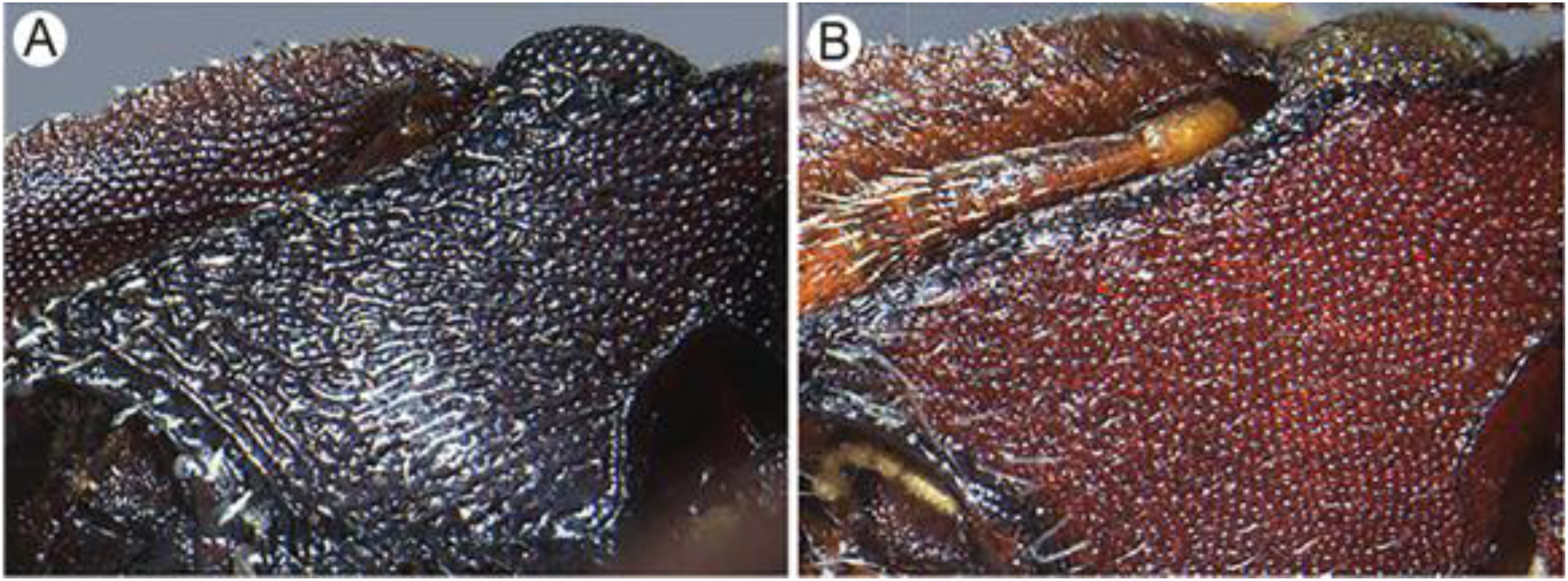
Head surface (ventral view): **A.** Rugulose. **B**. Punctate. Images from Oliveira et al. (2022), with their permission.

**Figure 6.**
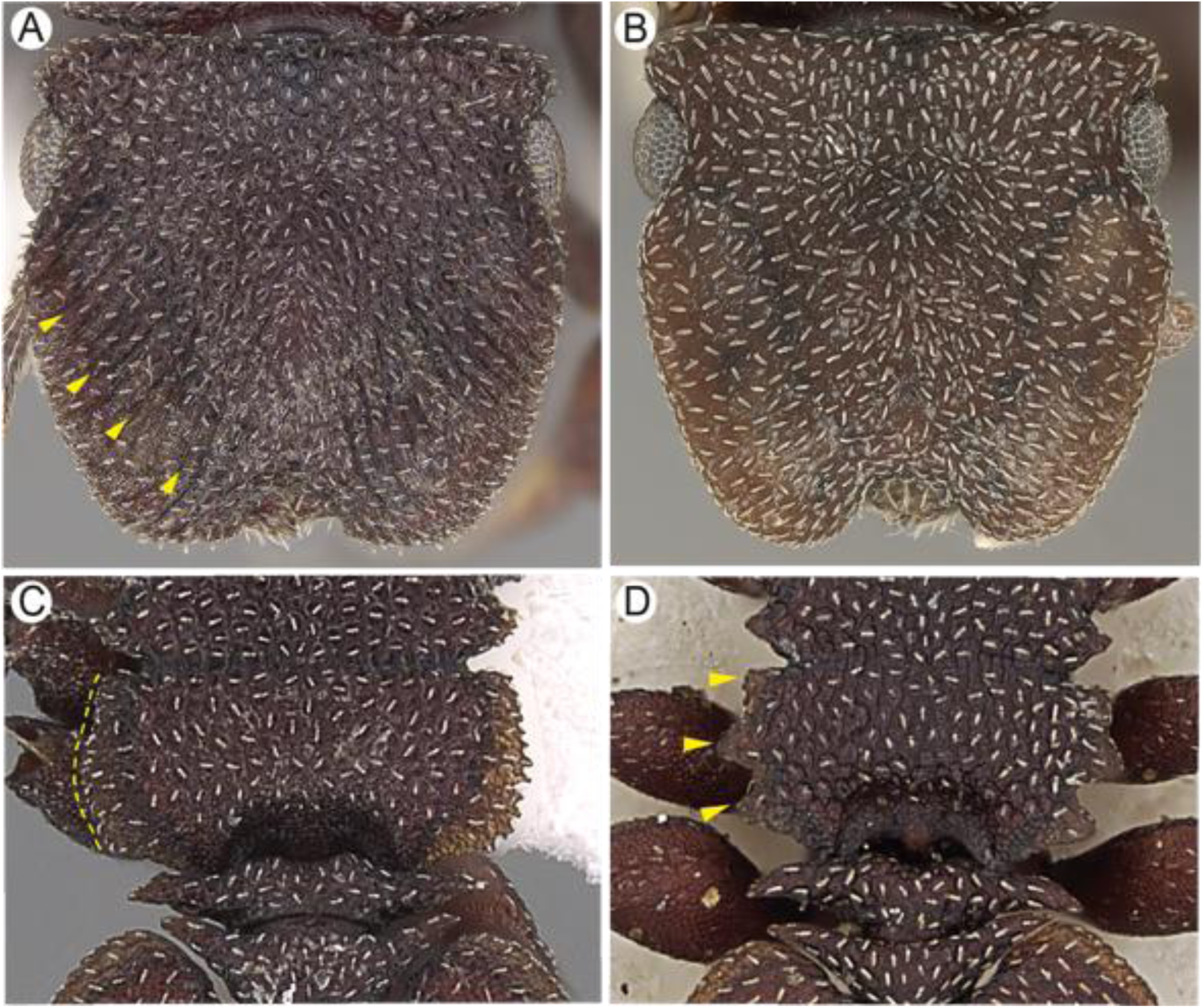
Frontal lobe (full-face view): **A.** Longitudinally striate. Image by Wade Lee. **B**. Barely striate. Image by Shannon Hartman. Lateral pronotal margin (dorsal view): **C.** Membranaceous, crenulate. Image by Wade Lee. **D.** Denticulate. Image by Ryan Perry.

**Figure 7.**
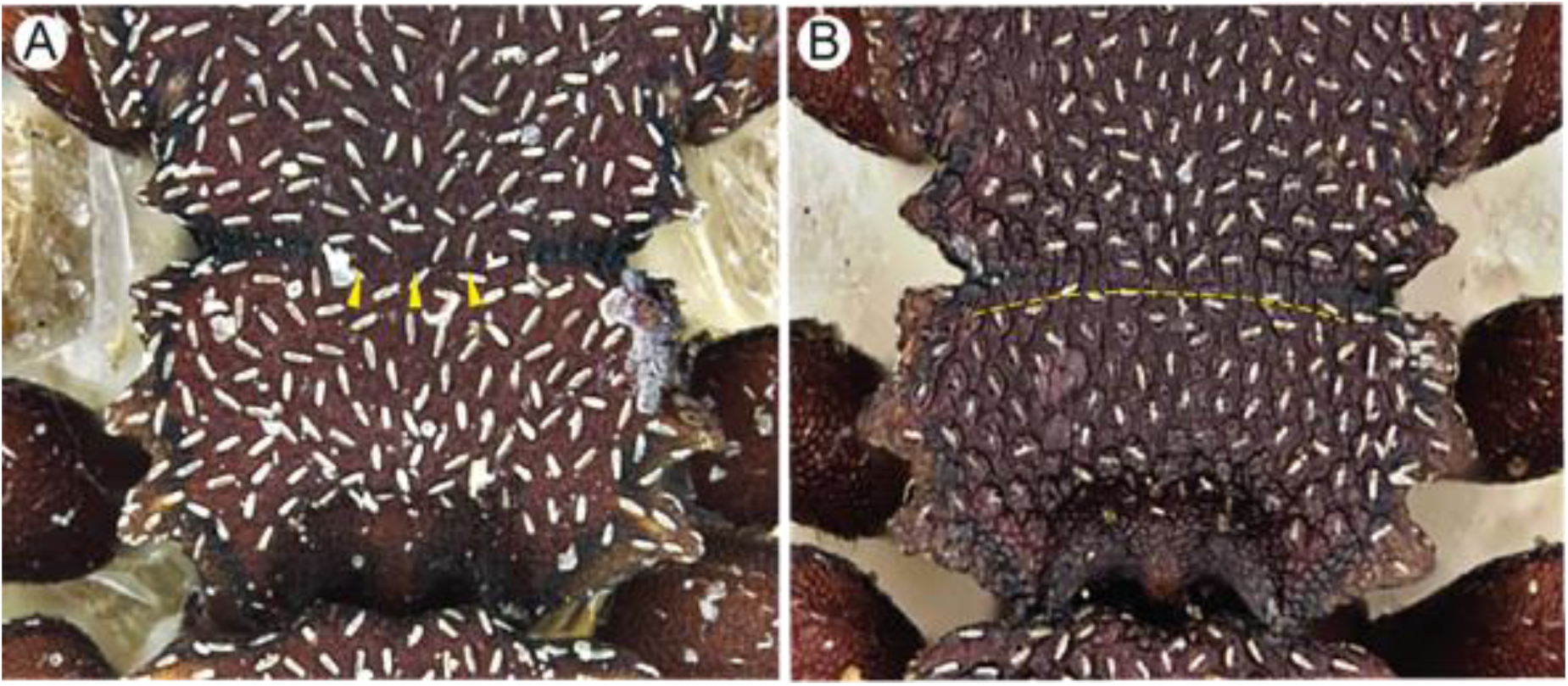
Propodeal groove (dorsal view): **A.** Absent medially (yellow arrowheads). Image by Zack Lieberman. **B.** Continuous. Image by Ryan Perry.

**Figure 8.**
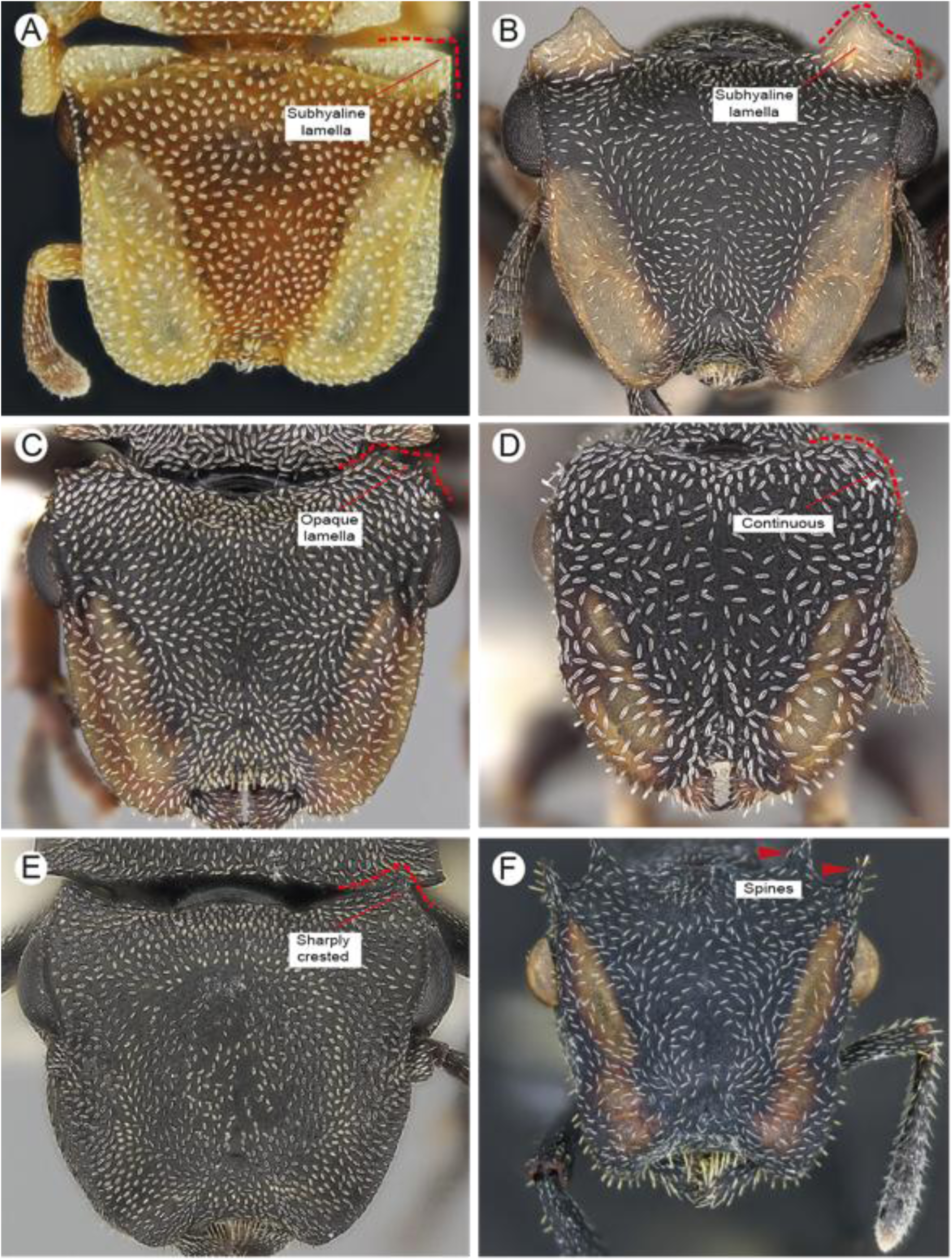
Posterolateral head margin (full-face view): **A, B.** Squared lamella. Image by Ryan Perry. **C.** Opaque lamella. Image by Wade Lee. **D.** Continuous. Image by Wade Lee. **E.** Sharply crested. Image by Shannon Hartman. **F.** Spiny. Image by Akayky Yumbay.

**Figure 9.**
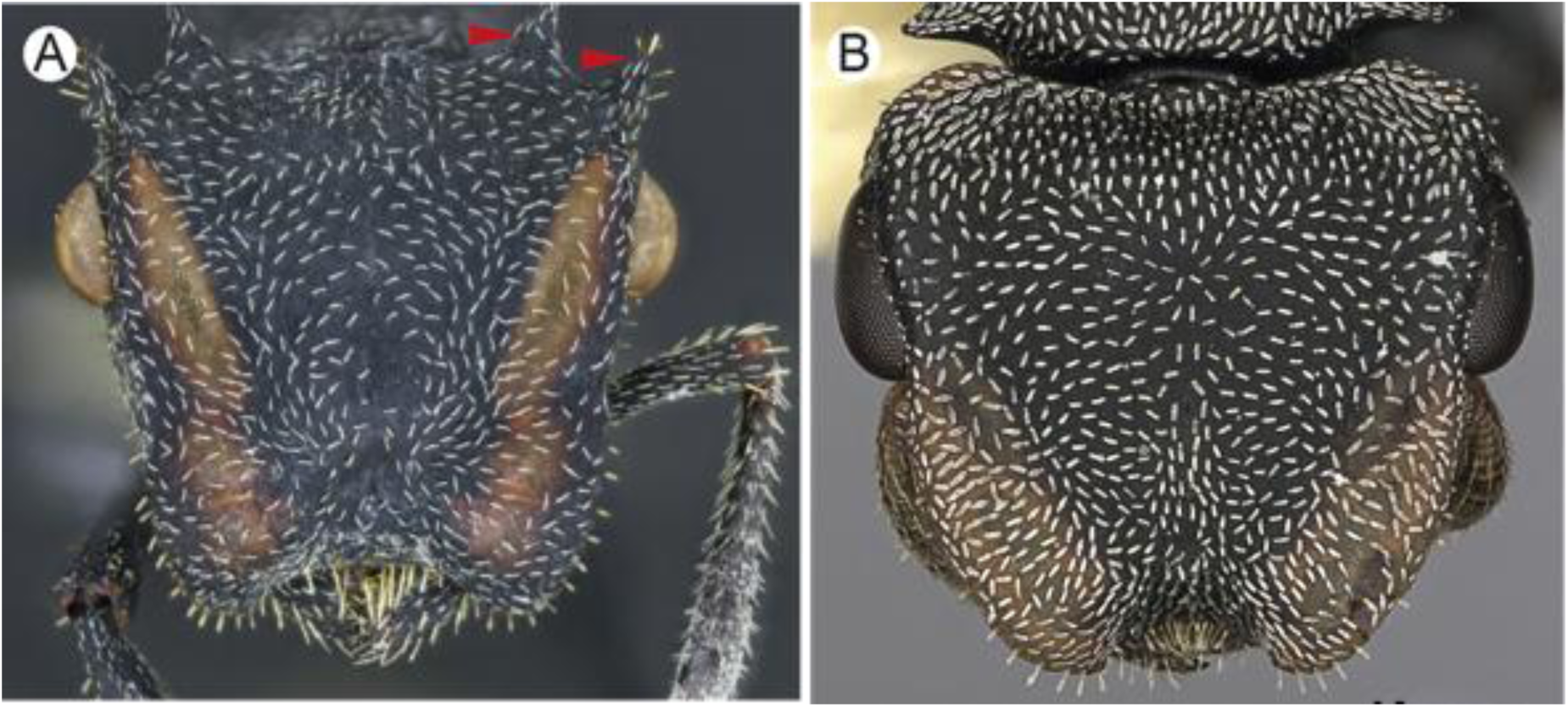
Posterolateral margin of head (full-face view): **A**. Spiny. Image by Akayky Yumbay. **B.** Without spines Image by Will Ericson.

**Figure 10.**
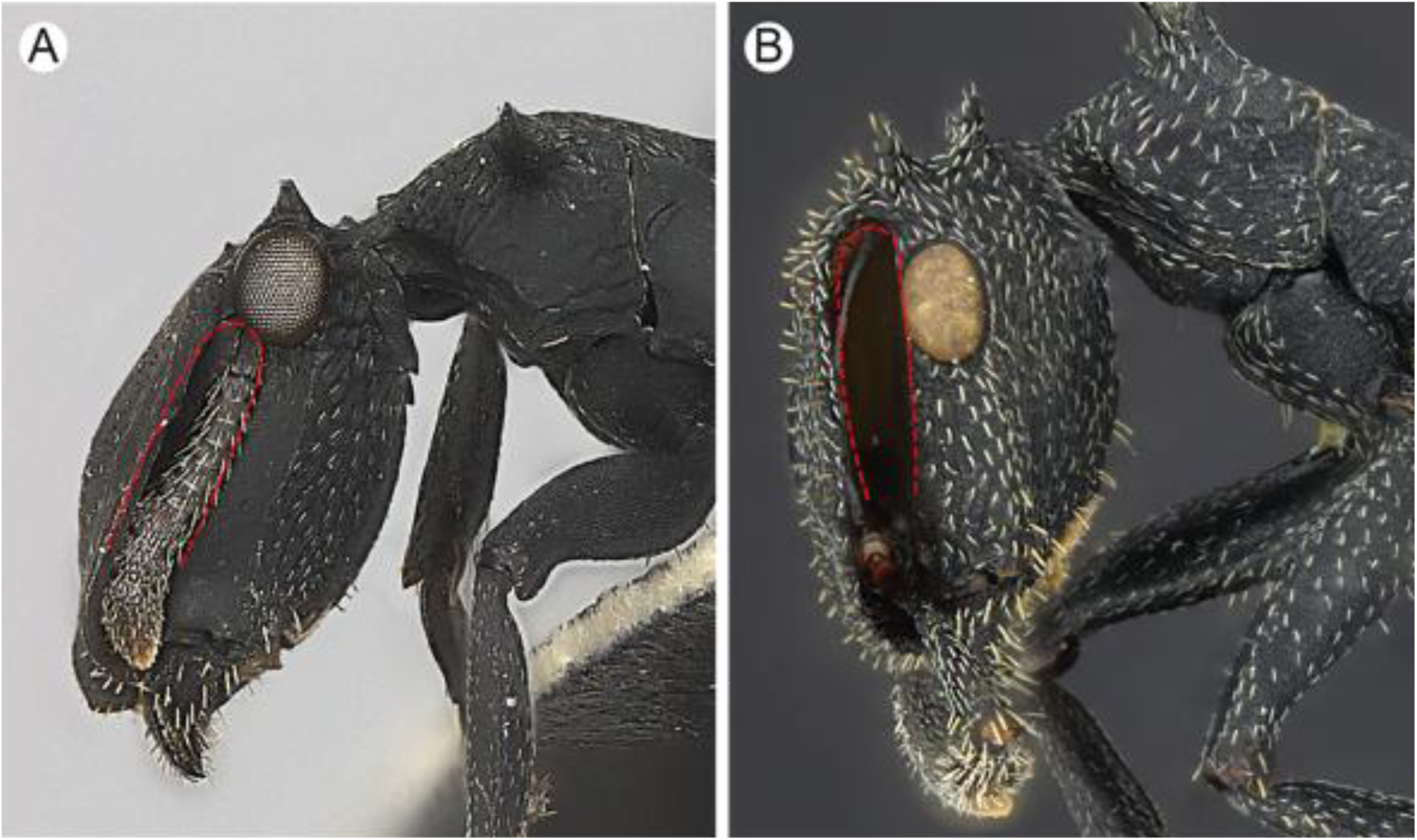
Eye placed (lateral view): **A.** Behind scrobe. Image by Shannon Hartman. **B.** Under scrobe. Image by Akayky Yumbay.

**Figure 11.**
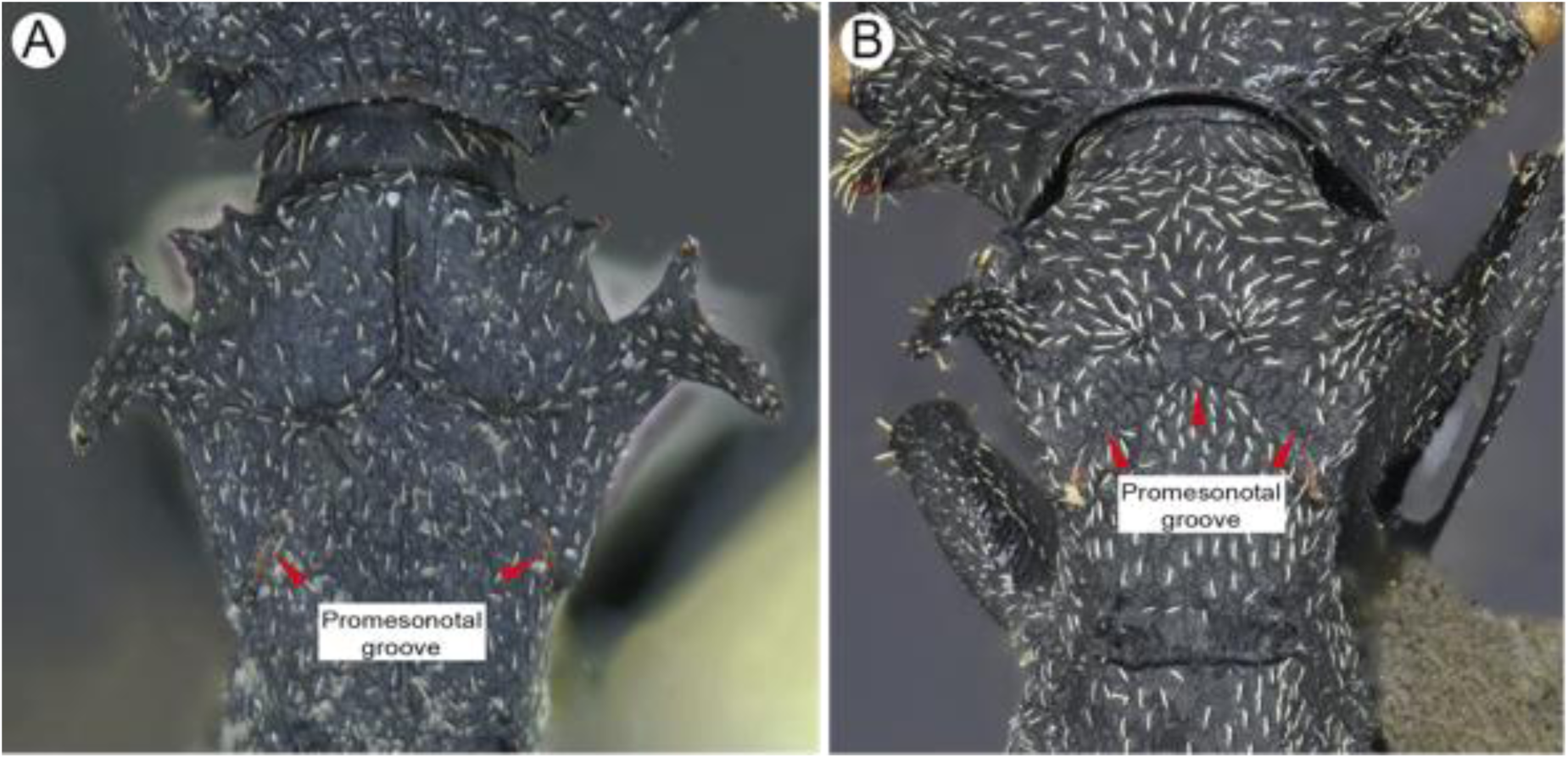
Pronotum (dorsal view): **A**. With inverted Y-shaped carina. **B.** Plain. Images by Akayky Yumbay.

**Figure 12.**
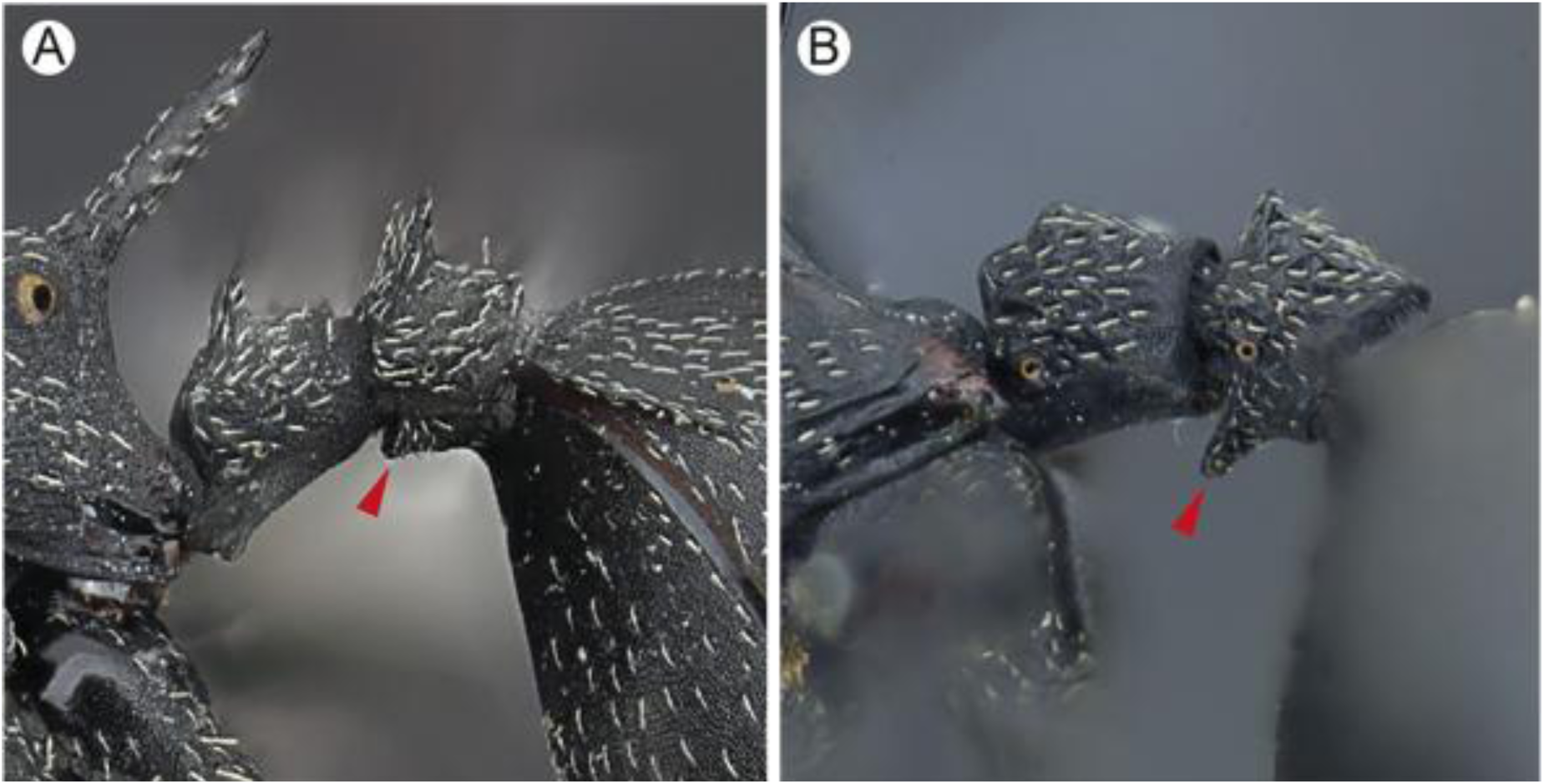
Subpostpetiolar process (lateral view): **A**. Short. **B**. Elongated. Images by Akayky Yumbay.

**Figure 13.**
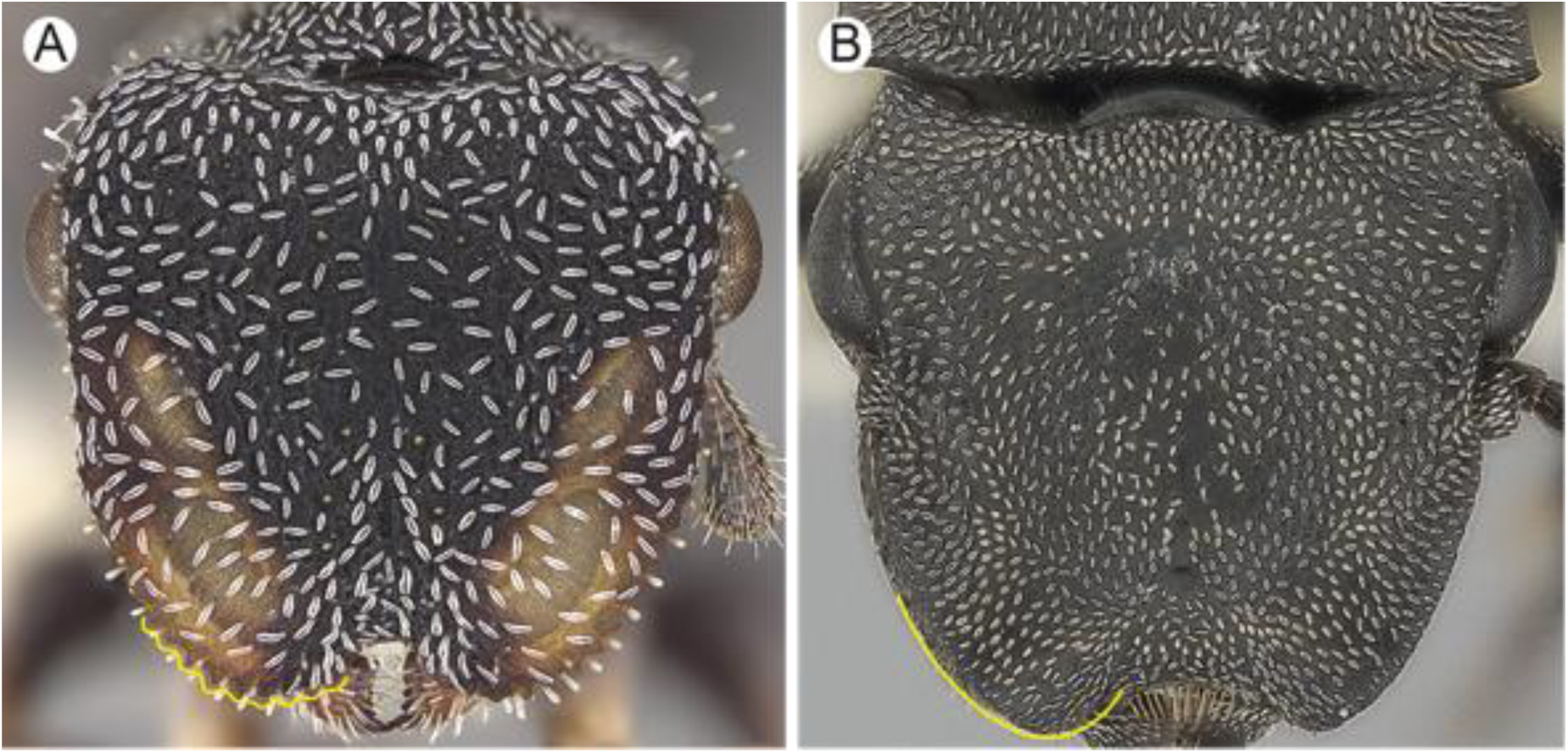
Anterolateral border of frontal carina (full-face view): **A.** Crenulate. Image by Wade Lee. **B**. Plain. Image by Shannon Hartman.

**Figure 14.**
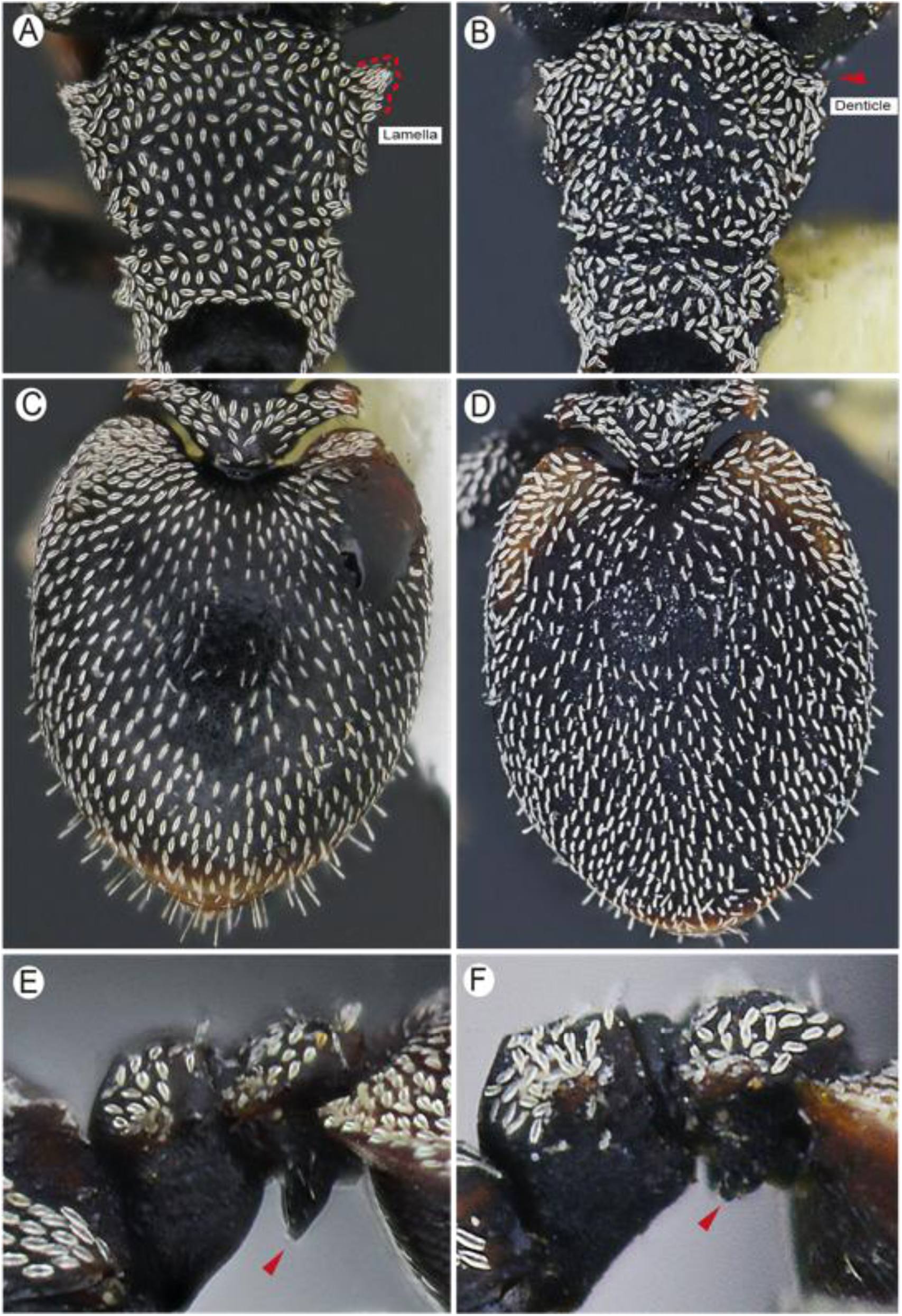
Lateral pronotal margin and gastric integument hue (dorsal view): **A.** Lamellate. **B.** Denticulate. **C.** Shiny. **D.** Opaque. Subpostpetiolar process (lateral view): **E.** Elongated. **F.** Short Images A, B by Akayky Yumbay; C – F by Adrian Troya.

**Figure 15.**
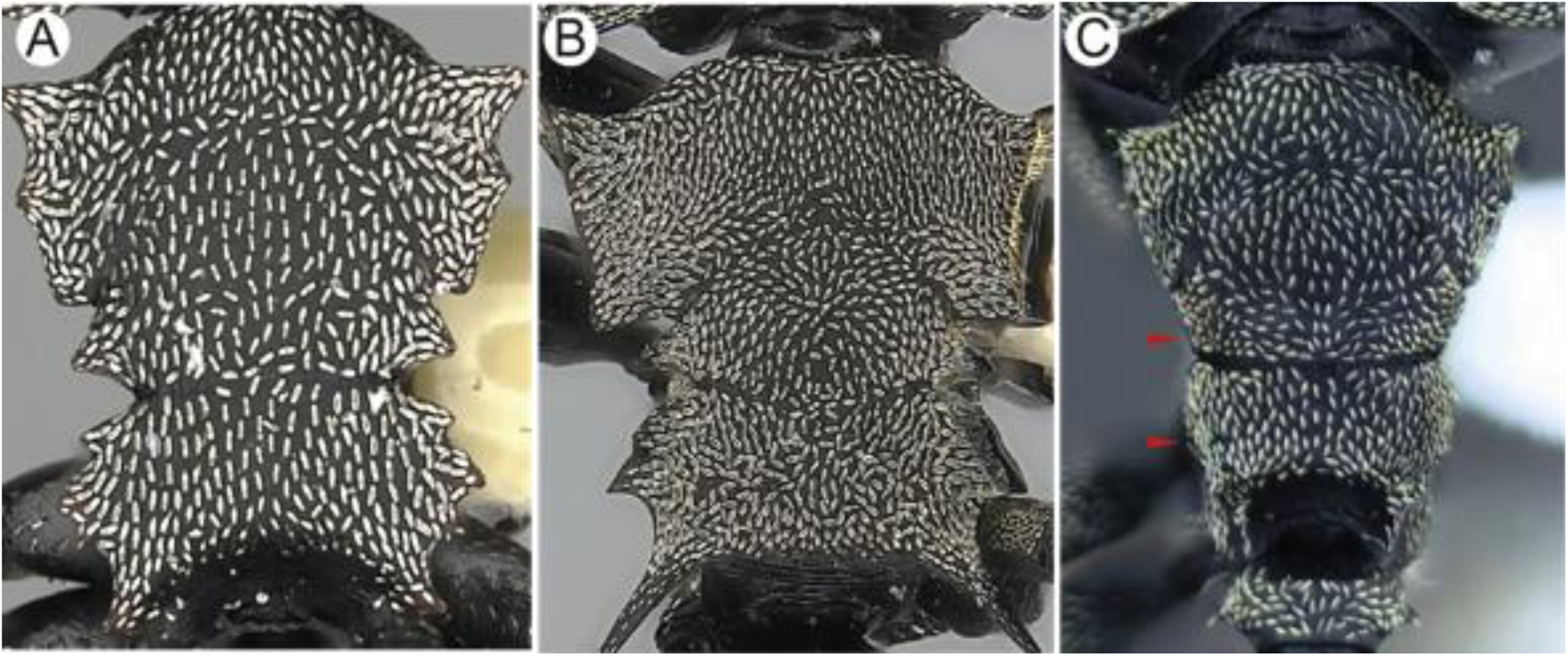
Lateral mesonotal and propodeal margins: **A**. Denticulate. Image by Will Ericson. **B.** Spiny. Image by Shannon Hartman. **C.** Plain. Image by Akayky Yumbay.

**Figure 16.**
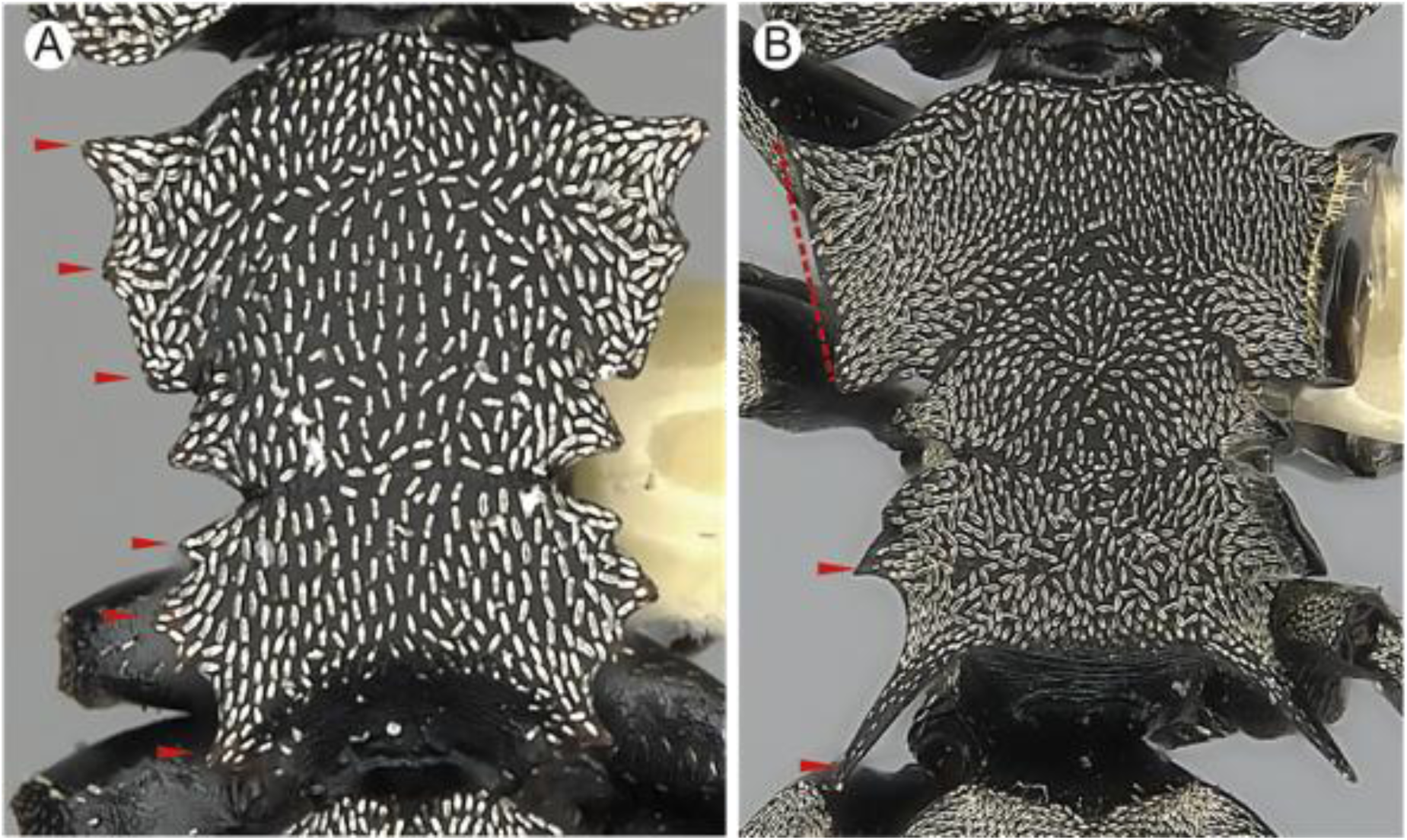
Lateral pronotal and propodeal margins (dorsal view): **A.** Denticulate (red arrowheads). Image by Will Ericson. **B.** Lamellate pronotum (red dotted line), spiny propodeum (red arrowheads). Image by Shannon Hartman.

**Figure 17.**
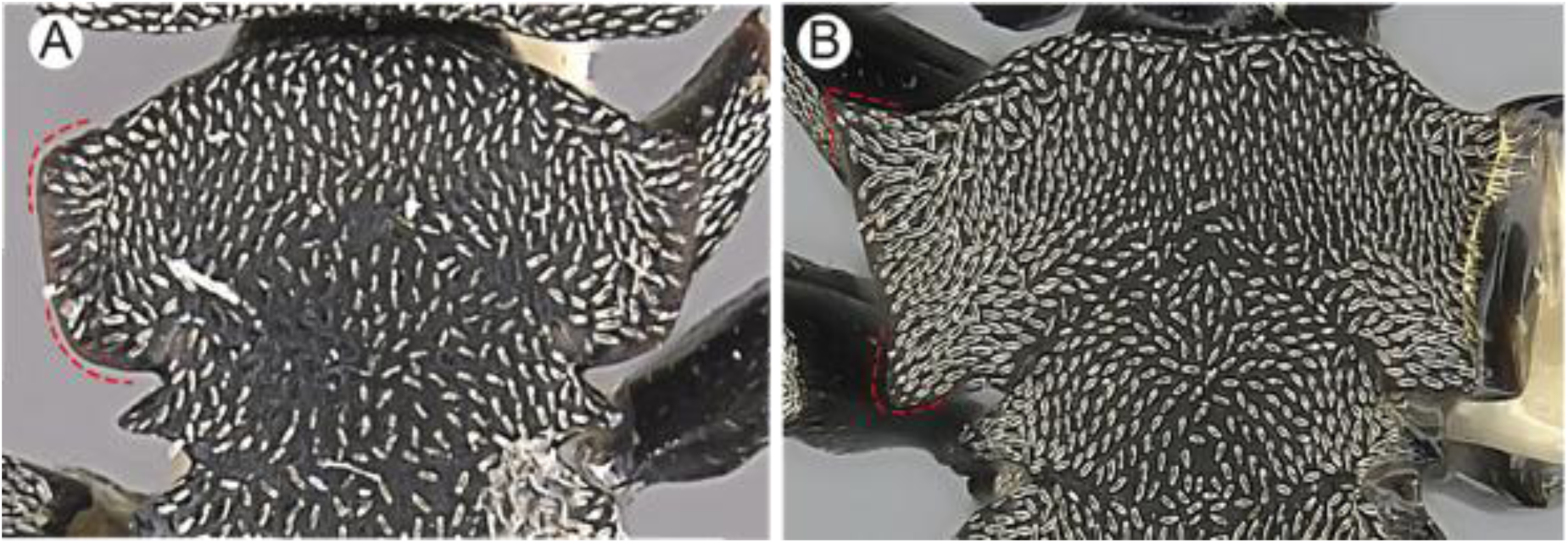
Humeral lamellae (dorsal view): **A.** Subacute to rounded anteriorly, rounded posteriorly. Image by Will Ericson. **B**. With spine anteriorly, acute posteriorly. Image by Shannon Hartman.

**Figure 18.**
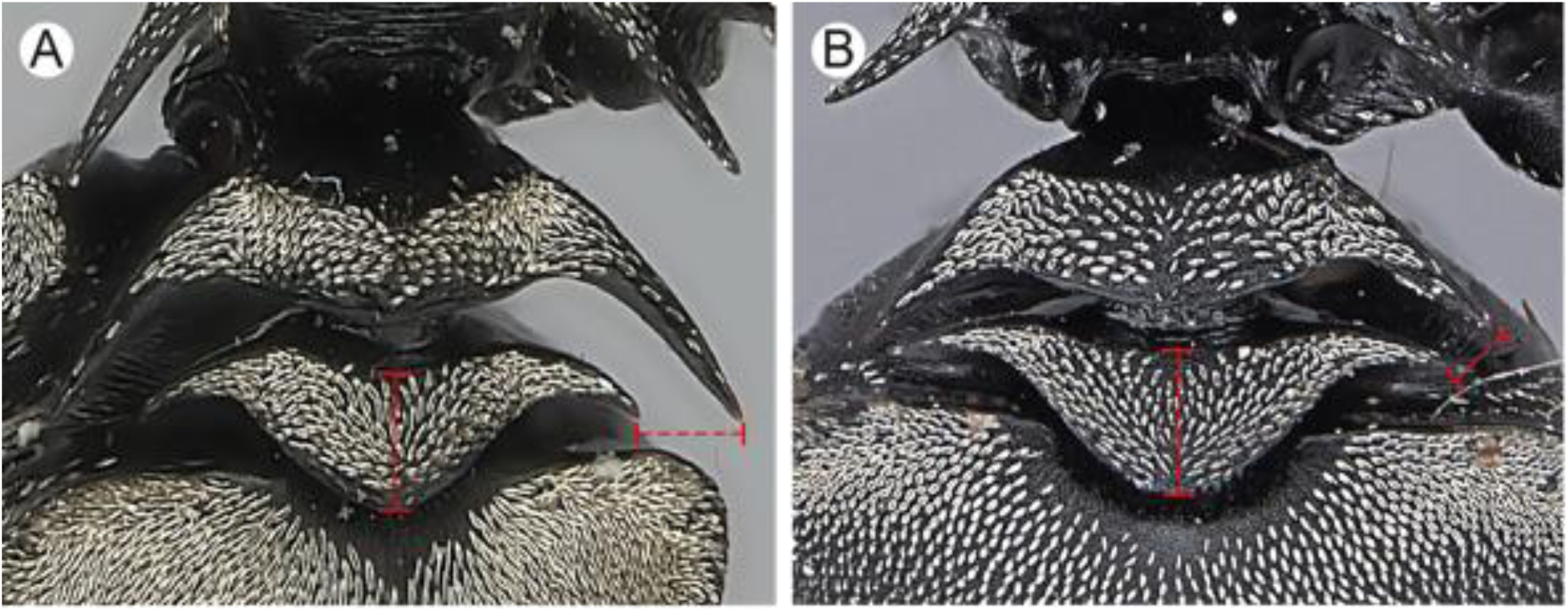
Distance from petiolar spinal tip to postpetiolar spinal tip as compared to mid postpetiolar length. **A**. Subequal. Image by Shannon Hartman. **B.** Shorter. Image by Alexandra Westrich.

**Figure 19.**
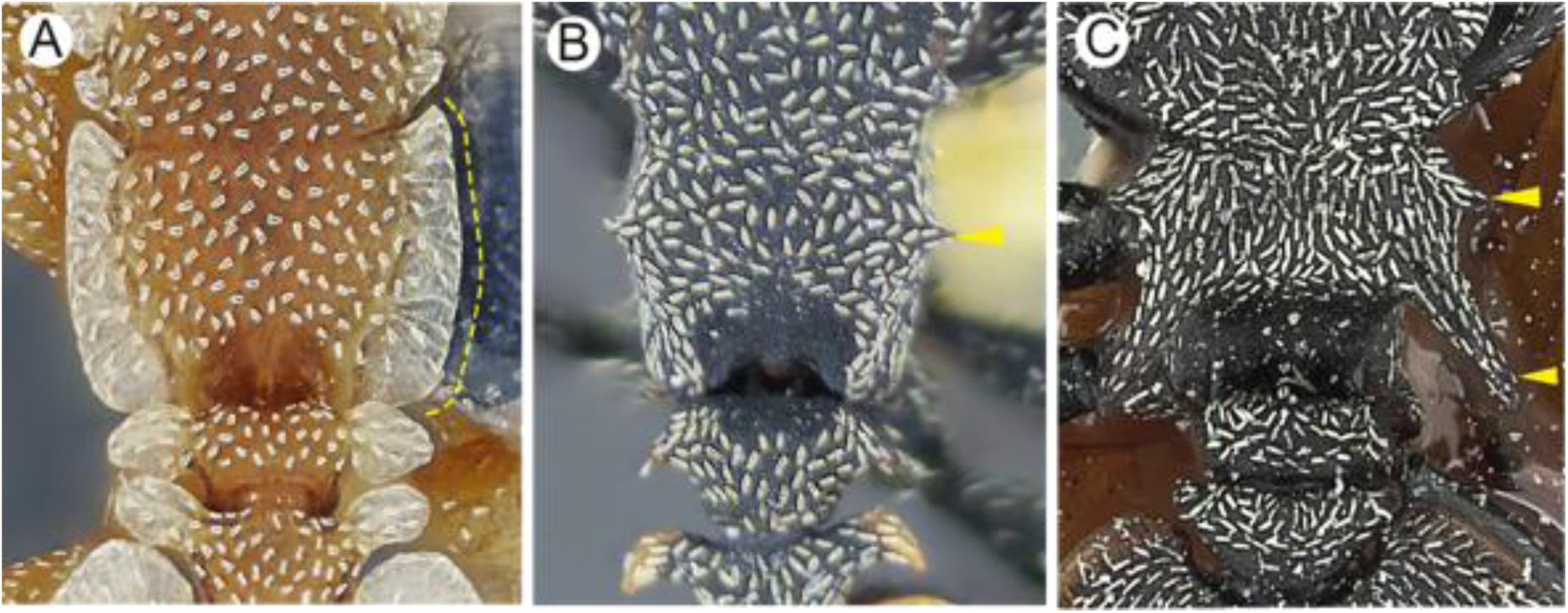
Lateral propodeal margin (dorsal view): **A**. Lamellate. Image by Elian Ilguan. **B.** Denticulate. Image by Akayky Yumbay. **C.** Spiny. Image by Zack Lieberman.

**Figure 20.**
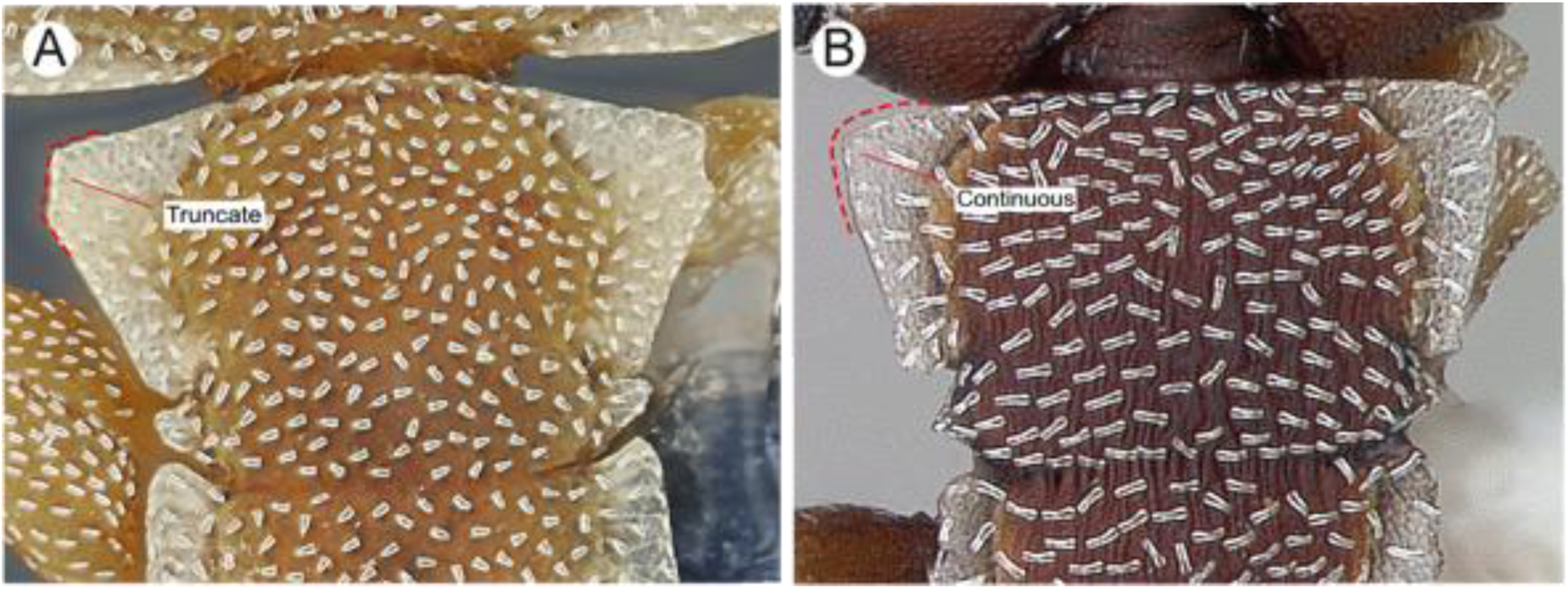
Pronotal lamella (dorsal view): **A.** Truncate. Image by Elian Ilguan. **B**. Continuous. Image by April Nobile.

**Figure 21.**
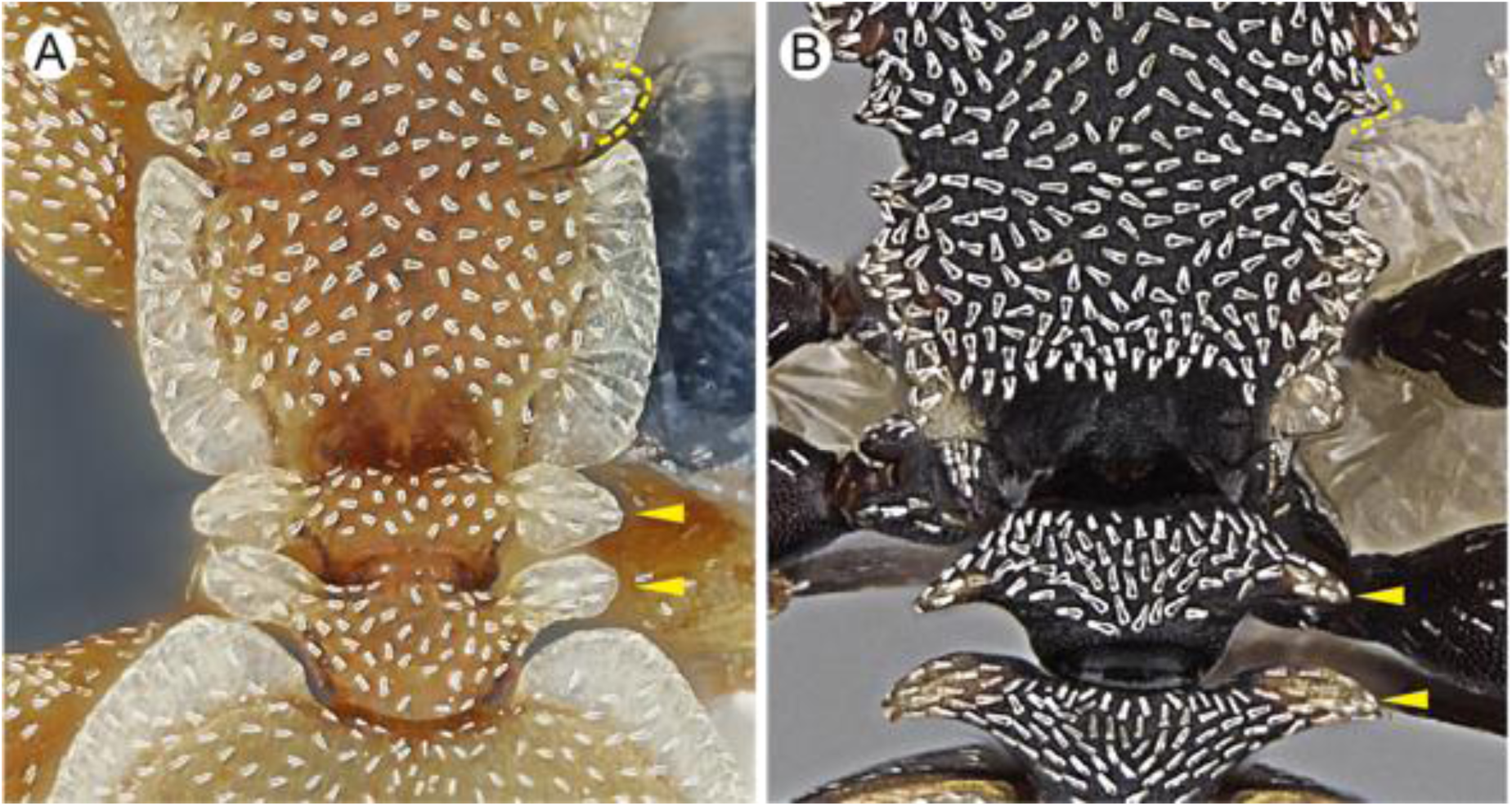
Mesonotal and petiolar lateral projections (dorsal view): **A.** Lamellate (yellow dotted line and arrowheads). Image by Elian Ilguan. **B**. Denticulate (yellow dotted line and arrowheads). Image by Wade Lee.

**Figure 22.**
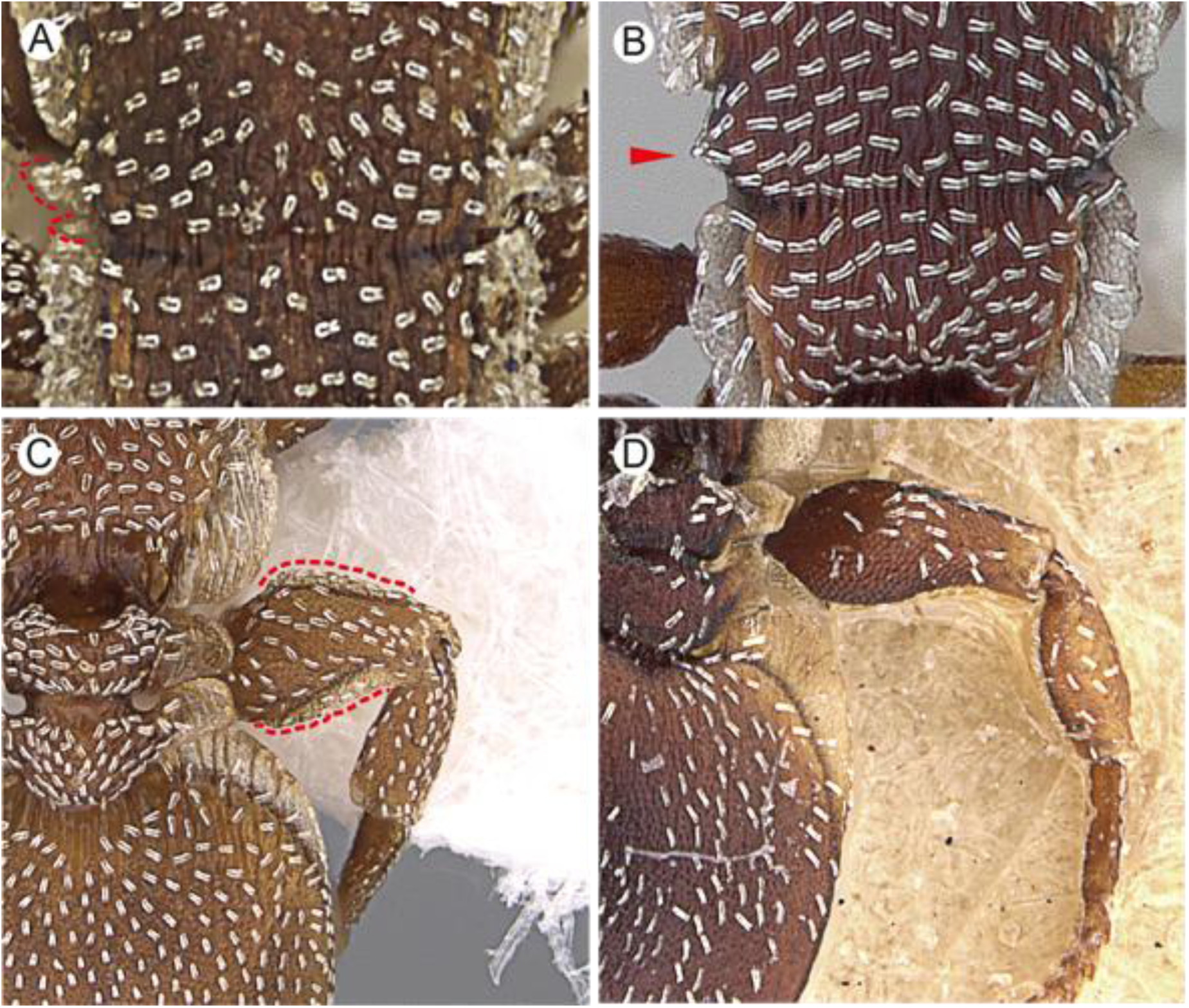
Lateral mesonotal margin (dorsal view) **A.** Lamellate. Image by Ryan Perry. **B**. Denticulate. Image by April Nobile. Hind femur (dorsal or ventral view): **C**. Lamellate. Image by Wade Lee. **D.** Without lamella Image by Will Ericson.

**Figure 23.**
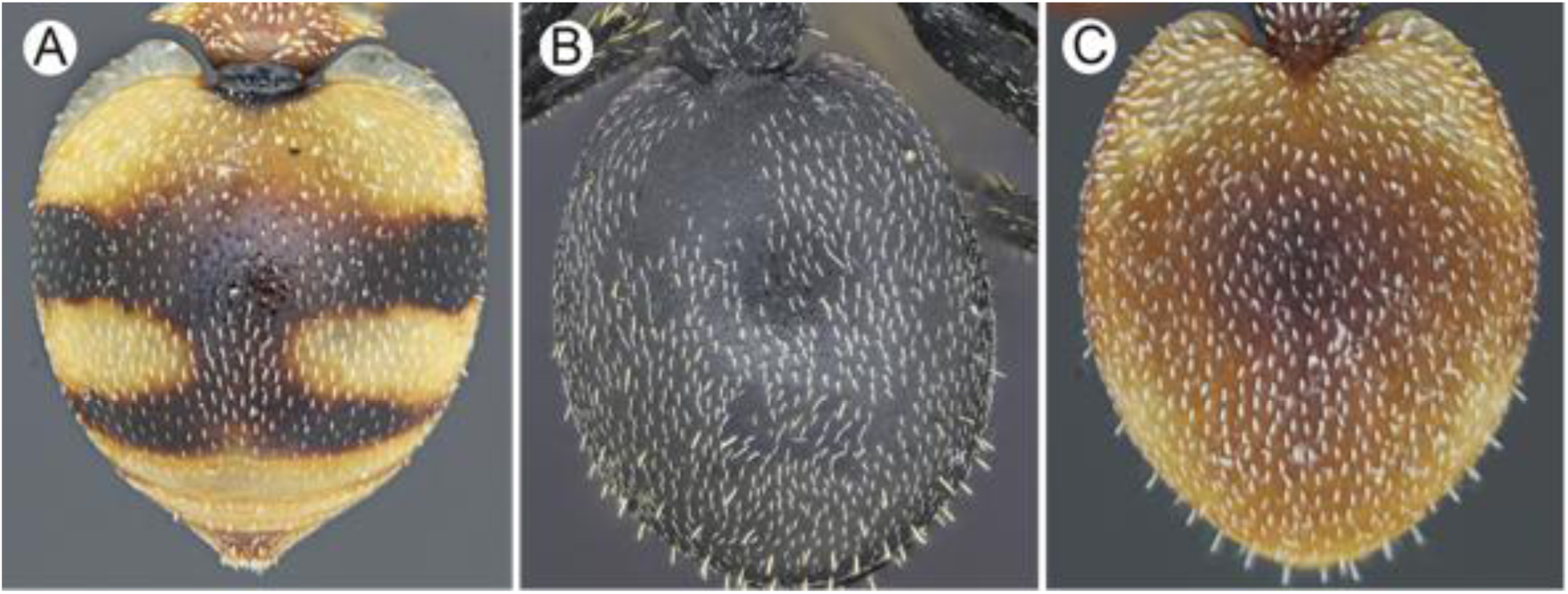
Gaster (dorsal view): **A.** Spotted **B.** Concolored. **C.** Heterocolored. Images by Akayky Yumbay.

**Figure 24.**
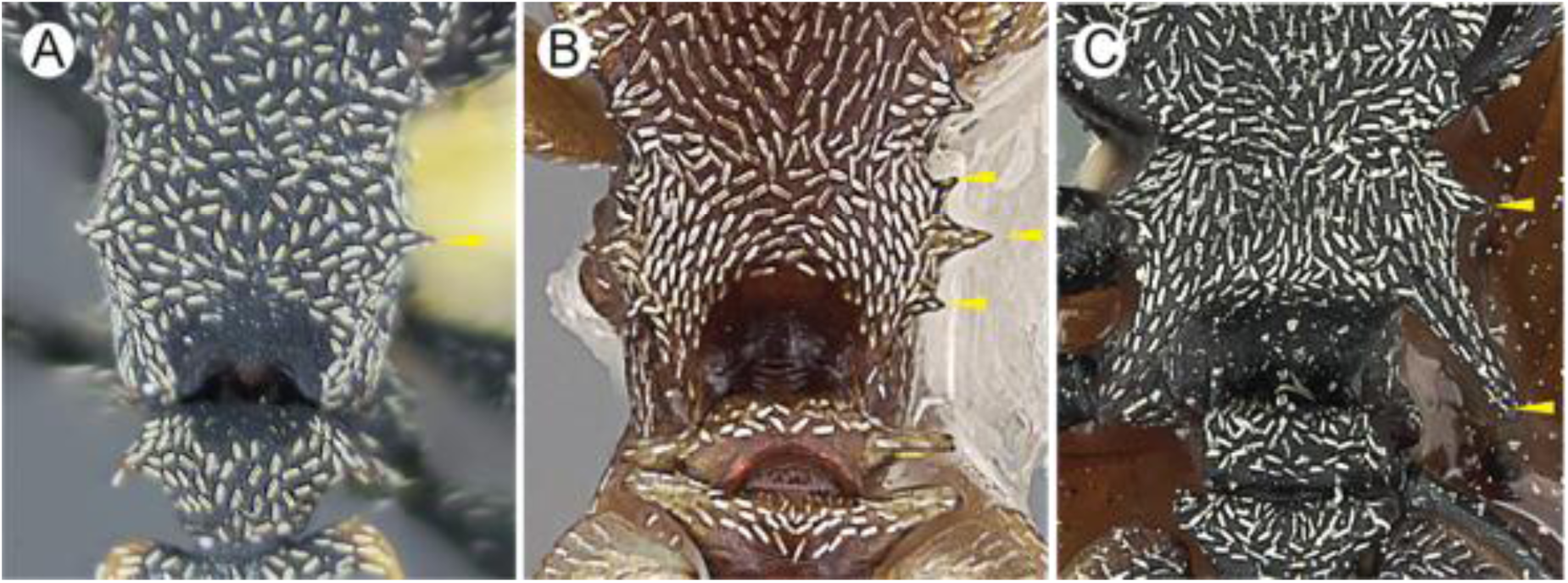
Lateral propodeal margin (dorsal view): **A.** Denticulate. Image by Akayky Yumbay. **B.** Tridenticualate. Image by Wade Lee. **C**. Spiny. Image by Zack Lieberman.

**Figure 25.**
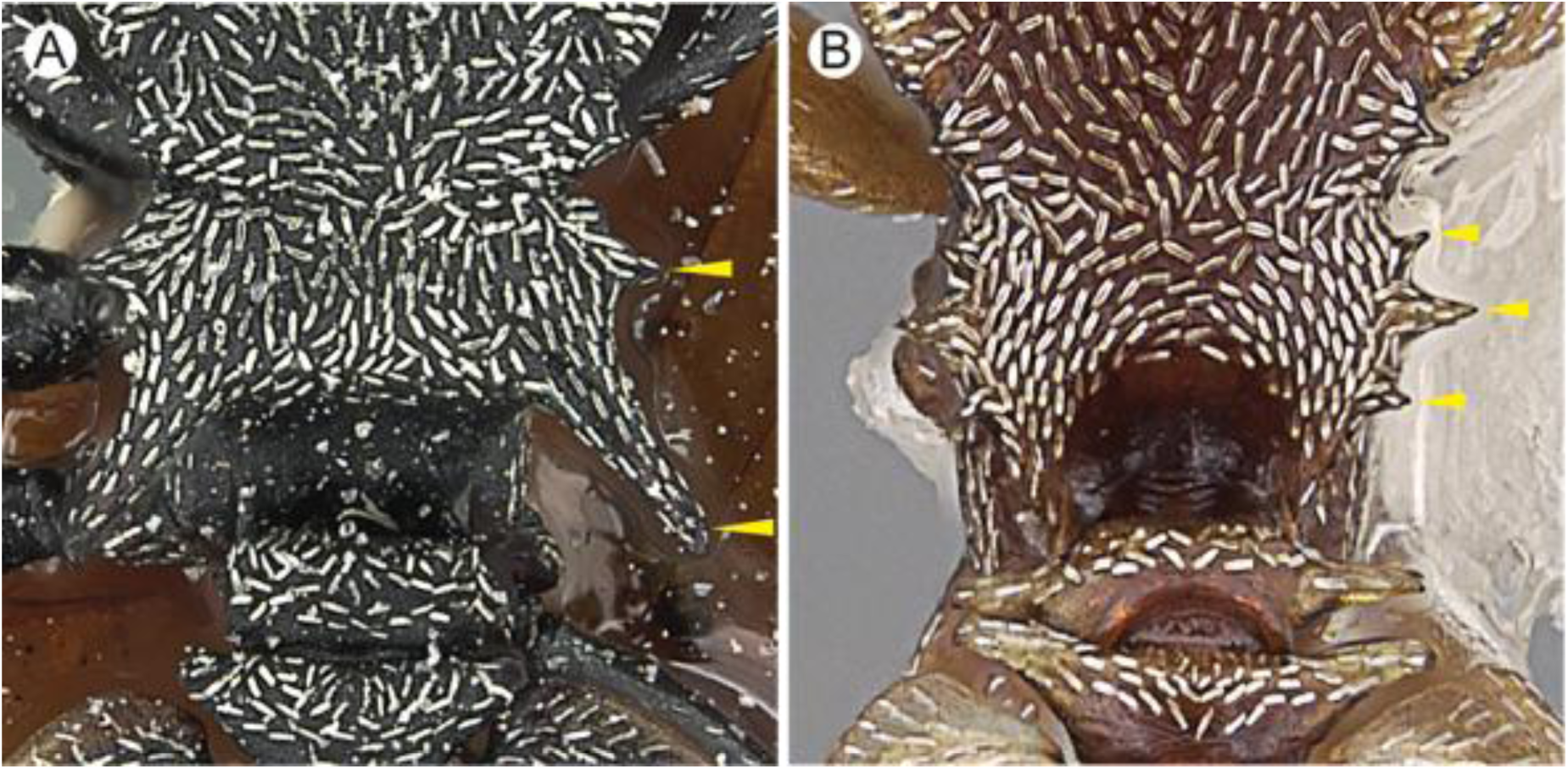
Lateral propodeal margin (dorsal view): **A.** Bispinose. Image by Zack Lieberman. **B**. Tridenticulate Image by Wade Lee.

**Figure 26.**
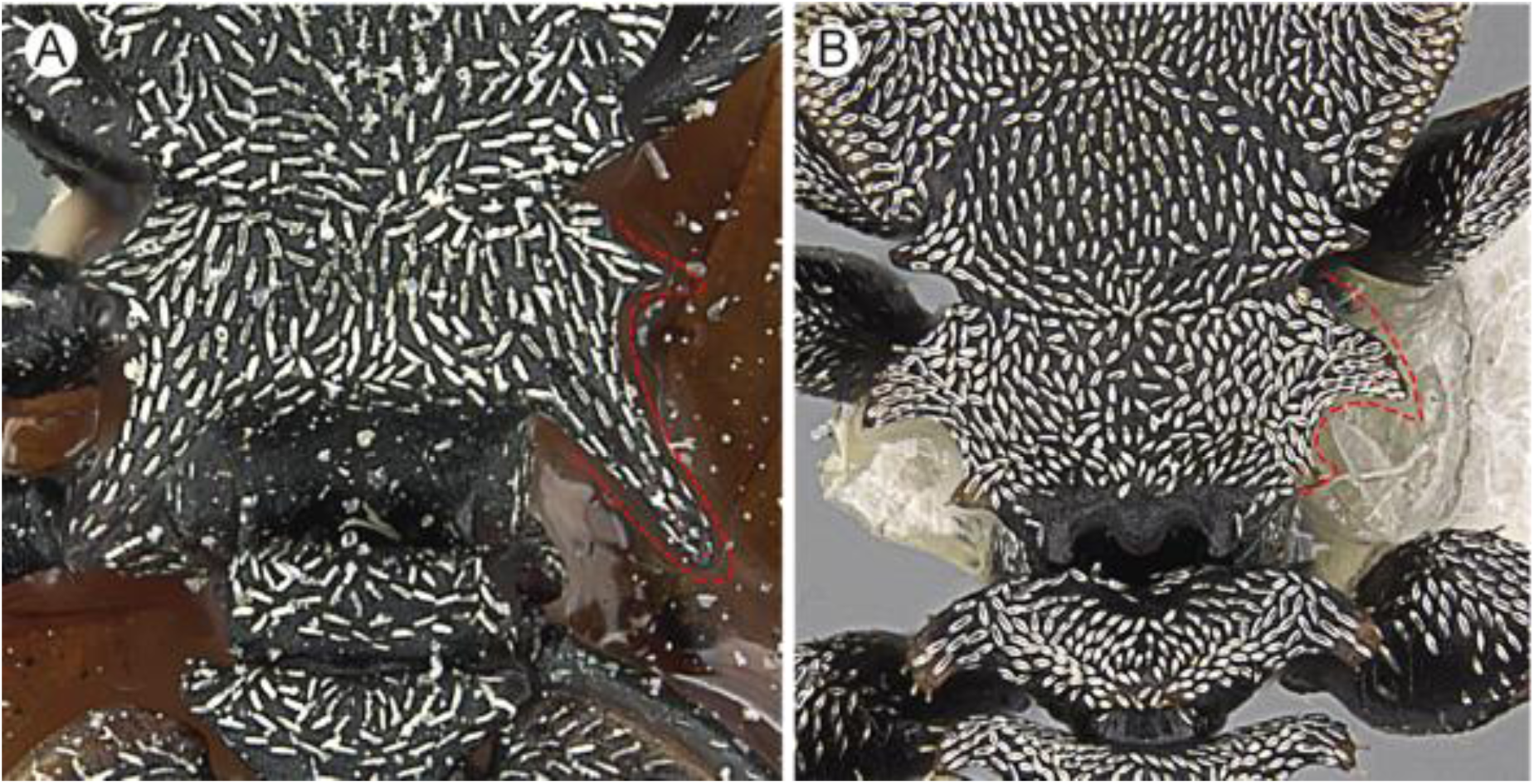
Posterior propodeal spine as compared to anterior spine (dorsal view): **A**. Longer. Image by Zack Lieberman. **B**. Shorter. Image by Wade Lee.

**Figure 27.**
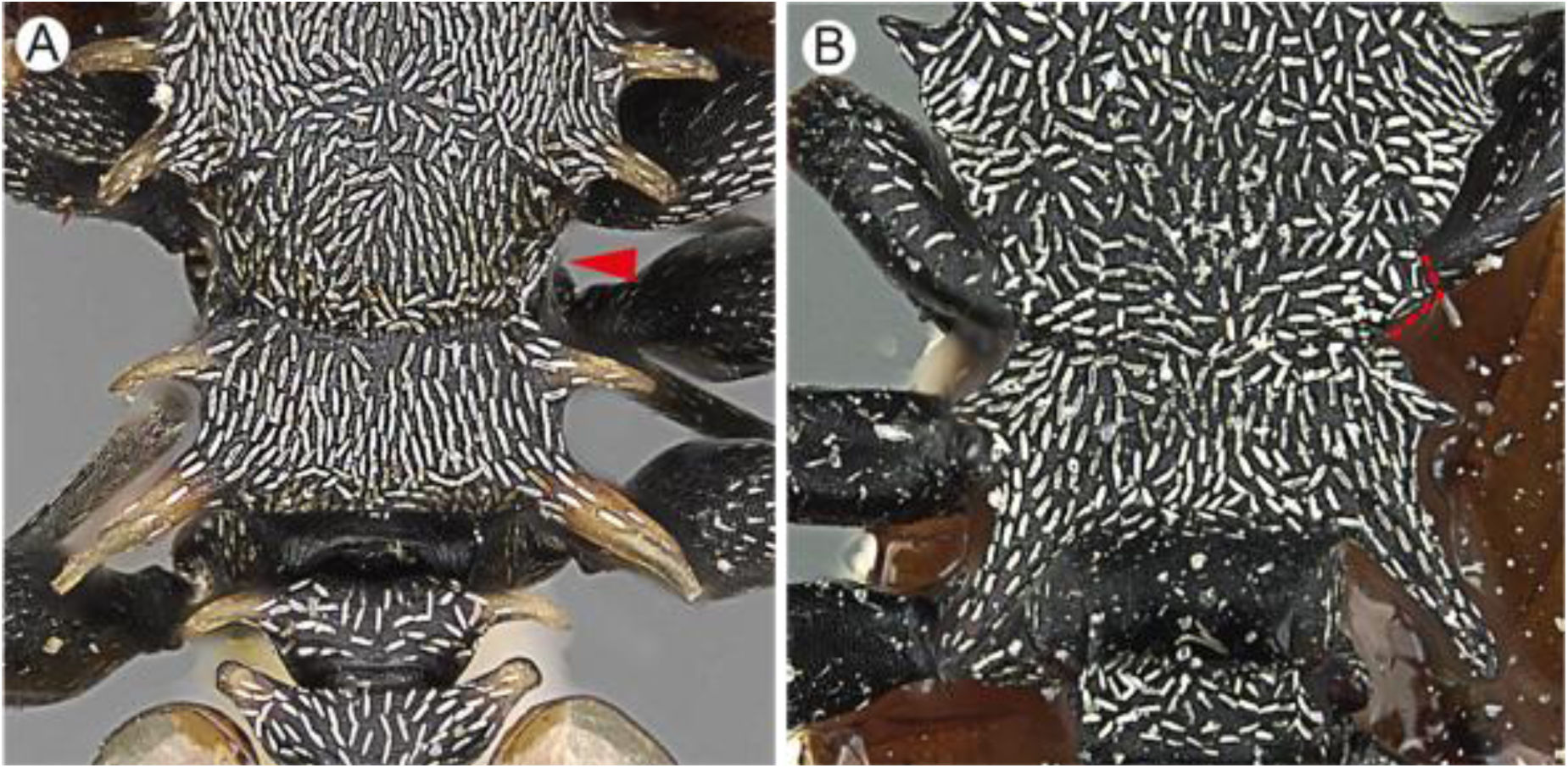
Lateral mesonotal margin (dorsal view)**: A.** Plain. Image by Ryan Perry. **B.** Denticulate. Image by Zack Lieberman.

**Figure 28.**
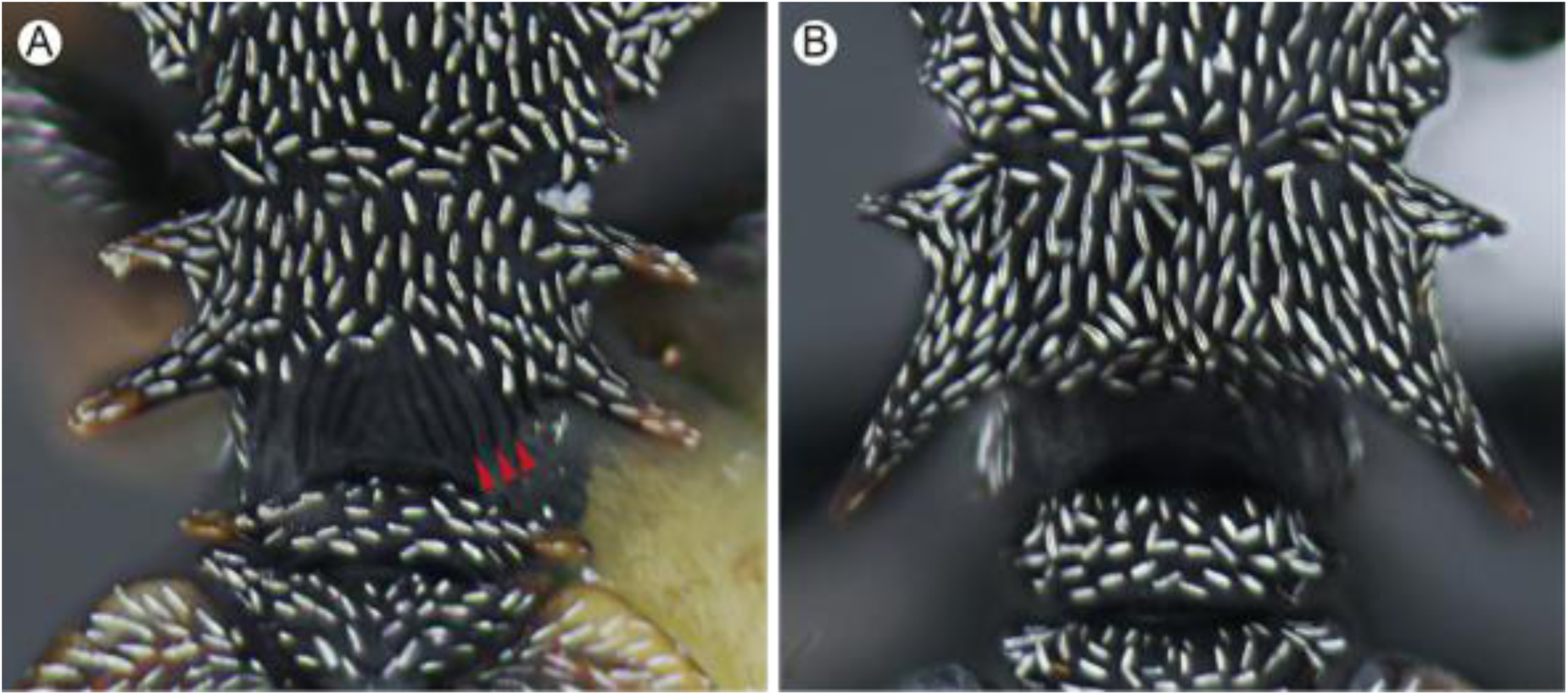
Declivous face of propodeum (dorsal view): **A.** Striate (red arrowheads). **B.** Plain or feebly punctate. Images by Akayky Yumbay.

**Figure 29.**
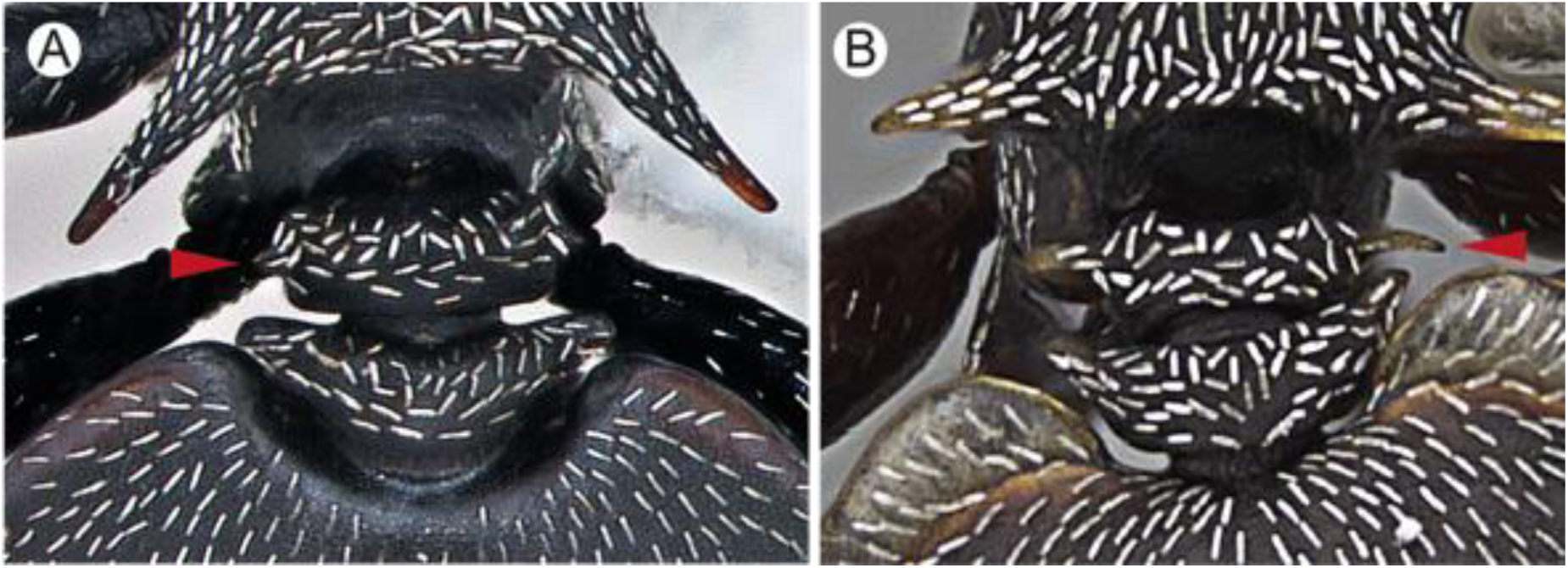
Lateral margin of petiole (dorsal view): **A.** Denticualate. Image by April Nobile. **B.** Spiny. Image by Wade Lee.

**Figure 30.**
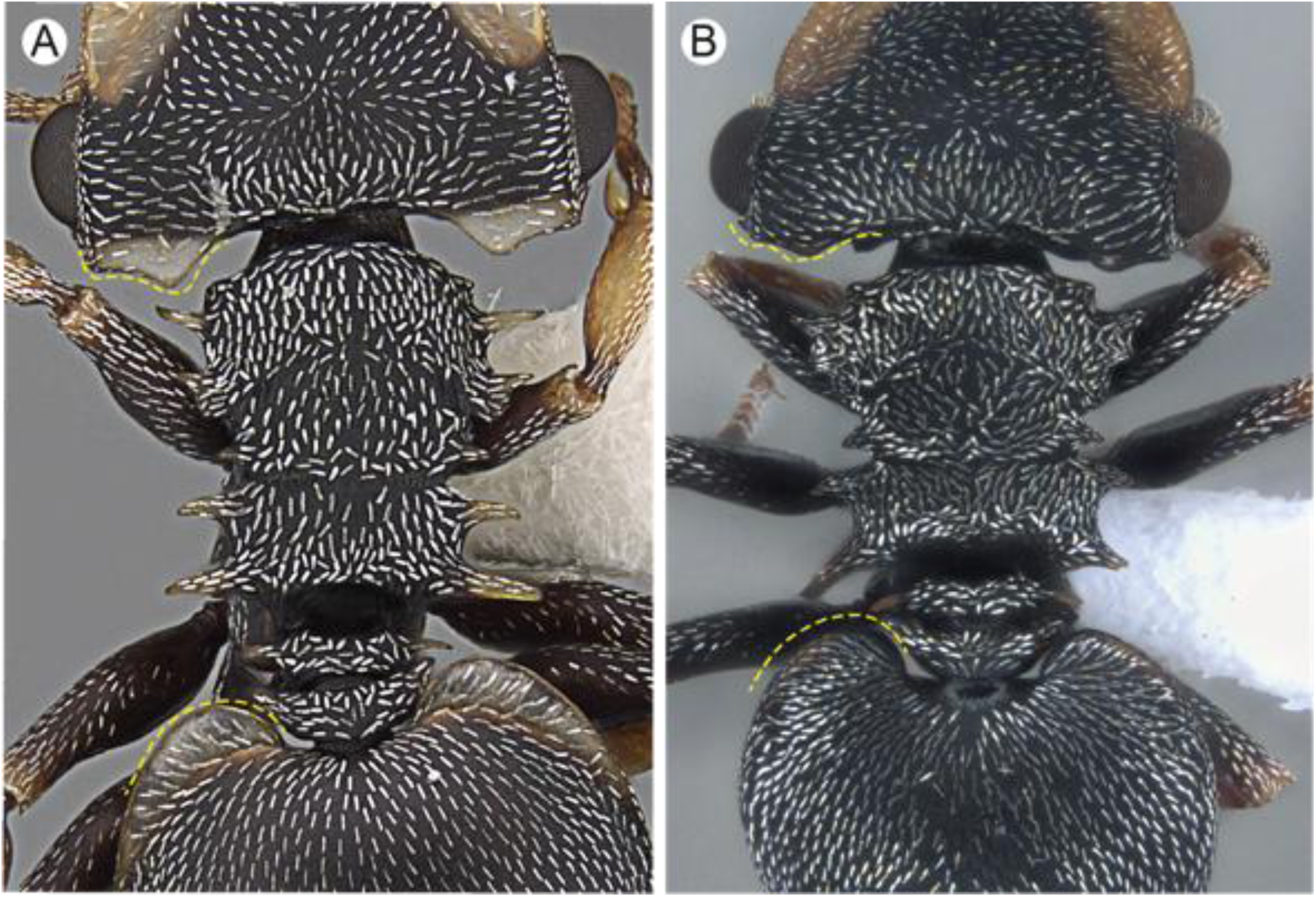
Head and gastric lamellae (dorsal view): **A:** Subhyaline. Image by Wade Lee. **B.** Concolorous to body hue. Image by Frederic Petitclerc.

**Figure 31.**
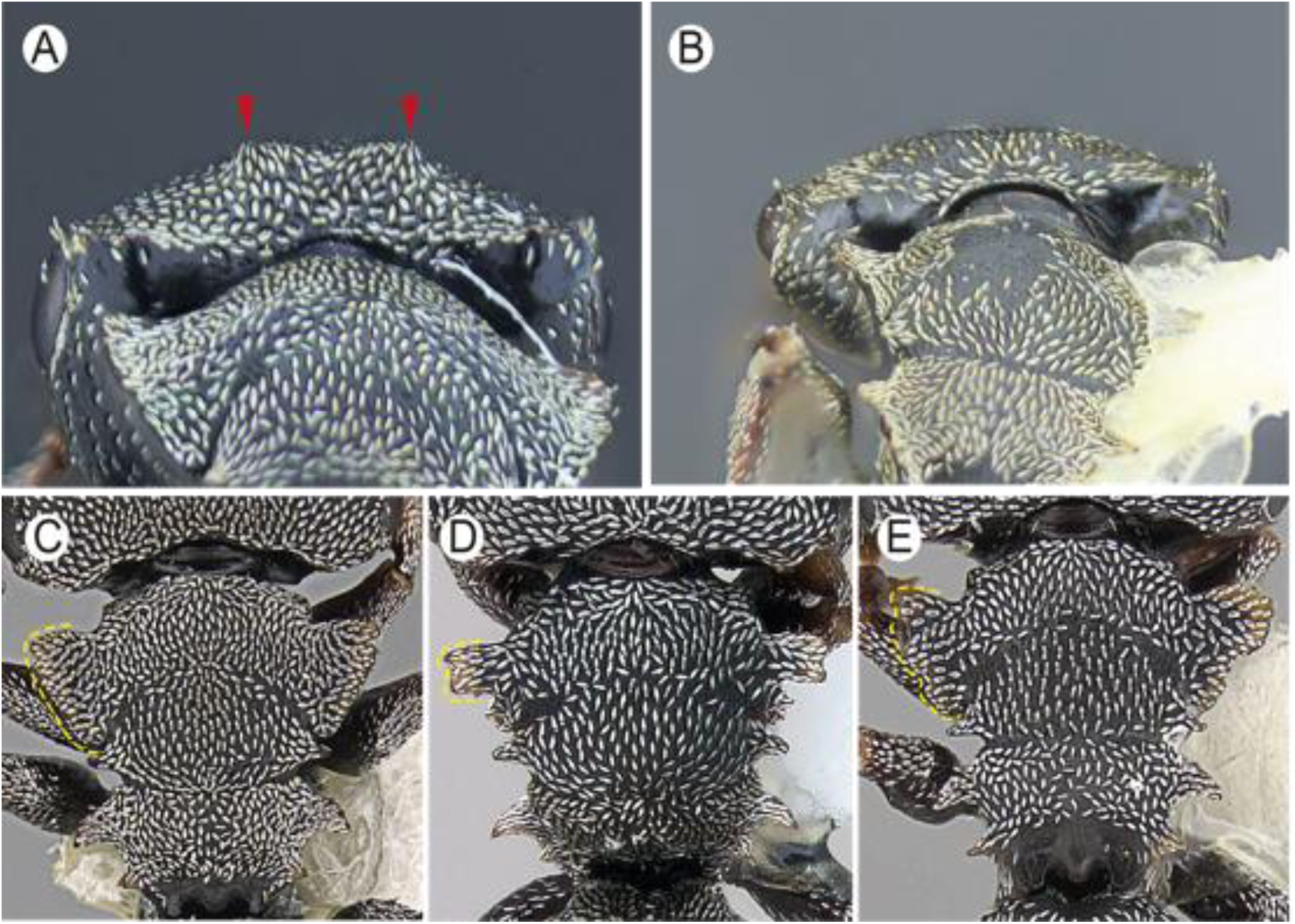
Vertex (posterior view): **A.** Denticulate. **B.** Plain. Images by Akayky Yumbay. Pronotal lamellar margin (dorsal view): **C.** Mostly plain. Image by Wade Lee. **D.** Bidentate. Image by April Nobile. **E.** Crenulate or notched. Image by Wade Lee.

**Figure 32.**
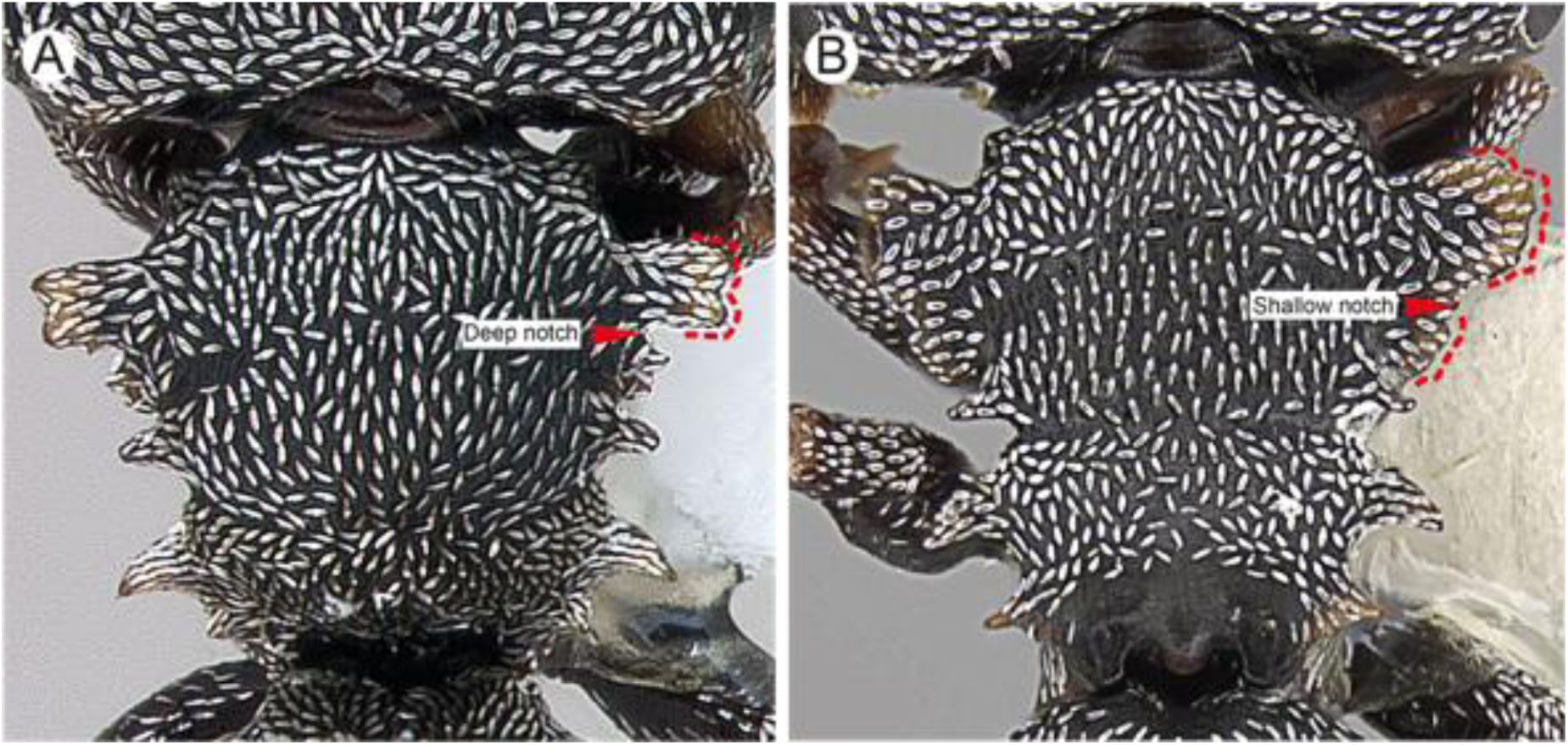
Pronotal lamella (dorsal view): **A**. Bidenticulate (dotted red line), followed by deep notch (arrowhead). Image by April Nobile. **B.** Broad and crenulate, with small notch medially. Image by Wade Lee.

**Figure 33.**
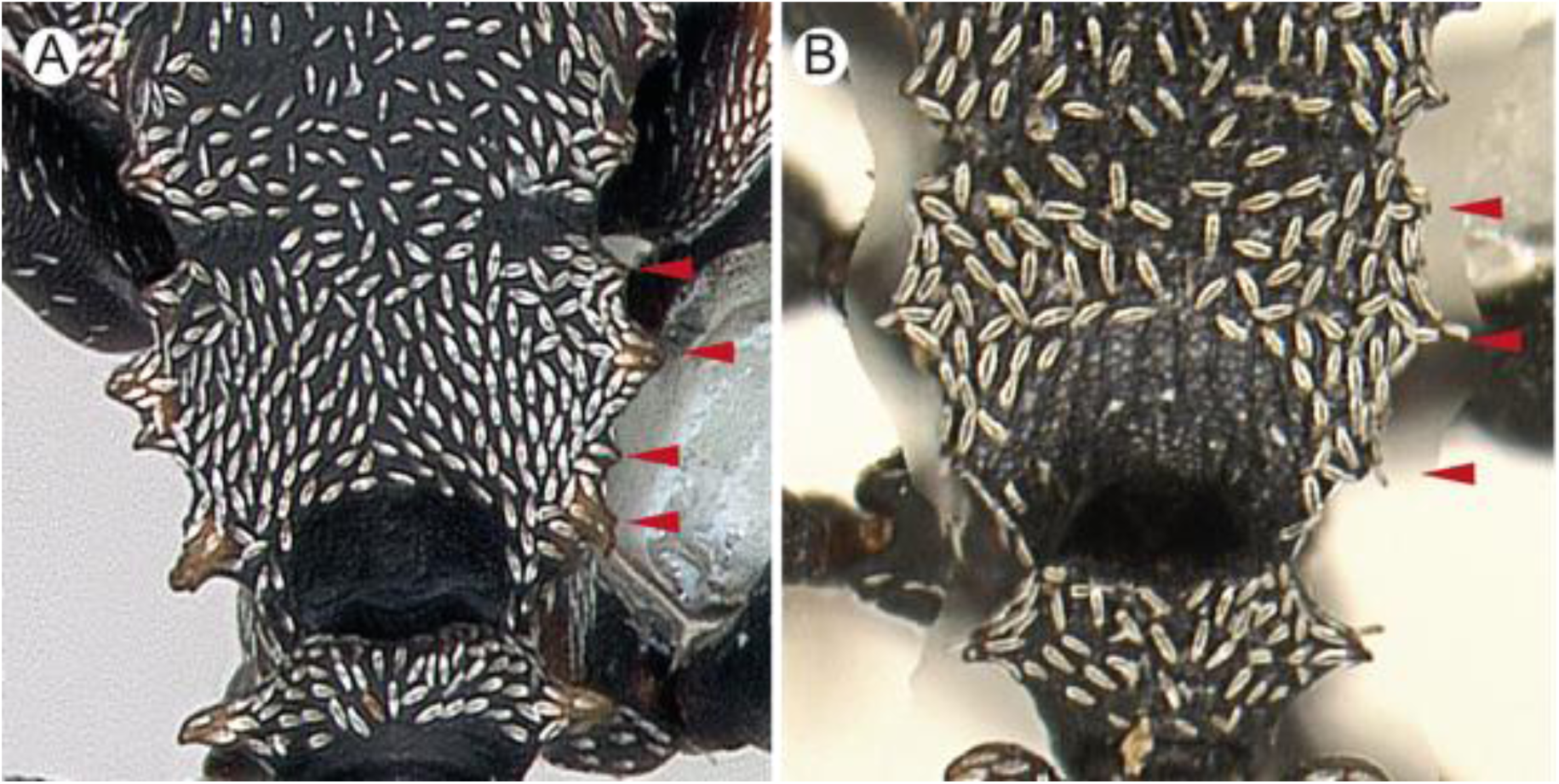
Propodeal denticles (dorsal view): **A.** Four to five. Image by April Nobile. **B**. Three. Image by C. Richart.

**Figure 34.**
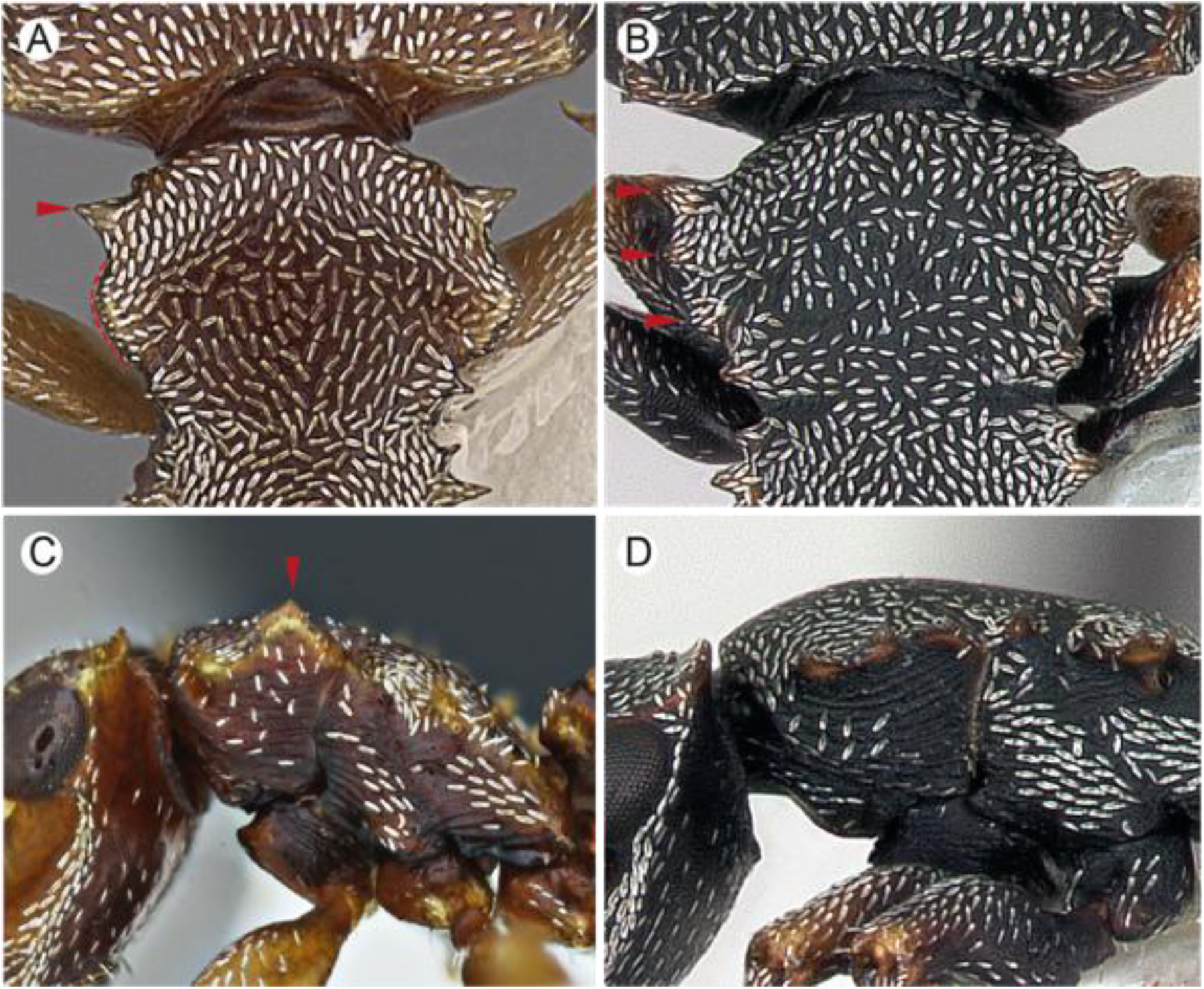
Lateral pronotal margin (dorsal view): **A.** With spine anteriorly (red arrowhead) followed by broad, crenulate lobe upturned. Image by Wade Lee. **B**. With three membranaceous denticles. Image by April Nobile; (lateral view): **C.** Crenulate lobe upturned. Image by Akayky Yumbay. **D.** Without upturned lobe. Image by April Nobile.

**Figure 35.**
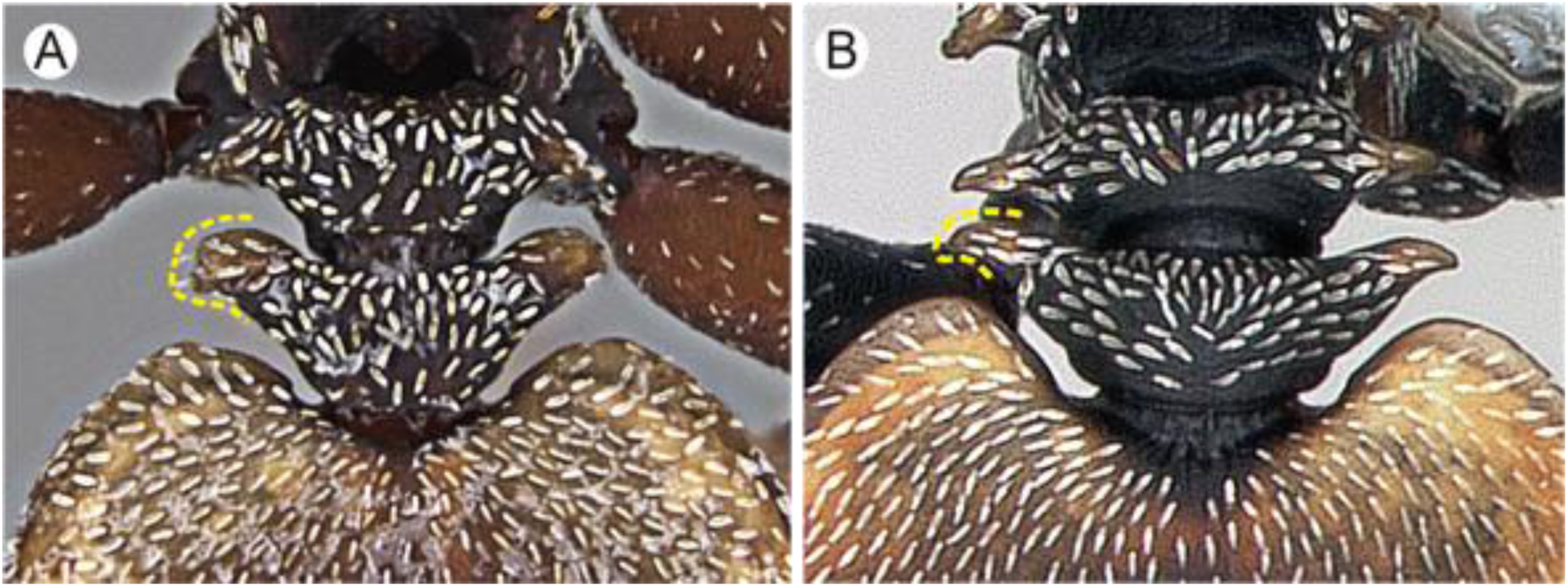
Postpetiolar lateral projection (dorsal view): **A.** Lamellar, wing-shaped. Image by Wade Lee. **B.** Spiny. Image by April Nobile.

*Cephalotes esthelae* Ilguan & Troya sp. nov.

#### Zoobank

Figs 36: A-D (☿ holotype); 37: A-D (☿ soldier); Fig 44C (distribution of records) **Type material.** Holotype. ☿; ECUADOR: Esmeraldas: RECC [Reserva Ecológica Cotacachi Cayapas], Salto de Bravo, 12 km S San Miguel, 10075500 ° N, 726000 ° W [0.682, -78.969], 260 m, Fumigación [method], B. Tierra Firme [habitat], Abril 2001, P. Araujo et al. [leg.] (MEPN: MEPNINVINV4999).

**Figure 36.**
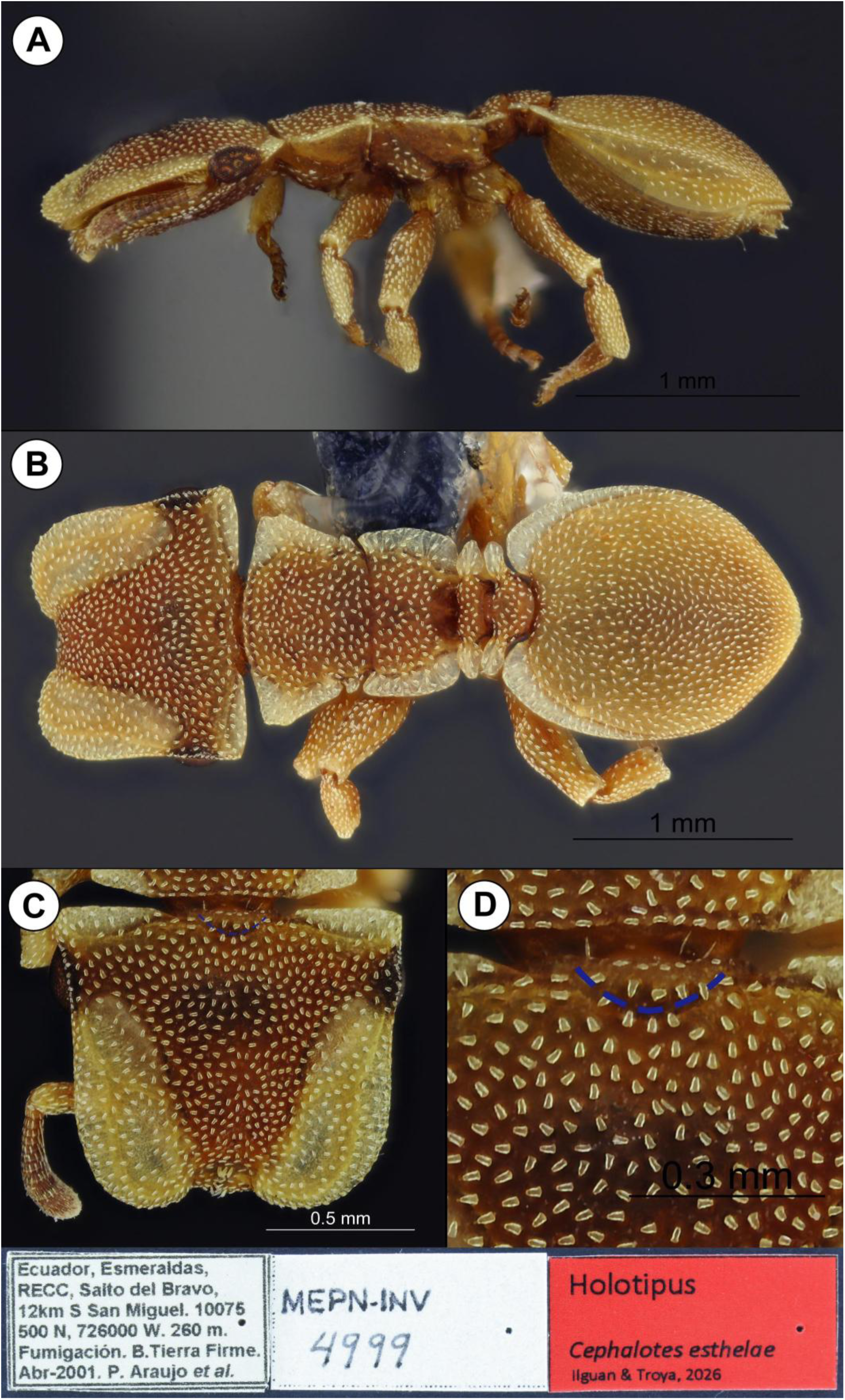
Holotype worker of *Cephalotes esthelae* **sp. nov.** (MEPN: MEPNINV4999) **A.** Full body, lateral view. **B.** Full body, dorsal view. **C.** Head, full-face view. **D.** Close-up of concavity on posteromedian cephalic margin (blue dashed line). Collection labels are shown at the image’s foot. Image by Elian Ilguan.

**Paratypes.** 5 ☿. Same data as holotype (MEPN: ☿ soldier, MEPNINV4996; ICN: ☿ soldier, MEPNINV4997; ICN: ☿, MEPNINV5002; MCZC: ☿, MEPNINV5001); Same data as holotype, except: La Tabla, 7km de Playa de Oro, río Santiago, 00°50’42”N, 78°44’40”W [0.845, -78.7444], 120 m, 01-Abr-2001(MEPN: ☿, MEPNINV28234).

**Etymology.** The specific name is dedicated to Esthela, mother of Elian Ilguan, one of the authors of this work. This is in recognition of her unconditional support and constant guidance in his personal and scientific training. The epithet is formed in the Latin feminine, singular genitive (International Code of Zoological Nomenclature: Art. 31.1.2).

**Diagnosis.** A member of the *pinelii.* + *grandinosus* species-group sensu Price et al. (2022). Workers: Posteromedian cephalic margin concave, in frontal view (Fig. 36C, D); posterior head margin broader than pronotum by about ¾ to one maximum eye diameter (Fig. 36B); propodeal lamellae transversely striate (Fig. 36B). Soldiers: Cephalic disc bigger (CI: 114-130 mm) than all other conspecifics of this species-group; propodeal dorsum slightly higher than mesonotal dorsum, in lateral view (Fig. 37A). Workers and soldiers: Body with abundant foveae bearing simple, mostly triangular-like hairs (Fig. 36B-D, 37B).

**Figure 37.**
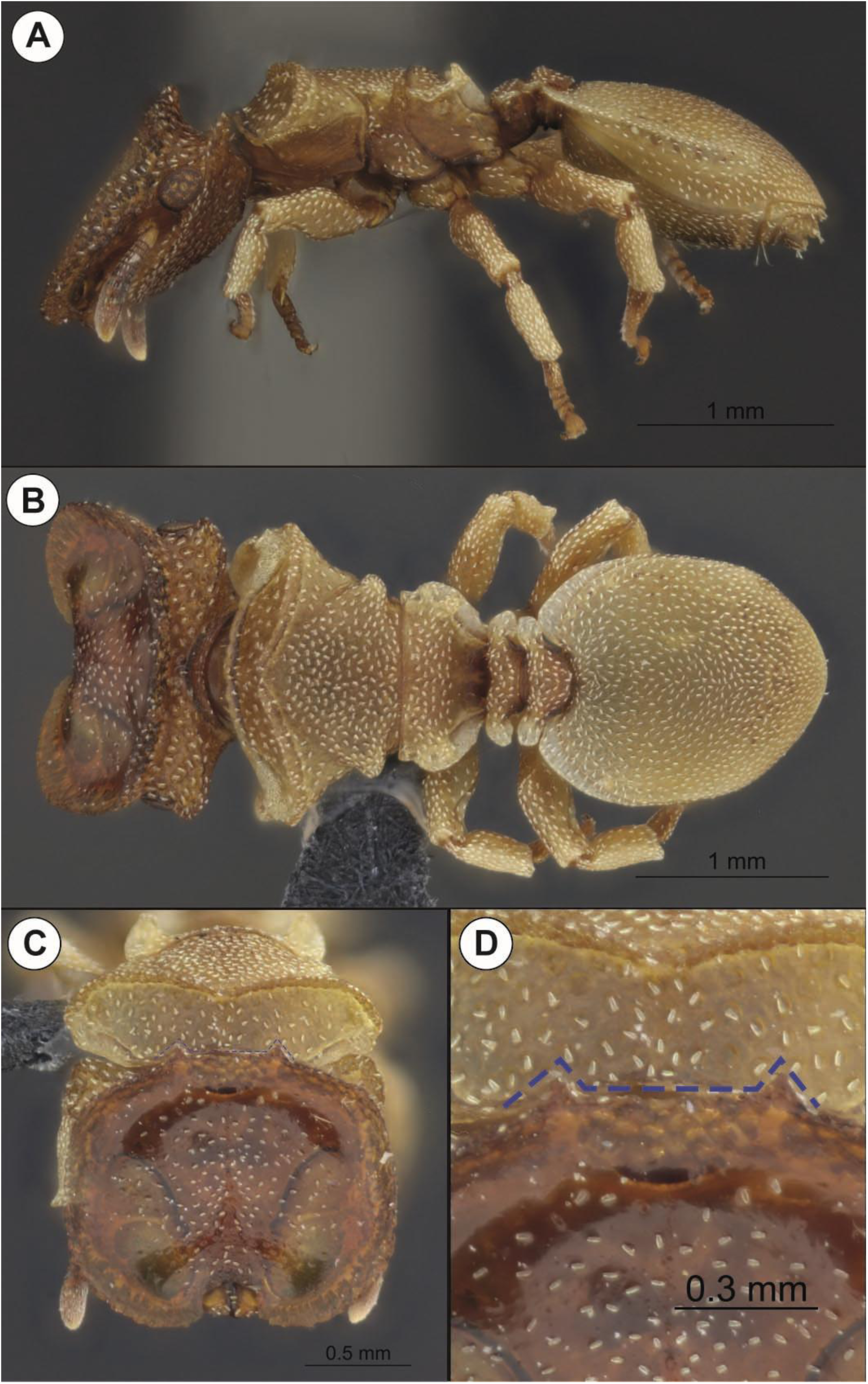
Soldier of *Cephalotes esthelae* **sp. nov.** (MEPN: MEPNINV4996) **A.** Full body, lateral view. **B.** Full, body, dorsal view. **C.** Head, full-face view. **D.** Close-up of teeth on posterior cephalic margin. Image by Elian Ilguan.

**Worker description.** Measurements (*n*=5): HW: 1.19-1.5 (1.3); HL: 0.99-1.19 (1.19); EL:

0.24-0.26 (0.25); CCHA: 0.2-0.24 (0.2); PW: 0.92-1.03 (1); WL: 1.1-1.16 (1.1); PTL:0.18-0.21 (0.18); PTW:0.62-0.66 (0.66); PPL:0.2-0.24 (0.22); PPW:0.57-0.66 (0.64); GL: 1.31-1.51 (1.42); GW:1.14-1.18 (1.18); HBL:0.31-0.34 (0.33); HBW:0.11-0.13 (0.13); Indices. CI: 114.71-130 (125); OI: 19.3-21.61 (19.68); PI:26.67-32.87 (26.67); HBI:37.14-42.86 (38.67) TL: 3.69-4.31 (4.08).

**Head.** *In frontal view:* Subquadrate, wider than long (CI: 115–130). Occipital corner with subhyaline lamella; posterolateral margin of frontal lobe covering most of eye dorsally; posteromedian margin deeply concave; width of posterior margin broader than maximum pronotal width; anterolateral margin feebly convex. Masticatory margin of mandible with one to two apical teeth, followed by three distal denticles, decreasing in size from apex to base. *In lateral view:* Anteromedian dorsal surface feebly convex, followed by slight depression and convexity on vertex (Fig. 36A). Lateroventral carina present, running from anterior genal region to occipital carina (Fig. 36A). Eye ovoid. Antennomeres II – X about same length, excluding scape and two- segmented club; apical antennomere about two times longer than precedent.

**Mesosoma.** *In dorsal view:* Anterior pronotal margin convex; humerus with subhyaline, triangular lamella, truncate distally, lateral margin straight, forming approximately 60° angle with imaginary intersecting horizontal line (Fig. 36B). Promesonotal suture absent.

Mesonotum with subhyaline, triangular lamella directed anterolateriad. Propodeum with broad, subhyaline lamella, shorter anteriorly, broader posteriorly; lateral margin straight to feebly convex; anterior lamellar margin round to weakly acute, posterior lamellar margin round. *In lateral view:* Propodeal spiracle round, just below propodeal lamella. Meso- and metapleural sutures absent; mesometapleural suture present, without groove. Metapleural gland opening barely discernible.

**Legs.** Meso- and metafemur angulate with or without distal, subhyaline lamella on posteroventral margin. Apices of pro-, meso-, and metatibia with weakly-developed, pointed external lamella. Probasitarsus subcylindrical; meso- and metabasitarsus flattened; probasitarsus about the same length of protarsomeres II – IV; meso- and metabasitarsus longer than respective tarsomeres II – V combined.

**Petiole and postpetiole.** *In lateral view:* Petiole about as high as long; dorsal surface of node relatively flat; subpetiolar process longitudinally ridged, with weakly developed, round tip directed ventrad. Postpetiole longer dorsally than ventrally; dorsal surface of postpetiolar node relatively flat; subpostpetiolar process well-developed, anteriorly lip-shaped. *In dorsal view:* Petiole, excluding lamellae, shorter than broad; anterior petiolar margin slightly concave; petiolar lamella well-developed, subhyaline, rhomboid-shaped, slightly longer than broad, distally subacute. Postpetiole, excluding lamellae, about as long as broad; anterior postpetiolar margin straight; postpetiolar lamella well-developed, subhyaline, longer than broad, distally round; nodal lamella slightly longer than width of node, postnodal lamella slightly shorter than length of postnode. Petiolar node longitudinally shorter than postpetiolar node.

**Gaster.** *In dorsal view:* AIV suboval, slightly longer (GL: 1.31-1.51) than broad (GW: 1.16-1.18); margin of posterior third straight; anteromedian margin u-shaped. Anterior subhyaline lamella present, extending posterad to approximately half length of AIV; anterior lamellar margin round, internal margin continuous with anteromedian margin of AIV. Posterior margin of AIV subacute. *In lateral view*: AIV convex both ventrally and dorsally.

**Color and pilosity.** Body light brown, except whitish to pale yellowish frontal lobes and lamellae; cuticle overall opaque, including legs.

Head, mesosoma and petioles with moderately abundant, appressed, mostly triangular-like, canaliculate hairs. Mandibles with suberect hairs. Canaliculate hairs on AIV relatively less abundant than rest of body and mostly linear or rectangular-shaped.

**Integument and sculpture.** Integument mostly foveate-reticulate, except on AIV and legs. Striae absent from head, mesosoma and petiolar dorsum; shallow striae present on dorsal and lateroventral surface of AIV, and ventrally on head. Propodeal and gastral lamellae bearing faint, horizontal and longitudinal striae, respectively. (Fig. 36B).

**Soldier description.** Measurements (*n*=2): HW: 1.61-1.64; HL: 1.25-1.28; EL: 0.26; CCHA:0.33; PW: 1.58-1.73; WL:1.42-1.47; PTL:0.22-0.24; PTW:0.72-0.73; PPL:0.24-0.26; PPW:0.68-0.7; GL: 1.64-1.69; GW:1.34-1.36; HBL:0.28-0.33; HBW:0.15; **Indices.** CI: 116.28-119.5; OI: 16.43; PI:30.3-32.84; HBI:46.67-53.85; TL: 4.77-4.94.

**Head.** *In frontal view*: Wider than long (CI: 116 – 119), disc-shaped, overall margin raised so that base of disc not visible laterally; short, longitudinal convexity anteromedially on disc floor; anterior margin slightly crenulate; posterior margin with pair of teeth medially; anterior margin with u-shaped cleft medially, lateral margin straight, anterior and posterior margins convex. Occipital corner bearing round, salient carina ventrally. Maximum width about same with of anterior pronotal crest. Masticatory margin of mandible with worn apical tooth, basally edentate. *In lateral view:* Dorsal margin mostly flat. Lateroventral carina present, running from anterior genal region to posteroventral occipital carina. Posterior cephalic margin strongly concave. Eye: as worker. Antenna: as worker.

**Mesosoma.** *In dorsal view:* promesonotum triangular-shaped, maximum width about two times mesopropodeal suture; promesonotal suture absent; anterior pronotal margin convex; pronotal crest acute anteriorly; promesonotal suture deeply impressed; lateral pronotal margin slightly concave, forming approximately 60° angle with imaginary intersecting horizontal line (Fig. 37B). Mesonotum with salient posterolateral, round carina. Propodeum subtriangular, with broad hyaline lamella laterally, anterior lamellar margin round; anterolateral propodeal margin with tiny denticle. Propodeal declivity concave. *In lateral view:* pronotal anterior margin straight; pronotal crest raised; mesonotum straight; propodeal declivity pronounced; posterior propodeal margin continued with dorsal margin, without forming angle. Posterior propodeal carina with vertically raised, round-tipped denticle.

Mesopleural suture absent; mesometapleural suture present, without groove. Propodeal spiracle round, about two spiracle diameters below propodeal carina. Metapleural gland opening barely discernible. *In frontal or dorsal views*: pronotal margin with subhyaline lamella.

**Legs.** As worker.

**Petiole and pospetiole.** As worker.

**Gaster.** As worker, except AIV longer (GL: 1.64 - 1.69) than broad (GW: 1.34 - 1.36).

**Color and pilosity.** As worker, except: head mostly ferruginous to dark brown; dorsum bearing much less appressed canaliculate hairs.

**Integument and sculpture.** As worker, except head dorsum polished.

### Comments

Within the *grandinosus* species-group the workers of *C. esthelae* are medium-sized (WL: 1.01 - 1.16 mm), and considering only the head size, this species is amongst the largest (HL: 0.92 – 1.09 mm; HW: 1.15 – 1.28 mm). In contrast, the average head size of the workers of the other *grandinosus* members is HL: 0.79 – 1.12 mm, and HW: 0.87 – 1.25 mm. The soldiers, however, exhibit a significantly longer head (HL: 1.60 mm) compared to that of the other known members of the *grandinosus* species-group (HL: 1.12 - 1.37 mm). From a morphological perspective *C. persimilis* and *C. persimplex* are the closest species to *C. esthelae,* which differs from them in the following: in frontal view, the head is broader than the pronotum by about one maximum eye length (Fig. 36B), while in both *C. persimilis* and *C. persimplex* the head is almost as broad as the pronotum (<u>CASENT0173700</u>, <u>CASENT0922544</u>); the posterior cephalic margin has a median concavity (Figs. 36C, D), which is absent in *C. persimilis* and *C. persimplex*. *Cephalotes foliaceus* is the only species in the *grandinosus* group with a similarly big head (HL: 1.10 mm; HW: 1.43 mm), but this species is easily distinguished from *C. esthelae* by the lamella of the first gastral tergite, which in *C. esthelae* runs posterad to about the median gastral region (Fig. 36B), whereas in C. *foliaceus* the lamella surrounds the complete gastral margin (<u>CASENT0904925</u>). *Cephalotes esthelae* and *C. persimplex* are the only species in the genus showing triangular-shaped canaliculate hairs on the dorsum of head and mesosoma. These are usually discernible with high magnification (≥ 60 X).

A single nontype worker (Fig. 38), collected near Lumbaqui town (Sucumbíos province) in a well-preserved Amazonian cloud forest, to the eastern Andean chain, shows a longer gaster (GL: 1.51 mm) as compared to the workers of the type material (GL: 1.31 - 1.42 mm) which were collected in lowland rain forests in Esmeraldas province, to the western side of the Andes. In addition, the gastric hairs of this Amazonian specimen are smaller and less abundant (Fig. 38A) as compared to those of the specimens of the type series.

**Figure 38.**
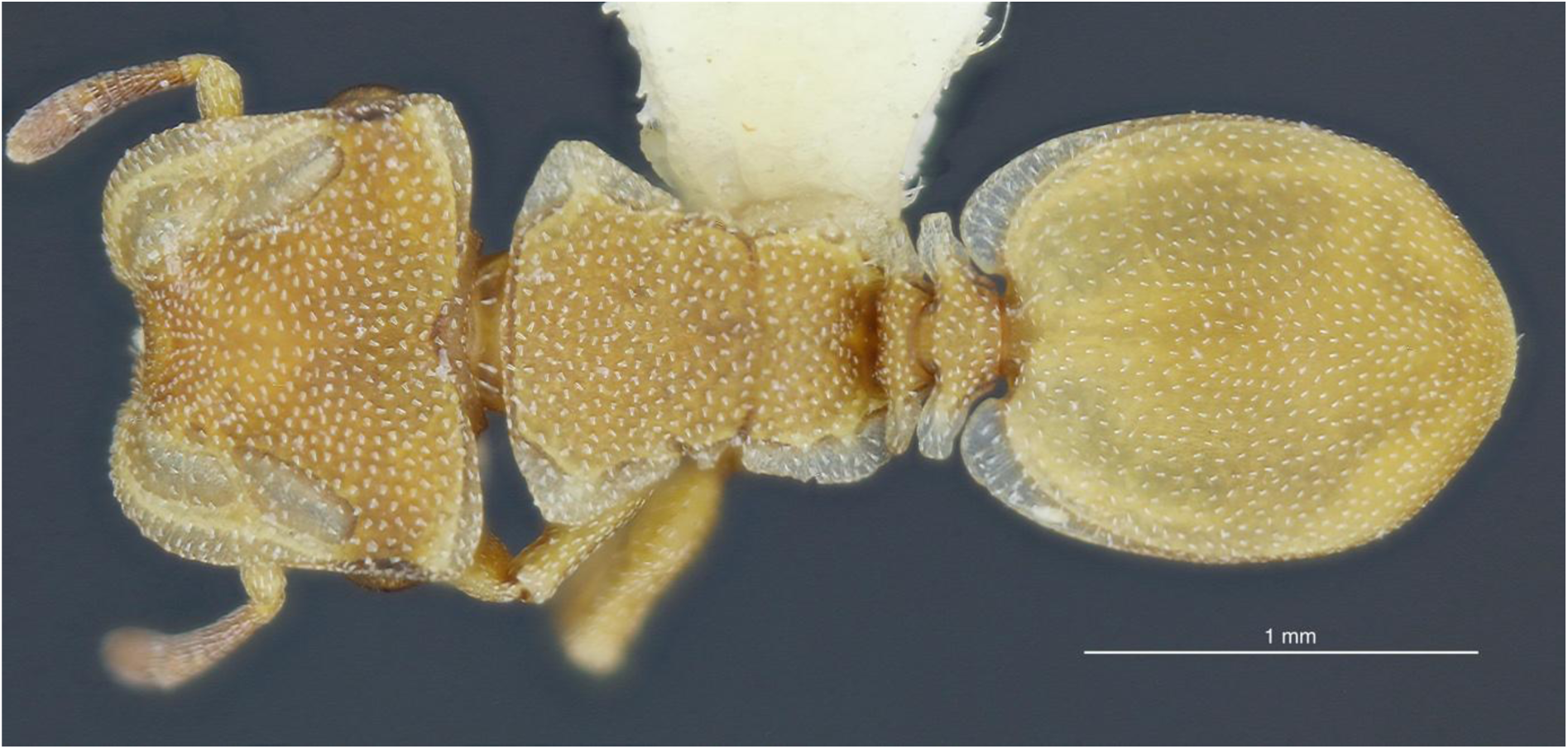
Worker of *Cephalotes esthelae* **sp. nov.** (MEPN: MEPNINV18178) from Lumbaqui town, Amazonia. Note the difference in gaster length and in the gastric hair size, as compared to the holotype. Image by Adrian Troya.

### Distribution notes

*Cephalotes esthelae* is found exclusively in Ecuador, inhabiting well-preserved terra firme forests with an elevation range between 120 m and 550 m. In the northern part of this country, its distribution reaches pristine forests near a small river called Puchuchoa, close to the town of Lumbaqui, in the western side of the province Sucumbíos. This is a transition zone between the lowland rain forests of Amazonia and the Andean foothills, mostly dominated by pre-mountainous cloud forests. To the west, this species is found in lowland well-preserved forests of the Reserva Ecológica Cotacachi Cayapas, province of Esmeraldas, which is part of the southern section of the Chocó Biogeographic. These ecosystems, characterized by their high biodiversity and endemism, are currently under increasing anthropogenic pressure due to high deforestation rates and changes in land use (Kleeman et al., 2022; Rivas et al., 2024), with still unknown, but likely negative impacts on the populations of many species

### Other material examined

☿, ECUADOR, Sucumbíos, Lumbaqui, Quebrada Puchuchoa, 00° 04’51,4”N, 77°16’43,6”W [0.081, -77.2788], 550 m, 23-Ene-2003 (MEPN: ☿, MEPNINV18178).

**Distribution.** Ecuador: Esmeraldas, Sucumbíos *Cephalotes sacha* Yumbay & Troya sp. nov. Zoobank:

Figs 39: A-F (☿ holotype); 40: A-C (☿ soldier); Fig 43B (distribution of records)

**Figure 39.**
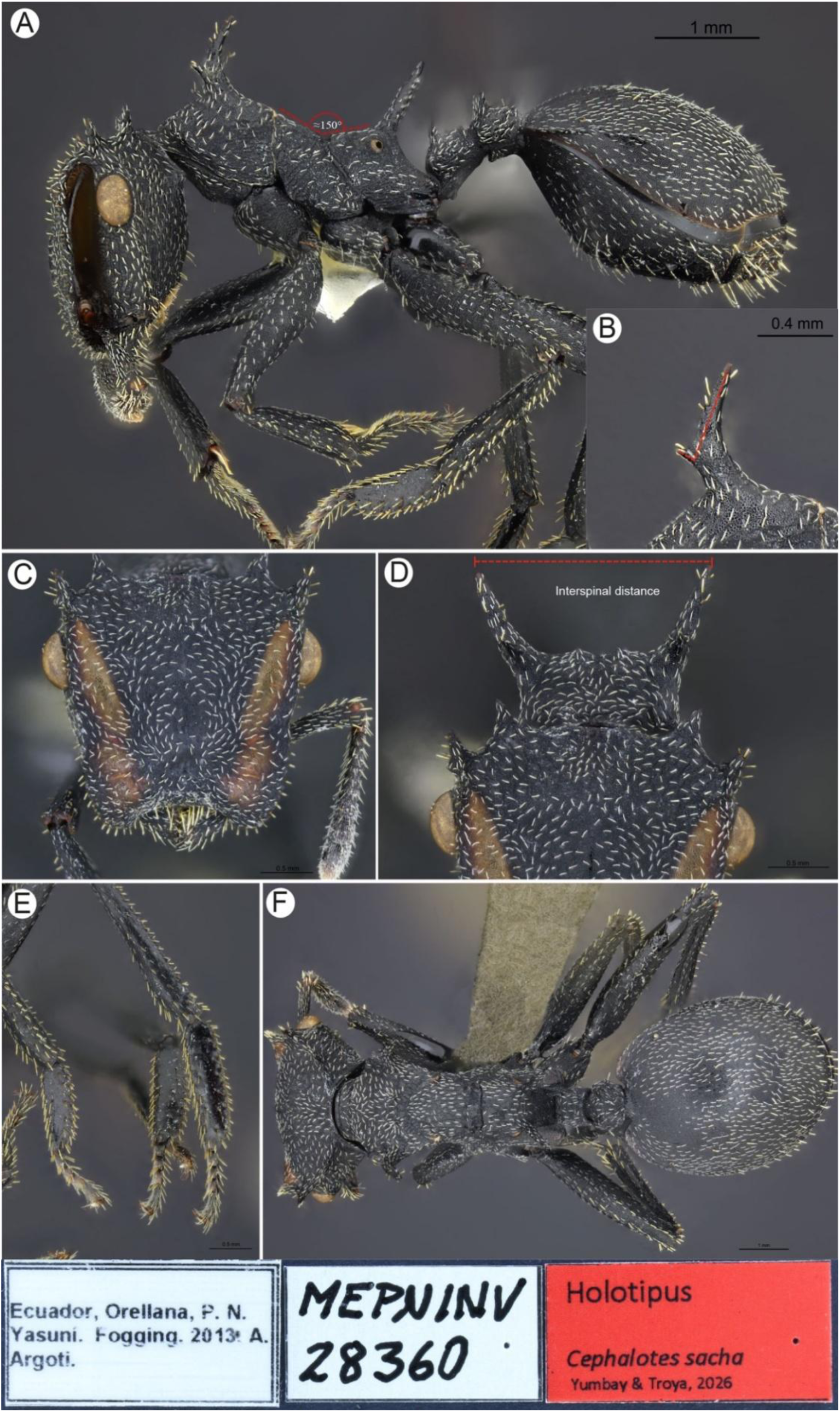
Holotype worker of *Cephalotes sacha* **sp. nov.** (MEPN: MEPNINV28360) **A.** Full body, lateral view. **B.** Pronotal spine, lateral view. **C.** Head, full-face view. **D.** Close-up of pronotal spines, frontal view. **E.** Meso-and metatarsi, lateral view. **F.** Full body, dorsal view. Collection labels are shown at the image’s foot. Image by Akayky Yumbay.

**Type material.** Holotype. ☿; ECUADOR: Orellana: P. N. [Parque Nacional] Yasuní. Fogging [method]. 2013. A. Argoti [leg.]. (MEPN: MEPNINV28360).

**Paratypes**: 12 ☿. Same data as holotype (MEPN: ☿ soldier, MEPNINV28356; ICN: ☿ soldier MEPNINV28448; DZUP: ☿, MEPNINV28361; IAVH: ☿, MEPNINV28363; ICN: ☿, MEPNINV28364; QCAZ: ☿, MEPNINV28358; MCZC: ☿, MEPNINV28359; USNM: ☿, MEPNINV28362); same data as holotype, except: Aguarico Cononaco, Guacamayo Saladero, 01° 17×40’’ S 76° 03’ 39’’W [-1.2944, -76.0608], 215 m Fogging, 02-feb-14 A. Troya & P.

Duque (MEPN: ☿ soldier, MEPNINV35175; ☿, MEPNINV 35174); ECUADOR: Sucumbíos: Lumbaqui, Fogging, ene- 2003, P. Araujo (MEPN: ☿, MEPNINV37758; MCZC: ☿, MEPNINV37759).

**Etymology.** The specific epithet is derived from the Kichwa “sacha”, meaning “forest” or “jungle”. The name alludes to the Amazonian rainforests where specimens of this species were found. This is a tribute to both the forests and the indigenous language spoken by numerous peoples and communities in Ecuador. As such, we highlight the need to preserve our natural and cultural heritage. The epithet is a noun in apposition, thus indeclinable (International Code of Zoological Nomenclature: Arts.11.9.1.2.; 31.2.1).

**Diagnosis.** A member of the *atratus* species-group. Workers: Mesonotum and propodeum forming a ∼ 150° angle, in lateral view (Fig. 38A); integument opaque; in lateral view, postpetiolar dorsal spines longer than subpostpetiolar process; body with simple hairs without foveae (Figs. 39A, C, D); in dorsal view, promesonotum without Y-shaped carina (Fig. 39F); anterior projection of pronotal spine about four times shorter than posterior projection, forming straight angle with it (Fig. 39B). Soldiers: Pronotum with pair of blunt teeth dorsally, and pair of salient, horizontally directed, angulate projections with truncate tips laterally (Fig. 40A); cephalic disc subquadrate with acute projection on occipital corner (Fig. 40B); anterolateral margin of gaster bearing slightly salient, non-translucent lamella which fades away in the first anterior fourth dorsally (Fig. 40C).

**Figure 40.**
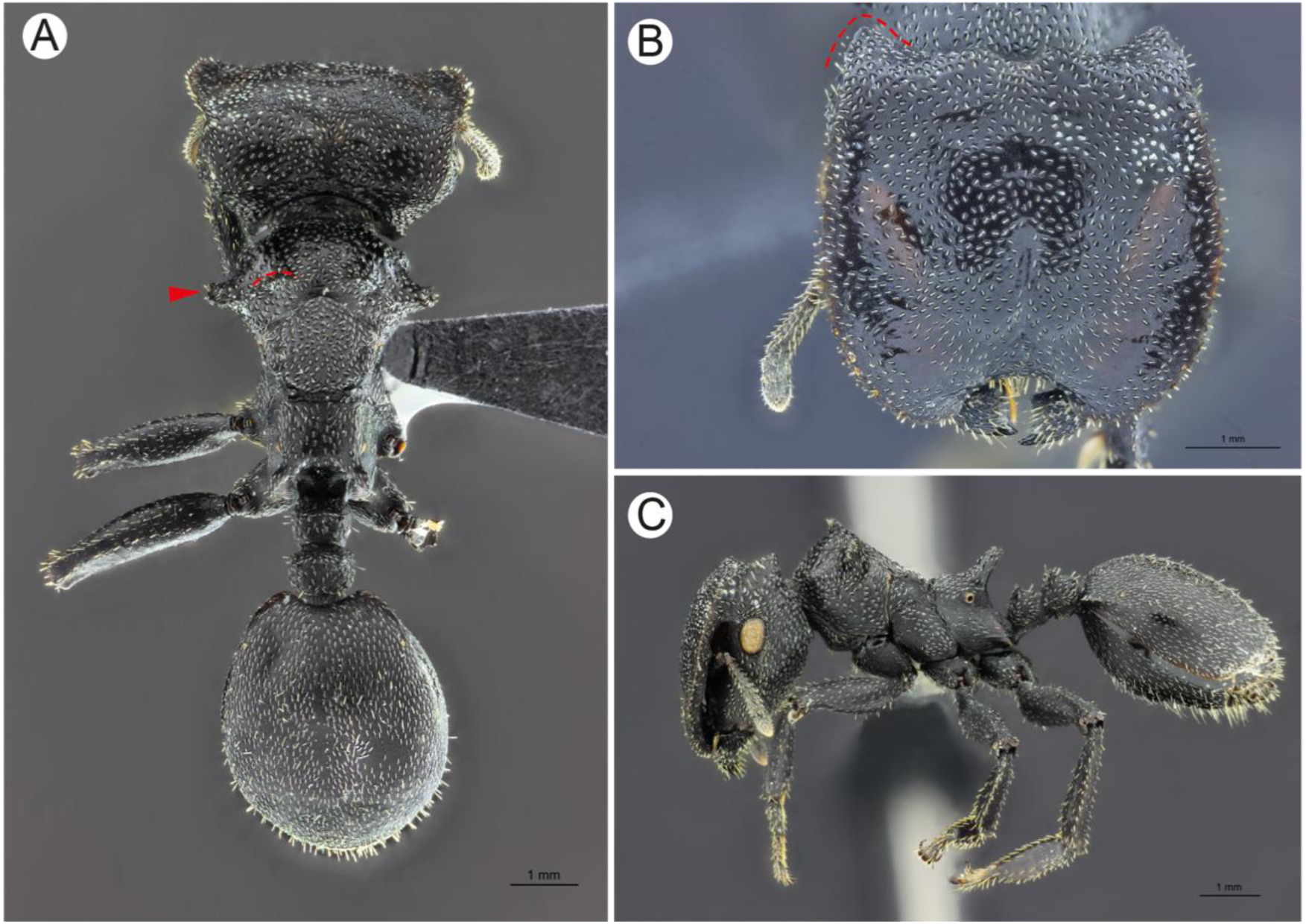
Soldier of *Cephalotes sacha* **sp. nov.** (MEPN: MEPNINV28356) **A.** Full body, dorsal view. **B.** Head, full-face view. **C.** Full body, lateral view. Red arrowhead: angulate projection with truncate tip; Red dashed line: pronotal blunt tooth. Image by Akayky Yumbay.

**Worker description.** Measurements (*n=*10): HW: 2.13-2.62 (2.34); HL: 2.1-2.53 (2.34); EL: 0.4-0.5 (0.43); PW: 1.5-1.79 (1.63); WL: 2.2-2.62 (2.49); PTL:0.44-0.59 (0.53); PTW:0.32-0.55 (0.54); PPL:0.44-0.55 (0.48); PPW:0.58-0.67 (0.64); GL: 2.3-2.88 (2.56); GW:2.1-2.27 (2.17); HBL:1.28-1.66 (1.44); HBW:0.29-0.42 (0.35); ID:1.43-1.82 (1.78); **Indices.** CI: 88.46-103.53 (100); OI: 18.19-21.51 (18.19); PI:93.75-137.5 (97.6); HBI:22.5-27.8 (24.44); TL:7.51-9.21 (8.37).

**Head.** *In frontal view:* Subquadrate, slightly longer than wide (CI: 88–103); occipital corner with pair of equally sized spines, each about ¾ maximum eye length; posterior margin feebly convex medially; lateral margin of frontal lobe mostly straight, weakly crenulate, slightly concave medially; anterior margin of frontal lobe weakly crenulate, round. Eye completely visible. Anterior clypeal margin broadly concave, at least ½ of mandible dorsum visible.

External mandibular margin convex, masticatory margin with two apical teeth, followed by one to three denticles distally. *In lateral view:* Slightly convex medially; lateroventral carina present, running from gena to posteroventral ocular margin. Postgena convex. Nuchal carina well-developed, slightly pointed anteriorly. Eye ovoid. Antennomeres II – VIII about same length, each smaller than apical antennomeres forming a three-segmented club; apical antennomere about ¼ longer than precedent.

**Mesosoma.** *In dorsal view:* Anterior pronotal margin slightly convex, humerus with bifurcate spine, feebly inclined laterad. Anteroventral pronotal carina salient, directed anterolaterally. Maximum pronotal width about 1.5 times broader than width of mesonotum. Promesonotum trapezoid, with angular lateral margins. Promesonotal suture absent. Mesopropodeal groove strongly impressed. *In lateral view*: Pronotum convex. Propleuron convex, slightly angulate. Mesopleural suture absent. Mesometapleural groove impressed, vanishing before reaching ventral cleft. Propodeum with long dorsolateral spine, about six times as long as basal width, inclined posterad, making about 150° angle with propodeal dorsal margin; propodeal declivity straight, inclined anterad, making about 90° angle with dorsum of straight propodeum.

Propodeal spiracle circular, located about 1.5 – 2 spiracular diameters from propodeal dorsal margin. Orifice of metapleural gland opening slit-shaped, horizontally oriented, barely discernible, bounded distally by horizontal carina.

**Legs.** Pro-, meso-, and metafemur broadened basally, distally narrowed; protibia angulate, slightly flattened laterally; meso- and metatibia angulate, flattened laterally; meso- and metabasitarsus strongly flattened laterally, longer than respective tarsomeres II – V combined. Tarsal claws bearing small preapical tooth.

**Petiole and postpetiole.** *In lateral view:* Petiolar node, excluding pair of dorsal spines, slightly longer than high; nodal spine well-developed, slightly longer than basal width; dorsal nodal surface convex; subpetiolar process well developed, longitudinally ridged, with blunt, anteroventral tip. Postpetiolar node about as long as high; postnodal spine well-developed, slightly longer than nodal spine; dorsal postnodal surface slightly convex; subpostpetiolar process weakly developed, lip-shaped. *In dorsal view*: Petiole as long as broad, anterior margin feebly convex, posterior margin feebly concave; lateral margins feebly convex. Postpetiole slightly broader (PPW: 0.57 – 0.67) than long (PPW: 0.44 – 0.55), anterior margin straight, posterior margin feebly convex; lateral margins convex.

**Gaster.** *In dorsal view:* AIV oval, longer (GL: 2.30 - 2.88) than broad (GW: 2.01 - 2.27). Anterior lamella small but well-developed, extending posterolaterad to about one sixth length of AIV where it fades away; anterior lamellar margin rounded, internal margin continuous with anteromedian margin of AIV; posterior margin of AIV rounded. *In lateral view*: AIV dorsally convex; ventrally strongly convex.

**Color and pilosity.** Body opaque black, head dorsum bearing brownish to pale yellowish longitudinal stripe running from anterior margin of frontal lobe to posterolateral head margin. Dorsal and ventral surface of head, mesosoma, petioles and anterior dorsal region of AIV with moderately abundant, white to golden, mostly appressed hairs; antennae, mandibles, tibiae, tarsi and gaster bearing few erect and suberect hairs.

**Integument and sculpture.** Integument rugulose, composed of puncta and irregular, linear, non-continued striae located posterodorsally, laterally, and ventrally on head, dorsally on mandibles, most of mesosoma, except ventrally, legs, dorsally on petioles, and anterodorsally on AIV.

**Soldier description.** Measurements (*n=*3): HW: 3.4-4.1; HL: 2.88-3.7; EL: 0.55-0.63; PW: 2.27-3.1; WL:3.4-4; PTL:0.64-0.77; PTW:0.58-0.83; PPL:0.58-0.67; PPW:0.74-0.93; GL: 3.35-3.8; GW:2.59-3.7; HBL:1.6-1.64; HBW:0.45-0.54; ID:2.3-3; **Indices.** CI: 90.24-97.33; OI: 16.22-19.1; PI:84.62-111.11; HBI:28-34; TL:10.65-12.99.

**Head.** *In frontal view*: Subquadrate, about as long as broad (CI: 90–97); anterolateral margin convex, crenulate; anterior margin strongly concave, smooth; posterior margin straight, medially bearing pair of blunt teeth; occipital corner angulate, pointed posterad. Frontal lobe completely covering malar area and gena, and almost completely covering eye. External mandibular margin convex, masticatory margin with one apical tooth, followed by one to three tiny denticles distally. *In lateral view:* Slightly convex dorsally and ventrally; lateroventral carina absent; nuchal carina reduced*. In frontal or dorsal view:* Pronotum shorter than maximum head width. Eye: as worker. Antenna: as worker.

**Mesosoma.** *In dorsal view:* Promesonotum triangular-shaped; anterior propodeal margin convex, humerus bearing stout, salient, truncate lobe; promesonotal suture shallowly impressed. Mesonotum pentagonal; mesopropodeal suture deeply impressed. Propodeum subquadrate, bearing stout, irregular, laterally diverging spine. *In lateral view:* Pronotal top acute, straight anterior margin making 90° angle with straight promesonotal margin.

Promesonotum making about 120° angle with propodeal dorsum. Propodeal spine inclined posterad, about three times as long as basal width; posterior propodeal margin inclined anterad. Propodeal spiracle round, about two spiracle diameters from propodeal dorsum.

Orifice of metapleural gland opening slit-shaped, horizontally oriented, barely discernible.

**Legs.** As worker.

**Petiole and postpetiole.** As worker, except: node about as long as high, in lateral view; nodal spines shorter, blunt-tipped; anterior tip of subpetiolar process spiny-shaped, in lateral view.

**Gaster.** As worker.

**Color and pilosity.** As worker, except: dorsal head stripes pale ferruginous, barely discernible.

**Integument and sculpture.** As worker, except: puncta smaller, and mostly on mesopleuron, propodeum, petiolar and postpetiolar dorsum, legs, and gaster. Impressed foveae on head, mesosoma, petiole and postpetiole.

### Comments

Among species in the *atratus* species-group, *C. sacha* is relatively small (WL: 2.20 – 2.62), and except for *C. serraticeps* and *C. alfaroi*, its morphologically closest taxa, *C. sacha* is very different from the other group members: *C. atratus*, *C. placidus*, *C. oculatus,* and *C. opacus*. In regards to *C. serraticeps, C. sacha* differs in the following characters: in dorsal view, the promesonotum of the worker lacks a Y-shaped carina (present in *C. serraticeps*) (Fig. 11A); the anterior projection of the pronotal spine is almost six times shorter than the spine’s length, (in *C. serraticeps* it is about three times shorter than the spine’s length); in lateral view, the mesonotum and propodeum make an obtuse angle of about 150° (Fig. 39A) (in *C. serraticeps* is about 180° or more, sometimes no angle is noticed) (see figure 11C in Pazmiño-Palomino and Troya, 2022); in frontal view, the interspinal distance (Fig. 39D) is narrower than the corresponding in *C. serraticeps*. In the soldiers, the occipital corner has a single acute carina (in *C. serraticeps* it has three distinct pointed teeth); the meso- and metapleural plates are foveate (pleural foveae are absent in *C. serraticeps*).

As of *Cephalotes alfaroi* we examined the image of the holotype worker (<u>CASENT0904899</u>) in AntWeb, and distinguished the following differences with respect to *C. sacha*: well-marked pleural striae (in *C. sacha* the pleural striae are shallower, less marked than in *C. alfaroi*); dorsum of gaster smooth and glossy, and in general, body relatively shiny (in *C. sacha* the gastral dorsum is opaque and striate, as well as the rest of the body); the propodeal spiracle is about three spiracular diameters off the propodeal dorsum (about 1.5 – 2 spiracular diameters in *C. sacha*). Regarding a non-type *C. alfaroi* worker from Corcovado National Park, at the pacific southwestern coastal region of Costa Rica (<u>CASENT0040145</u>), identified by Brian Fisher in AntWeb, we noticed the gastral dorsum has slight longitudinal striae anteriorly, but the rest of the integument is smooth and glossy, as in the holotype, which is from Jimenez, Limón province, in the northeastern Atlantic side of that country.

As to the soldier of *C. alfaroi*, its posterior vertexal border (near the occipital foramen) has a pair of well-developed teeth (absent in *C. sacha*); the propodeal spines of *C. alfaroi* are longer than those of *C. sacha*; the petiole of *C. alfaroi* is devoid of spines or teeth dorsally (in *C. sacha* the petiole has two well-developed, but short spines).

**Distribution notes.** *Cephalotes sacha* is only known from Ecuador, it inhabits well-preserved Amazonian rainforests, from approximately 190 m up to 600 m of elevation. To the northern range it reaches the forests of the Aguarico river, near Lumbaqui, a well-preserved area in western Sucumbíos province, close to the southern Colombian border. These forests are located in a transition zone between the Amazonian lowlands and the Andean foothills, which are currently subject to strong anthropic pressure (Kleemann et al., 2022). To the south it reaches the southern region of Parque Nacional Yasuní, near the Cononaco river, just at the border of Orellana and Pastaza provinces, and close to the western Peruvian border. This area is mostly composed by pristine, lowland rainforests showing a high annual rainfall (Bass et al., 2010). This latter region of Parque Nacional Yasuní is arguably the most well-preserved site of the whole Ecuadorian Amazonia.

### Natural history notes

The two specimens from Cononaco river were collected from a Myrtaceae tree, close to a stream. In winter, this collection site is probably flooded as judged by the water marks on the base of the Myrtaceae tree trunk.

**Distribution.** Ecuador: Orellana, Sucumbíos.

### New caste description

*Cephalotes dentidorsum* De Andrade, 1999 Soldier (Fig. 41)

**Figure 41.**
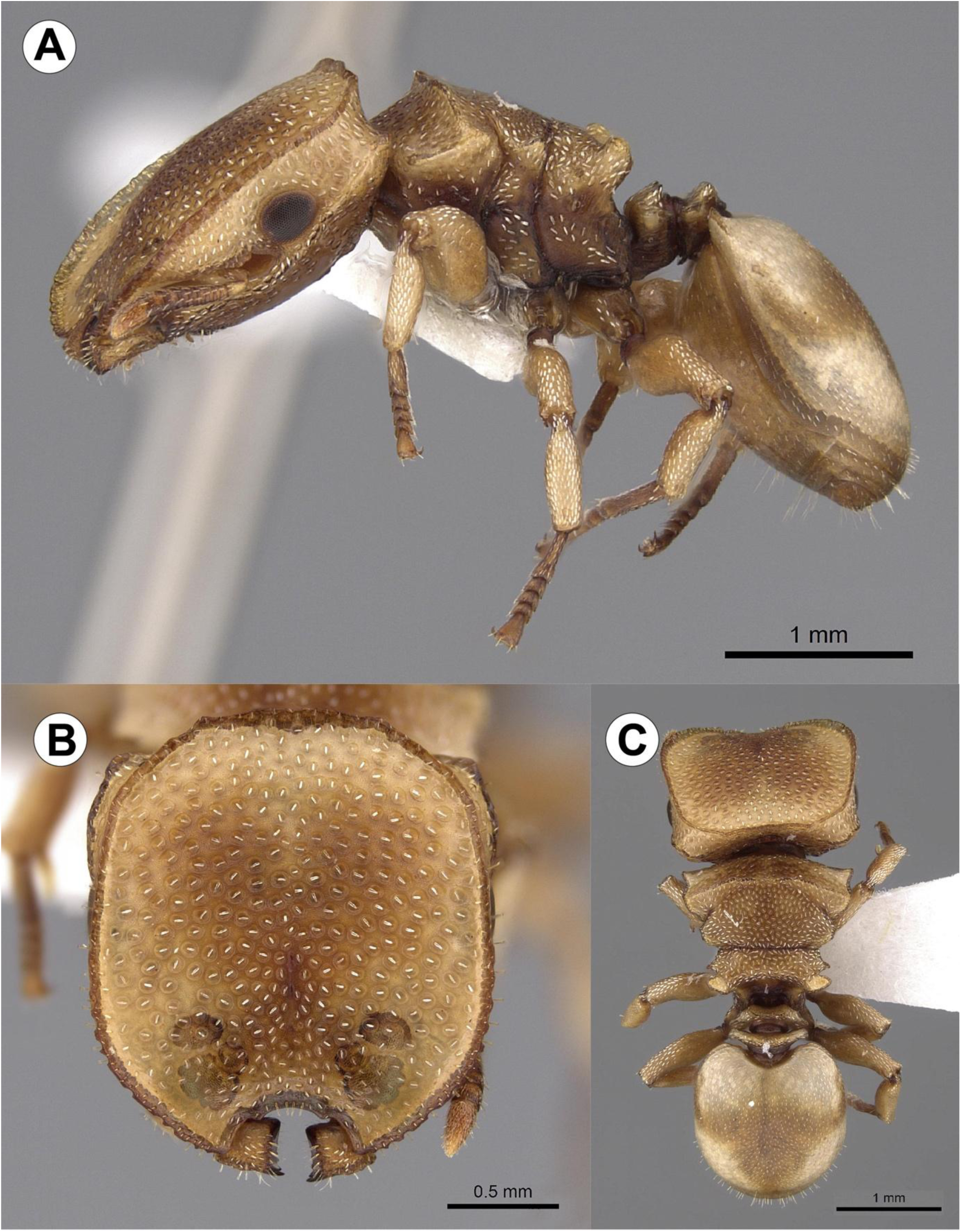
Soldier of *Cephalotes dentidorsum* (PSWC: CASENT0922531). **A.** Full body, lateral view. **B.** Head, full-face view. **C.** Full body, dorsal view. Image by Wade Lee.

**Figure 42.**
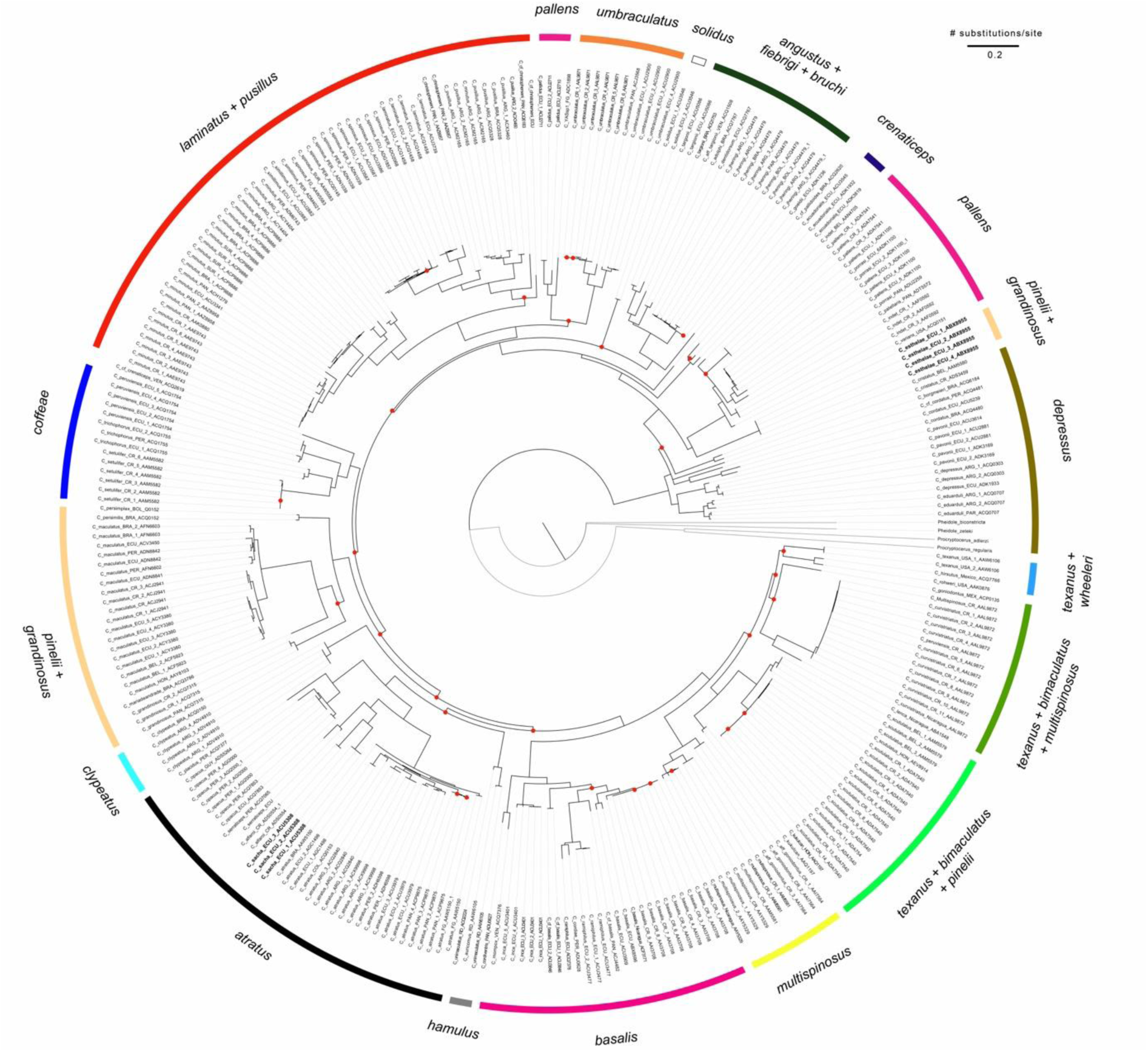
Maximum likelihood CO1 phylogeny of the genus *Cephalotes*. Species-groups are shown on the outer margin. Gray branches are outgroups. Newly described species are bolded. Nodes with < 70% support are shown with red dots. The tree tips contain: species name, country abbreviated, BIN (Bardode Index Number).

**Figure 43.**
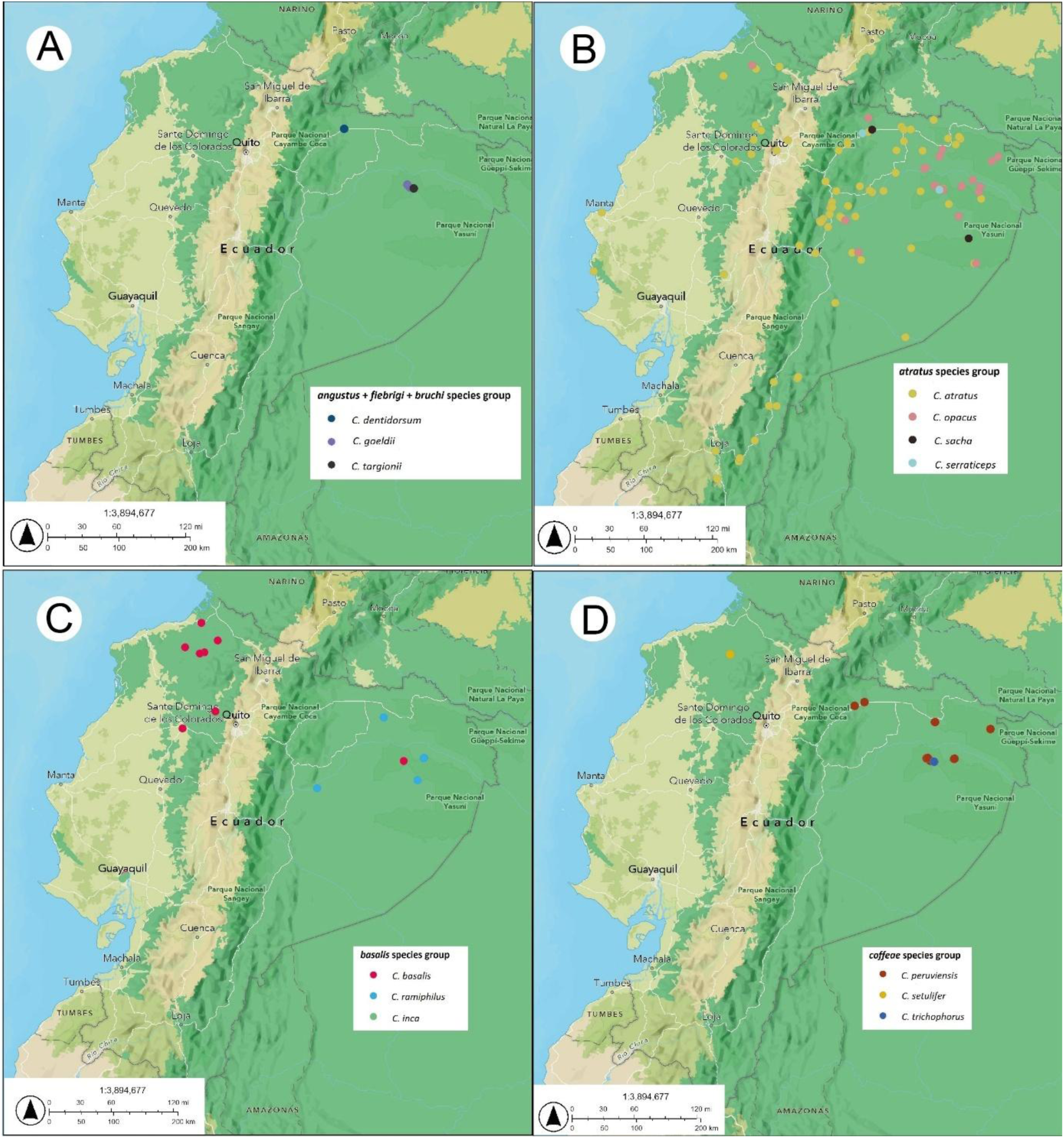
Distribution of examined species records in each species-group. (A) *angustus + fiebrigi + bruchi*. (B) *atratus*. (C) *basalis*. (D) *coffeae*.

**Figure 44.**
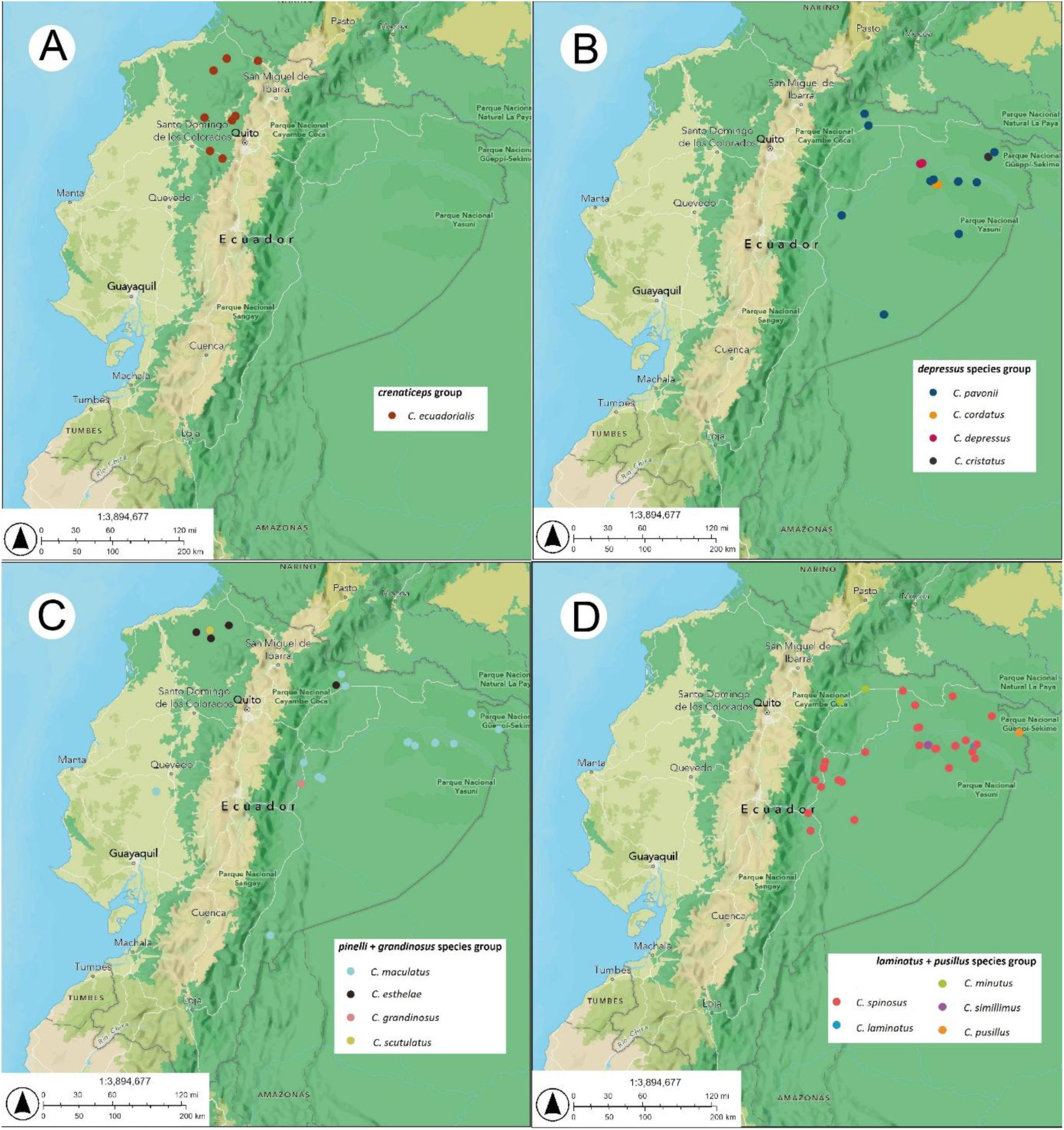
Distribution of examined species records in each species-group. (A) *crenaticeps*. (B) *depressus*. (C) *pinelii + gandinosus*. (D) *laminatus + pusillus*.

**Figure 45.**
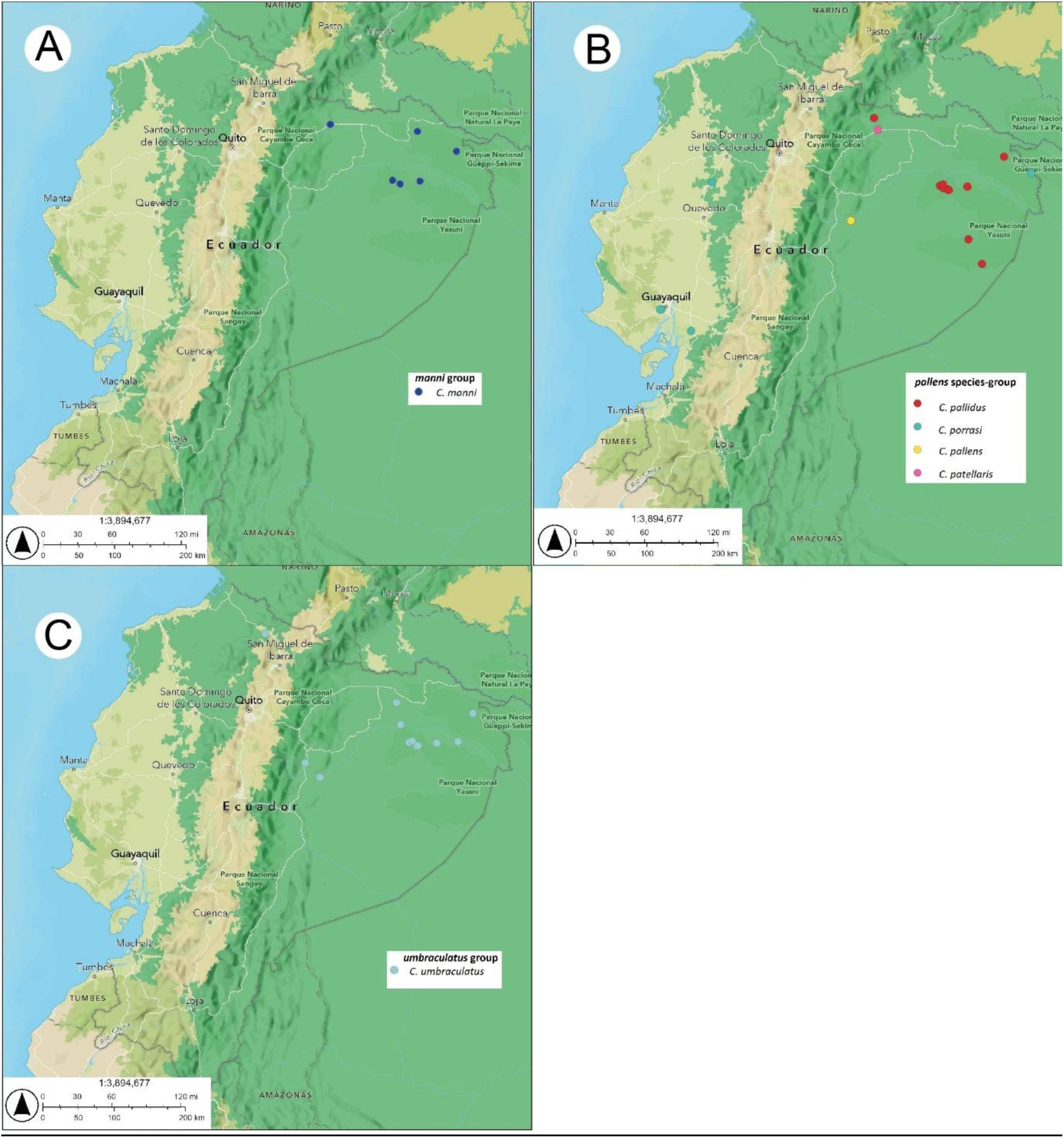
Distribution of examined species records in each species-group. (A) *manni*. (B) *pallens*. (C) *umbraculatus*.

**Material examined.** Ecuador. Orellana. Parque Nacional Yasuní, -0.66667, -76.3833, 120 m, 1☿ (soldier), 2005-07-05, Argoti, A., fogging, (MEPN: MEPNINV28455); Peru. San Martín. Convento, 26 km NNE Tarapoto, -6.266667, -76.3, 220 m, 1☿ (soldier), 1986-08-20, Ward, P. S., manual, edge of second-growth rainforest, ex *Tachigalia* Benth. & Hook.f. (PSWC: CASENT0922531).

**Head**. *In full-face view*: subquadrate slightly wider than long (CI 102-102.2). Mandible with four teeth, strongly angled laterally. Clypeus with pair of denticles weakly defined. Dorsum of head disc-shaped, slightly convex anteriorly. Frontal carina crenulate anteriorly. Antenna with three-segmented club. Dorsal margin of antennal scrobe with lateral carina and posterior rounded projection. Lateroventral head margin without carina. Posteroventral cephalic corner acute, separated from dorsal cephalic disc.

**Mesosoma**. *In lateral view:* pronotum ascending, with a transversal carina raised in a crest; pronotal crest not crenulate, with a median depression. In dorsal view, anterior margin of pronotum weakly convex, lateral margins with two pairs of denticles, the anterior one acute, the second one rounded at level of transversal carina of pronotum. Mesonotum and propodeum discontinuous and flat; mesonotum with a pair of blunt rounded denticles; propodeal groove strongly impressed; dorsal and declivous faces of propodeum meeting in a distinct propodeal angle. In dorsal view, lateral margins of propodeum with a pair of median spines and obtuse denticles strongly bent dorsally, with the apices curved anteriad. Femora not angulated dorsally, mid, and hind basitarsi not flattened, with subparallel dorsal and ventral faces.

**Legs.** As worker.

**Petiole and postpetiole.** As worker, except: In dorsal view, petiole compressed anteroposteriorly, anterior margin with a distinct median concavity, lateral spines curved posterad, dorsum with a pair of denticles, subpetiolar process broader and rounded anteriorly. Postpetiole slightly wider and longer than petiole, spines narrow and curved forward, dorsum with a “V” sharped elevation, subpostpetiolar process pronounced ventrally and compressed anteroposteriorly.

**Gaster.** As worker. Elongate, with narrow anterior lamellar expansions.

**Color and pilosity.** As worker, except: Head, dorsum of mesosoma and legs yellowish to ferruginous. Propleura predominantly ferruginous with the dorsal surface yellowish; meso-and metapleura predominantly yellowish with ferruginous spots. AIV tergite yellowish with a transverse dark macula. AIV sternite yellow-brownish. Head, mandibles, mesosoma, petiole, postpetiole and legs with appressed canaliculate hairs; frontal carinae with erect simple hairs. Gaster with appressed simple hairs; posterior portion of first sternite and edges of each tergite and sternite of gaster with erect simple hairs.

**Integument and sculpture.** As worker, except: Mandibles with reticulate sculpture; dorsum of head irregularly foveate. Dorsum of mesosoma foveate; lateral of mesosoma finely rugose with sparse foveae. Petiole and postpetiole with longitudinal striae. Gaster microalveolate.

Measurements (*n*=2): HW: 1.82-1.98; HL: 1.78-1.94; EL: 0.24-0.33; PW: 1.63-1.76; WL :1.32-1.62; PTL: 0.2-0.22; PTW: 0.75-0.76; PPL: 0.3-0.32; PPW: 0.76-0.79; GL: 1.3-1.41; GW: 1.45; HBL: 0.90; HBW: 0.35; Indices. CI: 102-102.2; OI: 18.1-12.1; PI: 26.3-29.3; HBI: 125; TL: 4.9-5.51.

### Comments

This rarely collected species is phylogenetically placed in the *C. angustus* group sensu De Andrade and Baroni Urbani (1999), confirmed in Oliveira et al. (2021), but sensu Price et al. (2022) is placed in the *angustus* + *fiebrigi* + *bruchi* group. Workers and soldiers of *C. dentidorsum* are highly similar to those of the also rare *C. adolphi,* known only for mid-eastern Brazil, mainly from the Cerrado (Oliveira et al., 2021) Workers differ only in the sculpture of the first gastral segment being opaque and with slight punctation in *C. adolphi*, and smooth and shining in *C. dentidorsum*. However, soldiers apparently differ in their cephalic index (longer head in *C. adolphi*, slightly wider in *C. dentidorsum*) and anapleural groove (well-impressed in *C. adolphi*, weakly impressed in *C. dentidorsum*). Despite these characteristics, it is important to note that it remains to be clarified whether they are indeed different species and not populational variants. As suggested by the results of our CO1 phylogeny (Fig. 42) these two are putative sister species, differing in less 0.1 substitutions per site (< 2% genetic divergence). Pazmiño-Palomino & Troya (2022) first recorded this species for Ecuador based on three fogging-collected Amazonian workers.

## Discussion

### Putative signals of divergence in a C. esthelae population and placement of the new species in their respective species-groups

Molecular data usually enables detection of genetic divergence that may not be apparent from morphology alone, and this is particularly relevant for species that exhibit certain phenotypic plasticity or have evolved in isolation (Vrijenhoek, 2009). From our phylogeny, unlike *C. sacha*, which forms a well-supported group with its similar species *C. alfaroi* and *C. serraticeps* (Fig. 2A), *C. esthelae* is genetically quite divergent, > 17%, from its known morphologically closest species, *C. persimilis*, *C. persimplex*, and *C. grandinosus* (Fig. 3B). Except for a few characters in the worker caste, notably the bigger head of *C. esthelae* as compared to the other three species, they are very similar among each other, which was also noted by De Andrade & Baroni-Urbani (1999).

While the placement of *C. sacha* in the *atratus* group was certainly expected because it shares several characters with *C. alfaroi* and *C. serraticeps*, as well as with *C. atratus*, the placement of *C. esthelae* within the *Cephalotes* tree, closer to species in the *pallens* group (Fig. 42), was not. A closer examination of the four sequences of *C. esthelae* in the alignment (Supplementary file 3) shows various non-homology sites as compared to its known morphologically closest species. The resulting genetic distance may be explained by the long geographic distance, more than a thousand kilometers, separating the Chocoan populations of *C. esthelae* from those of *C. persimilis* from Minas Gerais (Cerrado and Atlantic Forest biomes, southeastern Brazil), and from the Amazonian *C. persimplex*. Both the Andes as dispersal barrier, as well as differing climatic, ecological and physiognomic conditions of the habitats where these species live, are potential explanatory variables. However, with respect to the Panamanian and Costa Rican *C. grandinosus*, geographic distance may not be sufficient to explain genetic divergence because the populations of *C. esthelae* are not as separated from those of Panama and Costa Rica, than they are from the Amazonian *Cephalotes* species. In addition, these species inhabit in the same biome. Other factors like structure of the habitats and microhabitats (Abad-Franch et al., 2021), changes in feeding strategies (Guevara-Andino et al., 2025), unidentified geographic-ecological barriers, and even non-optimal field and museum preserving conditions (Ruppert et al., 2023) of the *C. esthelae* specimens before sequencing the material, could hold cues to explain such result.

*Cephalotes esthelae and C. sacha are* currently endemics for Ecuador, and further sampling will likely unveil additional site records in both Chocó and in Amazonia. As to *C. esthelae*, we here consider both populations, from Chocó and Amazonia, as belonging to a single species. However, the Amazonian population, represented by a unique individual (Fig. 38, MEPNINV18178), differs from those of Chocó in the shape of the gaster, which is longer in such Amazonian specimen (see comments under *C. esthelae* description), and its gastric pilosity is smaller and less abundant as compared to that of the Chocoan populations. These differences in morphology could represent signatures of lineage divergence, which may already be occurring due to a decline or complete breakdown of communication between the Chocoan and Amazonian populations that are separated by the Andean chain. We do not have DNA data of the Amazonian specimen to support this assumption, however, the Andes as a barrier restricting gene flow (Salgado-Roa et al., 2024; Salazar & Villalobos, 2024), and the distance separating these populations, approximately 200 km, are factors to consider in future analyses which will support (or refute) our hypothesis.

Evidence of speciation events promoted by vicariance or dispersion in the context of the Andes uplift in northwestern South America has been demonstrated before with birds (Hazzi et al., 2018), mammals (Vallejos-Garrido et al., 2023), frogs (Mendoza et al., 2015), among other groups, but also with ants, for example, *Atta cephalotes* (Muñoz-Valencia et al., 2022).

Because of the variation in morphology of the Amazonian population of *C. esthelae* as compared to the populations of Chocó, we excluded this specimen from the type material. This responds to the need to avoid taxonomic ambiguities by prioritizing stable diagnostic characters over geographic variants, a criterion applied in groups with morphological plasticity (Bickford et al., 2007).

### Additions to current knowledge in the distribution of other Cephalotes species

Taxonomic contributions in insects are especially relevant in tropical, highly diverse regions where usually a tiny fraction of the true diversity is known. Our proposal of *C. esthelae* and *C. sacha* not only expands our knowledge of the diversity in the genus but also highlights the need to revise scientific collections and their associated information in search of potential errors and/or unverified records. While examining collection records in Antweb we noted four records of *C. alfaroi* originating in Brazil (Roraima and Amazonas). To our current knowledge, that species occurs only in Costa Rica and Panama (Basset et al., 2012, Antweb.org). Besides locality, coordinates and collection date, no other information is provided in Antweb, more importantly the four records are not validated by a taxonomist. Oliveira et al. (2021) did not mention this species in their *Cephalotes* revision for Brazil. The lead author examined ant collections from Colombia, Ecuador, Peru, and Brazil, but did not find records either. We suggest the Brazilian records are misidentifications, and those specimens could be either *C. sacha* or *C. serraticeps*.

### Implications for the conservation of the habitats home of C. esthelae and C. sacha

The regions where the new species originate from are highly anthropized and threatened (A. Troya, personal observation), thus both species may be considered potentially vulnerable. In the Chocó region, mining concessions have caused severe deforestation and fragmentation, while in the Amazonian Lumbaqui town, agriculture expansion and urbanization continue to degrade natural habitats (Morales Corozo 2022). These impacts threaten the animals that depend on tree canopies for nesting and dispersal (Gindhart et al., 2024). On the other hand, although the populations of *C. sacha* live in the overall well-preserved Parque Nacional Yasuní, it faces significant threats due to extractive activities, particularly oil exploitation, which compromise both ecosystem integrity and local indigenous communities (Bass et al., 2010). Deforestation remains a leading cause of biodiversity loss in Neotropical insects, disrupting ecological functions and reducing habitat availability (Sánchez-Bayo & Wyckhuys, 2019).

## Conclusion

The exploration and documentation of new species is essential to filling gaps in taxonomic-biodiversity knowledge. The present new species hypotheses add to such aim. This study enhances our understanding of *Cephalotes* diversity in Ecuador and highlights the need to conserve endangered habitats and increase field research in understudied regions. To address these challenges, it is crucial to implement conservation strategies that promote habitat restoration, involving local communities in the protection and sustainable management of both degraded and well-preserved ecosystems. Further research will be required to confirm the presence of the current two new species in other Neotropical regions, thus improving our understanding of their distribution and biogeography.

Distribution of examined records

## ACKNOWLEDGEMENTS

We deeply thank Fabián Bersosa and Pablo Araujo who designed and led the project “Diversidad de Coleoptera de dosel de la Reserva Ecológica Cotacachi Cayapas,” funded by Fundación EcoCiencia Ecuador. The fieldwork of that project took place in several sites of the northern Ecuadorian Chocó where one of the new species was discovered. We are also grateful to such project’s research team: Paulina Rosero, Ana María Ortega, Paulo Guerra, and Nelson Miranda, for their contribution in field and lab work by classifying tens of thousands of specimens, many of which are likely undescribed species. We thank Adriana Argoti for collecting and depositing the other new species in MEPN. AT further acknowledges the many dedicated volunteers who, over the years, assisted with sorting ant material at MEPN. Without their efforts, numerous specimens would still remain unnoticed in alcohol-preserved collections. Finally, we appreciate the valuable feedback of the peers [insert here names] whose comments enhanced the quality of our manuscript.

## Notes

### Competing Interest Statement

The authors have declared no competing interest.

